# A transposase-derived gene required for human brain development

**DOI:** 10.1101/2023.04.28.538770

**Authors:** Luz Jubierre Zapater, Sara A. Lewis, Rodrigo Lopez Gutierrez, Makiko Yamada, Elias Rodriguez-Fos, Merce Planas-Felix, Daniel Cameron, Phillip Demarest, Anika Nabila, Helen Mueller, Junfei Zhao, Paul Bergin, Casie Reed, Tzippora Chwat-Edelstein, Alex Pagnozzi, Caroline Nava, Emilie Bourel-Ponchel, Patricia Cornejo, Ali Dursun, R. Köksal Özgül, Halil Tuna Akar, Reza Maroofian, Henry Houlden, Huma Arshad Cheema, Muhammad Nadeem Anjum, Giovanni Zifarelli, Miriam Essid, Meriem Ben Hafsa, Hanene Benrhouma, Carolina Isabel Galaz Montoya, Alex Proekt, Xiaolan Zhao, Nicholas D. Socci, Matthew Hayes, Yves Bigot, Raul Rabadan, David Torrents, Claudia L Kleinmann, Michael C. Kruer, Miklos Toth, Alex Kentsis

## Abstract

DNA transposable elements and transposase-derived genes are present in most living organisms, including vertebrates, but their function is largely unknown. PiggyBac Transposable Element Derived 5 (PGBD5) is an evolutionarily conserved vertebrate DNA transposase-derived gene with retained nuclease activity in human cells. Vertebrate brain development is known to be associated with prominent neuronal cell death and DNA breaks, but their causes and functions are not well understood. Here, we show that PGBD5 contributes to normal brain development in mice and humans, where its deficiency causes disorder of intellectual disability, movement, and seizures. In mice, Pgbd5 is required for the developmental induction of post-mitotic DNA breaks and recurrent somatic genome rearrangements. In the brain cortex, loss of Pgbd5 leads to aberrant differentiation and gene expression of distinct neuronal populations, including specific types of glutamatergic neurons, which explains the features of PGBD5 deficiency in humans. Thus, PGBD5 might be a transposase-derived enzyme required for brain development in mammals.

**One-Sentence Summary:** PiggyBac Transposable Element Derived 5 (PGBD5) is required for brain development in humans and mice through genetic and epigenetic mechanisms.

---

Vertebrate brain development requires neuronal cell diversification and self-organization into signaling networks (*1*). While cell diversification is required for the development of many tissues, the development of nervous and immune systems is also uniquely dependent on DNA break repair and developmental apoptosis (*2–9*). For example, several human DNA damage repair deficiency syndromes, such as ataxia telangiectasia (AT) and Seckel syndromes, exhibit both abnormal brain neuron development and immune lymphocyte deficiencies. Likewise, mice deficient for the evolutionarily conserved DNA repair factors, such as *Xrcc5/Ku80*, are also characterized by abnormal neuron and lymphocyte development. In developing immune lymphocytes, efficient end-joining DNA repair is required to ligate DNA breaks induced by the domesticated DNA transposase RAG1/2 during somatic diversification of immunoglobulin receptor genes. Somatic genetic neuronal diversification of cell adhesion receptors was originally proposed more than 50 years ago to provide a mechanism for the complex organization of vertebrate brains (*10*). Initially considered for clustered protocadherins based on their structural similarity to the immunoglobulin receptor genes (*11, 12*), somatic DNA breaks have now been detected in a diverse set of neuronal genes (*13, 14*). Indeed, recent single-cell sequencing studies have found extensive somatic genetic mosaicism in human neurons (*13, 15–17*), as bolstered by numerous prior studies in mice (*13, 18, 19*). While somatic DNA rearrangements and developmental apoptosis are known to be essential for the evolution and function of vertebrate adaptive immunity, the mechanisms of somatic DNA breaks and neuronal apoptosis during brain development remain obscure.

Recently, we found that PiggyBac Transposable Element Derived 5 (PGBD5), the most evolutionarily conserved domesticated DNA transposase-derived gene in vertebrates, is expressed in most childhood solid tumors where it mediates sequence-specific oncogenic DNA rearrangements dependent on its putative nuclease activity and cellular end-joining DNA repair (*20–22*). PGBD5-induced DNA rearrangements in human cells have been validated by multiple laboratories (*20, 23*), and recently also confirmed independently by Bigot et al (*24*). Since most PGBD5-expressing childhood solid tumors share a common neuroectodermal developmental origin (*21, 24, 25*), we hypothesized that PGBD5 may be required for normal nervous system development, at least in part by mediating somatic DNA rearrangements in developing neurons. Indeed, PGBD5 is expressed predominantly in nervous system tissues, and the brain in particular (Fig. 1A-B & S1A-B), with the highest expression in glutamatergic neurons followed by GABAergic neurons, both in humans (Fig. S2A&C) and mice (Fig. S2B&D).

**Fig. 1.**
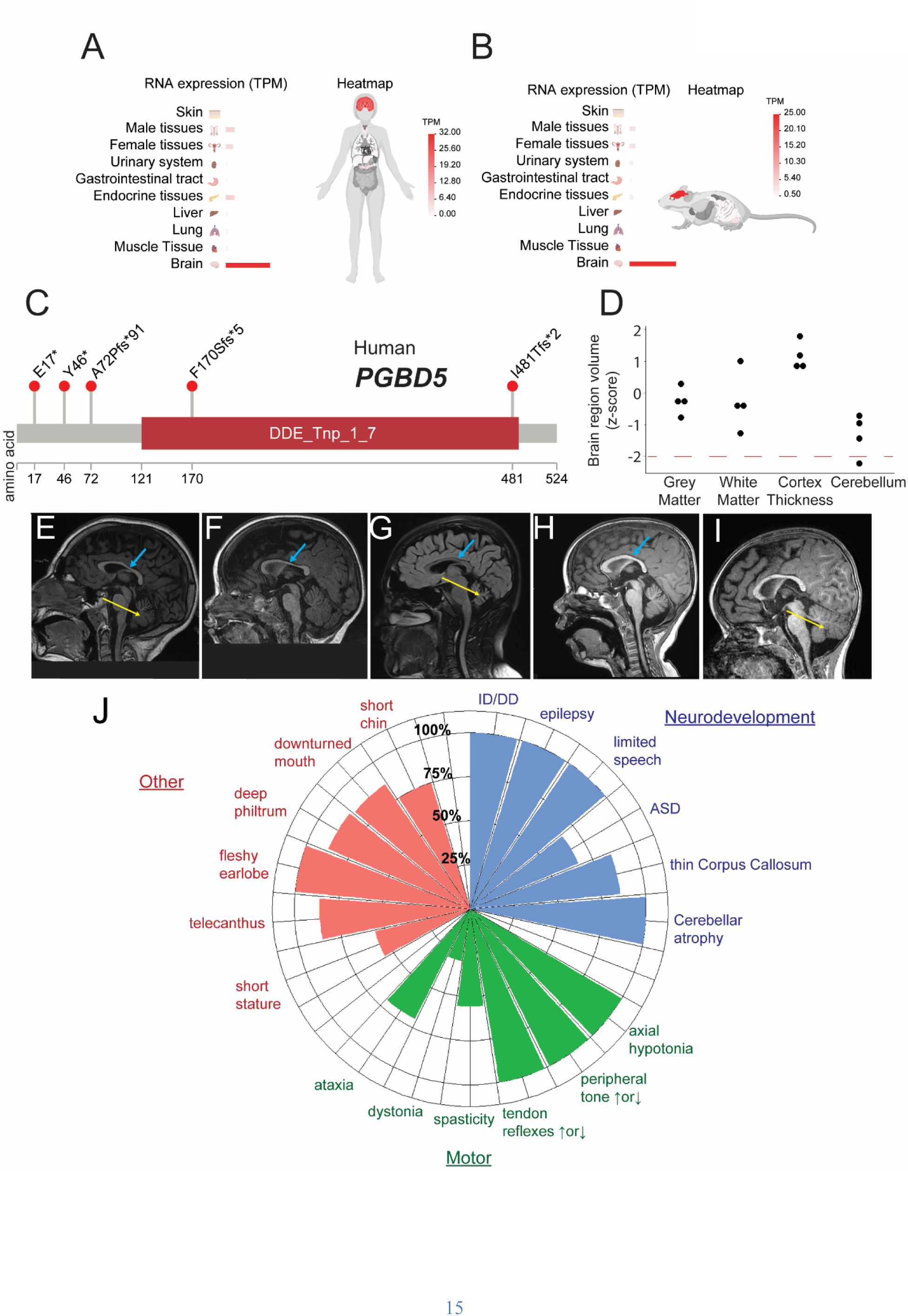
*PGBD5* is specifically expressed in neuronal tissues and its deficiency in humans is associated with abnormal brain development. (**A-B**) Bar graphs showing specific neuronal tissue expression of *PGBD5* in human (**A**) and mouse (**B**) tissues. Color gradient from red to white indicates gene expression in transcripts per million reads (TPM). (**C**) Schematic of the primary structure of PGBD5 with observed genetic variants, most of which appear to be loss-of-function upstream of the evolutionarily conserved DDE_Tnp_1_7 transposase domain (red box). **D,** Comparison of volumes of different brain structures between 4 PGBD5 patients and age/sex matched controls. Dashed red line indicates 2 standard deviations from controls. **E-I,** Sagittal MRI brain images demonstrating thin corpus callosum (thick blue arrow) in patients 2 years and older and decreased cerebellar size (thin yellow arrow) in patients 6 years and older. **E,** Patient 1.1 (10 years) with progressive cerebellar atrophy determined after repeat imaging as compared to 4 years of age, **F**, Patient 1.2 (2 years) with CC thinning present, **G,** Patient 2.1 (15 years) with marked cerebellar atrophy, **H,** Patient 5.1 (3 years) with some thinning of CC, **I**, Patient 5.2 (21 months) with cerebellar atrophy. **J**, Phenogram summarizing frequency of conserved features in neurodevelopmental, motor, and congenital anomaly domains. Frequency calculated using patients with data provided, excluding N/A from denominator. ID/DD=intellectual disability/developmental delay, ASD=autism spectrum disorder.

To investigate the function of PGBD5 in human brain development, we identified five unrelated consanguineous families with *PGBD5* mutations using GeneMatcher (*26*). Exome sequencing analysis demonstrated distinct homozygous *PGBD5* mutations segregating with affected family members. We confirmed the observed *PGBD5* mutations using genomic PCR and Sanger sequencing (Figure S1C, Table S1). *PGBD5* mutations in affected individuals consisted of predicted nonsense and frameshift variants, most of which occurred upstream of the evolutionarily conserved aspartate triad thought to be required for the biochemical activity of the *PGBD5* transposase-homology domain in cells (Fig. 1C). Expression of cDNAs encoding the observed *PGBD5* c.49G>T (E17*) and c.509del (F170Sfs*5) mutations led to substantial reduction of PGBD5 protein in HEK293T cells (Fig. S1D-E). While additional studies will be needed to establish the effects of observed mutations on endogenous loci in PGBD5-expressing cells, at least some of the phenotypes of the affected individuals can be attributed to the loss of PGBD5 protein.

Importantly, affected children with inherited PGBD5 mutations shared conserved clinical phenotypes across neurodevelopmental and motor domains (Fig. 1D-J; Supplemental Clinical Summaries; Table S2). While quantification of brain MRI volumes did not identify quantitatively significant changes (Fig. 1D), visual analysis identified thin corpus callosum (6/7) and reduced cerebellar size associated with widening of the vermis folia (7/7) which became more apparent in patients older than 6 years or on follow-up imaging (Fig. 1E-I). Neurodevelopmental features include intellectual disability and developmental delay (ID/DD; 10/10), epilepsy (9/9), limited or no speech (9/9), autism spectrum disorder or social delay (ASD; 4/6). Prominent motor features included axial hypotonia (9/9), increased peripheral tone (7/9) or decreased peripheral tone (2/9), increased tendon reflexes (5/9) or decreased tendon reflexes (4/9). Less frequently, we observed spasticity mainly affecting the legs (5/9), intermittent dystonia (3/10), and ataxia (7/10). Some patients were of short stature (5/9), although height and head circumference were generally normal. We noted some dysmorphic features, including telecanthus (6/7), fleshy earlobes (8/8), deep philtrum (7/8), downturned mouth corners (6/7), and short chin (6/8; Fig. 1J; Data S1 and Table S2). While additional patients will be needed to define the full spectrum of human PGBD5 deficiency syndrome, these findings indicate that PGBD5 mutations are associated with developmental delay, intellectual disability, ataxia-dystonia, and epilepsy.

To investigate the physiologic functions of PGBD5 in nervous system development, we used a dual recombinase-mediated cassette exchange to engineer *Pgbd5^fl/fl^* mice, in which *Pgbd5* exon 4 is flanked by *loxP* sites (*27*). We bred *Pgbd5^fl/fl^* mice with *EIIa-Cre* mice to generate *Pgbd5^-/-^*mice (Fig. S3A), as confirmed by genomic PCR (Fig. S3C) and Sanger sequencing. *In situ* hybridization microscopy analysis of the hippocampus, which has some of the highest density of Pgbd5-expressing neurons (Figs. S1A-B, S2A-D, S3B), revealed no measurable Pgbd5 exon 4 transcript expression in *Pgbd5^-/-^* mice (Fig. S3B), though RNA sequencing analysis revealed residual *Pgbd5* transcripts lacking exon 4, consistent with potential retention of truncated alleles lacking the transposase domain (Fig. S3D). We also engineered *Pgbd5^3xFLAG-HA-P2A-eGFP^* knockin mice which permit specific detection of endogenous Pgbd5 expression in cells (Fig. S4A) and confirmed that Pgbd5 is expressed in neurons but not astrocytes or microglia, as established by specific co-staining with NeuN, GFAP, and TMEM119, respectively (Fig. S4B-D).

We found that both *Pgbd5^-/-^* and their *Pgbd5^wt/-^* littermate mice were born at the expected Mendelian ratios (Fig. S5A-B), but *Pgbd5^-/-^* mice were runted and had significantly smaller brains, as compared to their wild-type littermates (t-test *p* = 3.4E-3 and 1.6E-2, for females and males, respectively; Fig. S5C-H). Given the neurodevelopmental deficits associated with PGBD5 deficiency in humans, we used specific behavioral tests correlating with features of human PGBD5 deficiency to examine *Pgbd5^-/-^*mice (Fig. S6A-C). Automated video tracking locomotor test analysis revealed significantly increased locomotor activity of *Pgbd5*^-/-^ and *Pgbd5^wt/-^* mice, as compared to their wild-type littermates (ANOVA *p* = 5.4E-7 and 5.5E-7, for females and males, respectively; Fig. 2A-B). We assessed anxiety using the elevated plus maze test (EPM; Fig. 2C). Both female and male *Pgbd5*^-/-^ mice traveled longer distances in the open maze arms (normalized to total distance traveled) as compared to their wild-type littermates, indicating reduced avoidance of the anxiogenic open arm of the EPM, consistent with reduced anxiety-like behavior (ANOVA *p* = 2.7E-6; Fig. 2D). Reduced avoidance of female and male *Pgbd5*^-/-^ mice was also reflected by their increased entry and time spent in the open arms (Fig. S6A-B). *Pgbd5^wt/-^* females and males exhibited an intermediate phenotype in open arm time and entries indicating that a partial deficit in *Pgbd5* expression is sufficient to elicit the EPM phenotype.

**Fig. 2.**
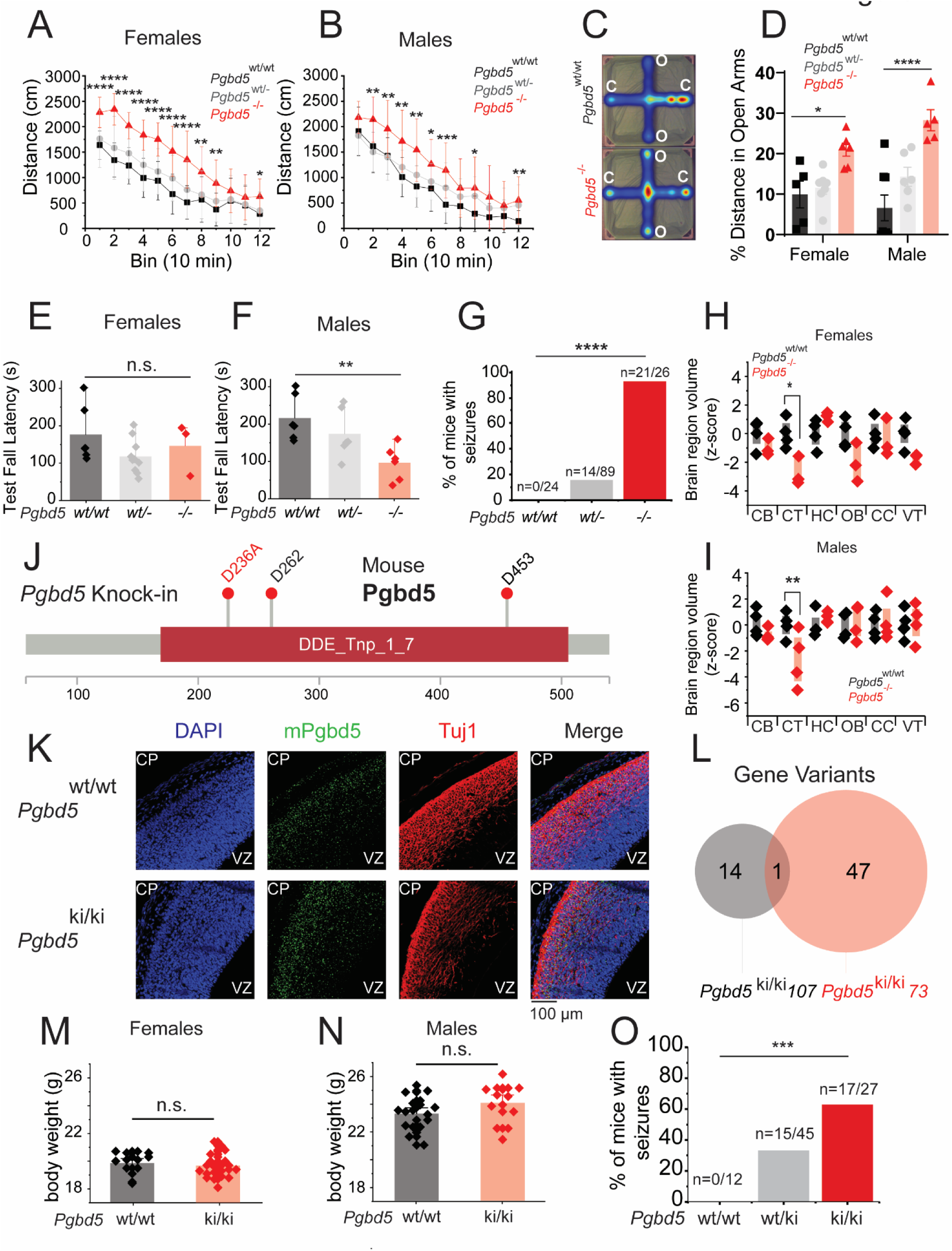
*Pgbd5* knock-out and knock-in mice reproduce behavioral and brain developmental deficits associated with human *PGBD5* mutation. **A-B**, Whisker plot analysis of the distance traveled in locomotor assays of 12-week old female (**A**) and male (**B**) *Pgbd5^wt/wt^* (black), *Pgbd5^wt/-^* (grey), and *Pgbd5^-/-^* (red) mice, demonstrating significantly increased activity of *Pgbd5*-deficient mice (Female and male *n*=12 and two-way ANOVA *p* = 5.41E-7 and *n*=12 and *p* = 5.47E-5, respectively; post-hoc Tukey test \**p*<0.05, ** *p*<0.01, \*\*\**p*<0.001, \*\*\*\**p*<0.0001). **C**, Representative heatmaps of elevated plus maze assay (color index from dark blue (low) to red (high) indicates time spent in the area), with **D**, bar plot of the percentage of the distance traveled in the open arm by 12-weeks old *Pgbd5^wt/wt^*(black), *Pgbd5^wt/-^* (grey), and *Pgbd5^-/-^* (red) mice. *Pgbd5*-deficient mice exhibit a significantly increased propensity to explore the open arms (Two-way ANOVA *p* = 2.6E-6 for genotype and *p =* 0.3 for sex; * Tukey test *p* = 0.019 and **** *p* = 2.67E-7). **E-F**, Bar plots of probe day Rotarod fall latency in females (**E**) and males (**F**), showing significant reduction by *Pgbd5^-/-^* (red) as compared to *Pgbd5^wt/wt^*in male mice (** = One-way ANOVA *p* =9.2E-3, Tukey’s test *p* = 7.5E-3 in males; n.s= One-way ANOVA *p* =0.2, Tukey’s test *p* = 0.76 ** ANOVA. **G**, Bar plot of seizure activity of *Pgbd5^-/-^* versus *Pgbd5^wt/wt^* litter mate mice (χ^2^-test *p* = 5.8E-7). *n* indicates number of mice with seizures over the total number of mice assayed. **H-I**, Box plots of *z*-scores of brain MRI volumetric measurements of 60-day old *Pgbd5^-/-^* as compared to *Pgbd5^wt/wt^* mice, showing significant reduction in cortex and ventricle size brain regions in *Pgbd5*-deficient female (*= Two-way ANOVA *p =* 9E-4, Cortex Bonferroni-adjusted *p=*5.7E-3) (**H**) and male (**= Two-way ANOVA *p=*0.28, Cortex Bonferroni-adjusted *p=*1.9E-2) (**I**) mice. **J,** Pgbd5 primary protein sequence schematic indicating the location of conserved aspartate triad and in red, the exon 2 D236A (ki) substitution. ENSMUST00000140012.8 Pgbd5 transposase domain highlighted in red. **K**, Representative fluorescence *in situ* hybridization micrographs of coronal sections of heads of 14.5-day old *Pgbd5^ki/ki^* as compared to *Pgbd5^wt/wt^* litter mate embryos, showing similar expression of *Pgbd5* transcripts between *Pgbd5^wt/wt^* and *Pgbd5^ki/ki^* litter mates. Green staining indicates mouse *Pgbd5* RNA, red indicates Tuj1 staining of postmitotic neurons and blue denotes nuclei stained with DAPI. Scale bar = 100 μm. **L**, Venn diagram of gene variants detected in the whole-genome sequencing of *Pgbd5^ki/ki^ 107* and *Pgbd5^ki/ki^73* founder lines. Only the *Pgbd5* gene variant was found to be shared between founder lines. **M-N,** Total body weight of 60-day old *Pgbd5^wt/wt^*(black), and *Pgbd5^ki/ki^*(red) mice, shows no difference of total weights in females (n.s. *p =* 0.3) (**M**) and males (n.s. *p =* 0.6) (**N**) mice. **O**, Bar plot of seizure activity *Pgbd5^-/-^*versus *Pgbd5^ki/ki^* litter mate mice (χ^2^-test *p* = 7.41E-05). n indicates number of mice with seizures over total number of mice assayed; O = open arm, C = closed arm, CB = cerebellum, CT = cortex, HC = hippocampus, OB = olfactory bulb, CC = corpus callosum, VT = ventricles.

Prompted by the motor deficits of *PGBD5*-deficient humans, we assayed *Pgbd5*-deficient mice using the Rotarod performance test (Figs. 2E-F & S6D-J). Despite having no significant differences in grip strength (*p* = 0.08; Fig. S6H), *Pgbd5^-/-^* mice exhibited significantly reduced rotarod fall latency, consistent with impaired motor learning in males (One-way ANOVA *p* = 9.2E-3, post-hoc Tukey’s test *p* = 0.5E-3; Fig. 2E-F). Both male and female *Pgbd5*^-/-^ mice also exhibited thermal hypersensitivity (Fig. S6K), without any apparent gait effects (Fig. S6I-J). Lastly, we assayed *Pgbd5*-deficient mice for susceptibility to seizures. We found that most *Pgbd5^-/-^* mice developed partial motor and generalized tonic-clonic seizures in response to stressful handling as compared to their wild-type littermates (χ^2^-test *p* = 5.8E-7; Fig. 2G). To investigate the anatomic basis of this complex behavioral syndrome, we used high-resolution manganese-enhanced MRI (MEMRI) to analyze brain architecture in *Pgbd5*-deficient mice. This revealed significant reductions in the cortical volumes in *Pgbd5^-/-^* male and female mice, as assayed using quantitative volumetric mouse brain atlas analysis (ANOVA Bonferroni-adjusted *p =* 1.9E-2 and 9E-4, respectively; Fig. 2H-I, Fig. S7B-C). Overall, *Pgbd5*-deficient mice display complex behavioral deficits, including seizures, behavioral and motor deficits, and structural brain abnormalities that resemble the human PGBD5 deficiency syndrome.

PiggyBac-type enzymes utilize conserved aspartate residues to catalyze DNA hydrolysis and rearrangements, with analogous residues required for the cellular DNA remodeling activities of PGBD5 (*21, 28*), though whether PGBD5 functions enzymatically as a transposase, recombinase, or a different type of nuclease needs to be determined (*24*). To test whether mouse brain development requires Pgbd5 nuclease activity, we used CRISPR engineering to generate *Pgbd5^D236A/D236A^*knock-in mice (*Pgbd5^ki/ki^*; Fig. 2J & S8A), in which one of the evolutionarily conserved aspartate residues required for cellular DNA activity was mutated to inactive alanine (*20*). We confirmed that the analogous mutation in human *PGBD5* does not impair cellular protein stability by Western immunoblotting or its ability to associate with chromatin by ChIP-seq (*20, 21*). We verified *Pgbd5^D236A^* mutation in two independent founder strains using genomic PCR and Sanger sequencing (Fig. S8A), germ-line transmission by restriction enzyme mapping (Fig. S8B), and lack of off-target gene mutations by whole-genome sequencing (Fig. 2L; Table S3). Homozygous *Pgbd5^ki/ki^* mice exhibited physiologic and unaltered expression of endogenous Pgbd5 mRNA as compared to their *Pgbd5^wt/wt^*littermates, as assessed using in situ hybridization with *Pgbd5*-specific probes (Fig. 2K). *Pgbd5^ki/ki^* mice showed no difference in body weights as compared to their wild-type littermates (Fig. 2M-N) but exhibited tonic-clonic seizures similar to *Pgbd5*-deficient mice (χ^2^-test *p* = 7.4E-05; Fig. 2O). Thus, brain functions of Pgbd5 at least in part require specific aspartate activity of its transposase-homology domain.

Mammalian neurogenesis occurs largely during embryonic development, with mouse cortex development peaking in the mid-to-late embryos (*29*). Given the well-defined layered organization of the mouse motor cortex, we analyzed its architecture in 14.5-day old (E14.5) embryos. Prior studies of this developmental period have also documented that post-mitotic neurons accumulate extensive DNA breaks and activate end-joining DNA repair as they migrate to the mantle layer upon differentiation of progenitor neuroblasts in the ventricular zone (*2, 3, 7*). Thus, we used a neuron-specific tubulin isoform Tuj1 as a specific marker of post-mitotic neurons (*30*), and immunofluorescence microscopy to examine post-mitotic neurons in the brain cortices of 14.5-day old embryonal (E14.5) *Pgbd5^-/-^* mice (Fig. 3A-C & S9-S13).

**Fig. 3.**
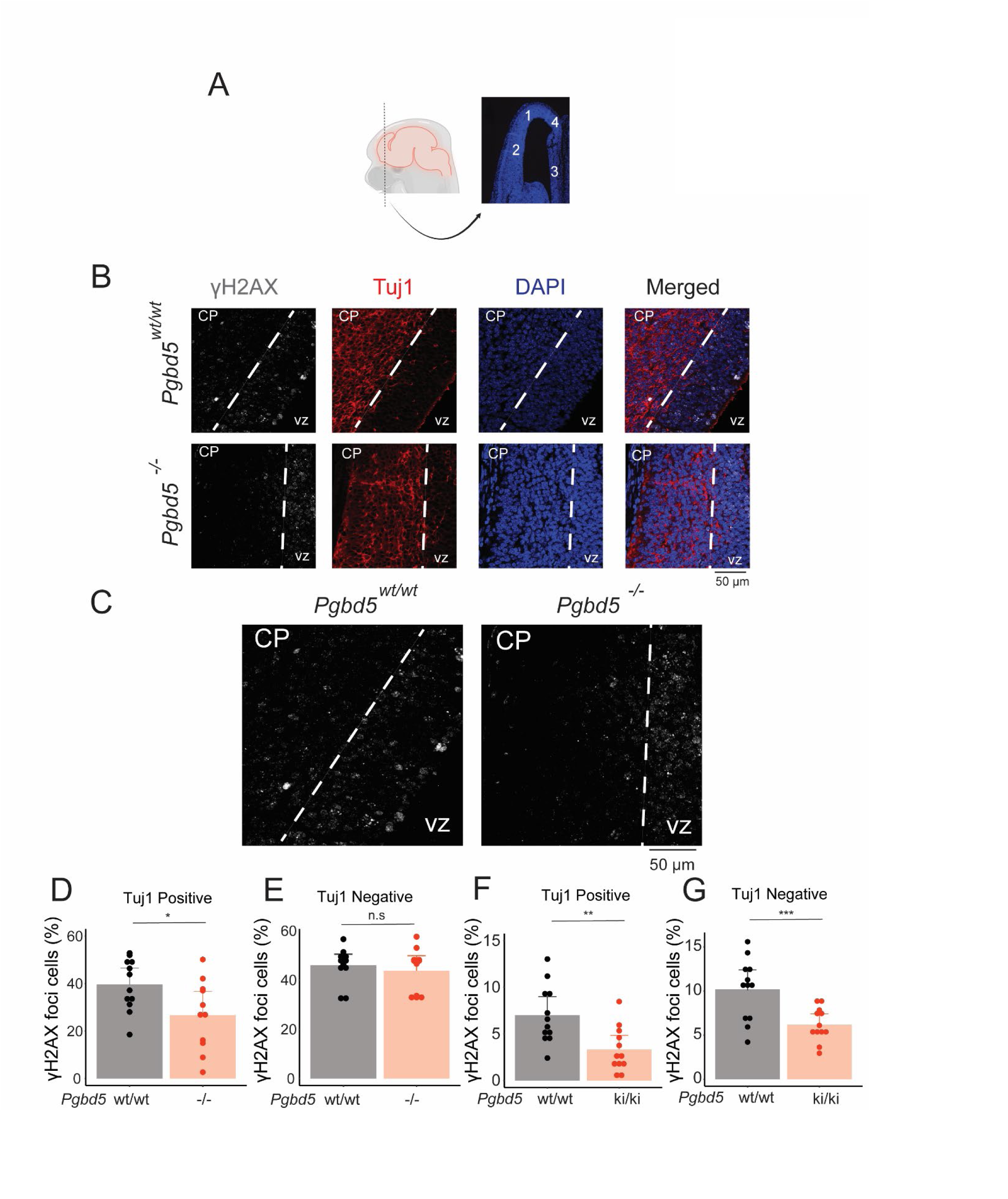
*Pgbd5* is required for developmental induction of DNA breaks in postmitotic cortical neurons. **A**, Schematic showing representative coronal section of a 14.5-days old embryo mouse forebrain and the regions selected for further quantification. **B,** Representative immunofluorescence micrographs of *Pgbd5^wt/wt^* and *Pgbd5^-/-^* 14.5-day old litter mate embryos. DAPI shown in blue stains nuclei, γH2AX in white indicates sites of double-strand DNA break repair, and Tuj1 in red marks differentiated postmitotic neurons; CP = cortical plate, VZ = ventricular zone. **C**, Enlarged representative γH2AX immunofluorescence micrographs from panel A of *Pgbd5^wt/wt^* (left) and *Pgbd5^-/-^* (right) 14.5-day old litter mate embryos stained for γH2AX in white. **D-G**, Quantification of γH2AX in postmitotic (Tuj1 positive) and proliferating neurons (Tuj1 negative) in the *Pgbd5^wt/wt^* and *Pgbd5^ki/ki^*mice. **D**-**E,** Bar plots showing the percentages of cells with punctate γH2AX staining in Tuj1-positive (**D**) and Tuj1-negative neurons (**E**) in *Pgbd5^wt/wt^* versus *Pgbd5^-/-^* mice (t-test \**p* = 0.029 and n.s. *p =* 0.6 for % of positive cells). **F**-**G,** Bar plots showing the percentages of cells with punctate γH2AX staining in Tuj1-positive (**F**) and Tuj1-negative neurons (**G**) in *Pgbd5^wt/wt^*versus *Pgbd5^ki/ki^* mice (t-test \*\**p* = 3.8E-3 and \*\*\**p* = 2.9E-3 for Tuj1 positive and negative cells, respectively).

Using γH2AX as a specific surrogate of neuronal DNA break repair (*31*), we observed a significant reduction in the number of neurons with γH2AX foci specifically among post-mitotic (Tuj1-positive), as compared to proliferating (Tuj1-negative) neuronal precursor in *Pgbd5^-/-^* mice as compared to their wild-type littermates (t-test *p* = 0.029; Fig. 3D-E & S14A-B). We confirmed the specificity of this effect by analyzing the fraction of Tuj1-negative neurons in brain cortical neurons, which showed no significant differences between *Pgbd5*-deficient and wild-type littermate brains (Fig. 3D-E & S15A). The observed Pgbd5-dependent neuronal γH2AX foci were specifically induced during cortical neuronal development in E14.5 embryos, as analysis of E12.5 brain cortices which contain only a single mantle layer revealed no significant differences (Fig. S16). Importantly, *Pgbd5^ki/ki^* mice also exhibited significant reduction of γH2AX foci as compared to wild-type littermate controls (t-test *p* = 3.8E-3 and 2.9E-3 for Tuj1-positive and negative neurons, respectively; Fig. 3F-G) Thus, Pgbd5 and enzymatic activity of transposase-homology domain are specifically required for the developmental induction of DNA breaks and/or their resolution during cortical brain development.

While Pgbd5 contains an evolutionarily conserved and transposase-derived gene with nuclease cellular activity (*20, 21*), it is possible that the observed DNA breaks in neurons occur independently of its DNA breakage activity. To determine whether Pgbd5-dependent neuronal DNA breaks require DNA double-strand break repair, we analyzed genetic interaction between *Pgbd5* and *Xrcc5/Ku80*, the key factor in non-homologous end-joining (NHEJ) DNA repair (Fig. 4E). NHEJ DNA repair is required for the ligation of double-strand DNA breaks induced by many ‘cut-and-paste’ DNA transposase enzymes and their domesticated derivatives like PGBD5 and RAG1/2 (*32*). Similar to human DNA damage repair deficiency syndromes, such as AT and Seckel syndromes, *Xrcc5^-/-^* mice have neurodevelopmental defects, associated with unrepaired DNA breaks and extensive neuronal apoptosis during cortical development, as well as severe combined immunodeficiency due to the failure to repair RAG1/2-induced DNA breaks and rearrangements in developing lymphocytes (*33*).

**Fig. 4.**
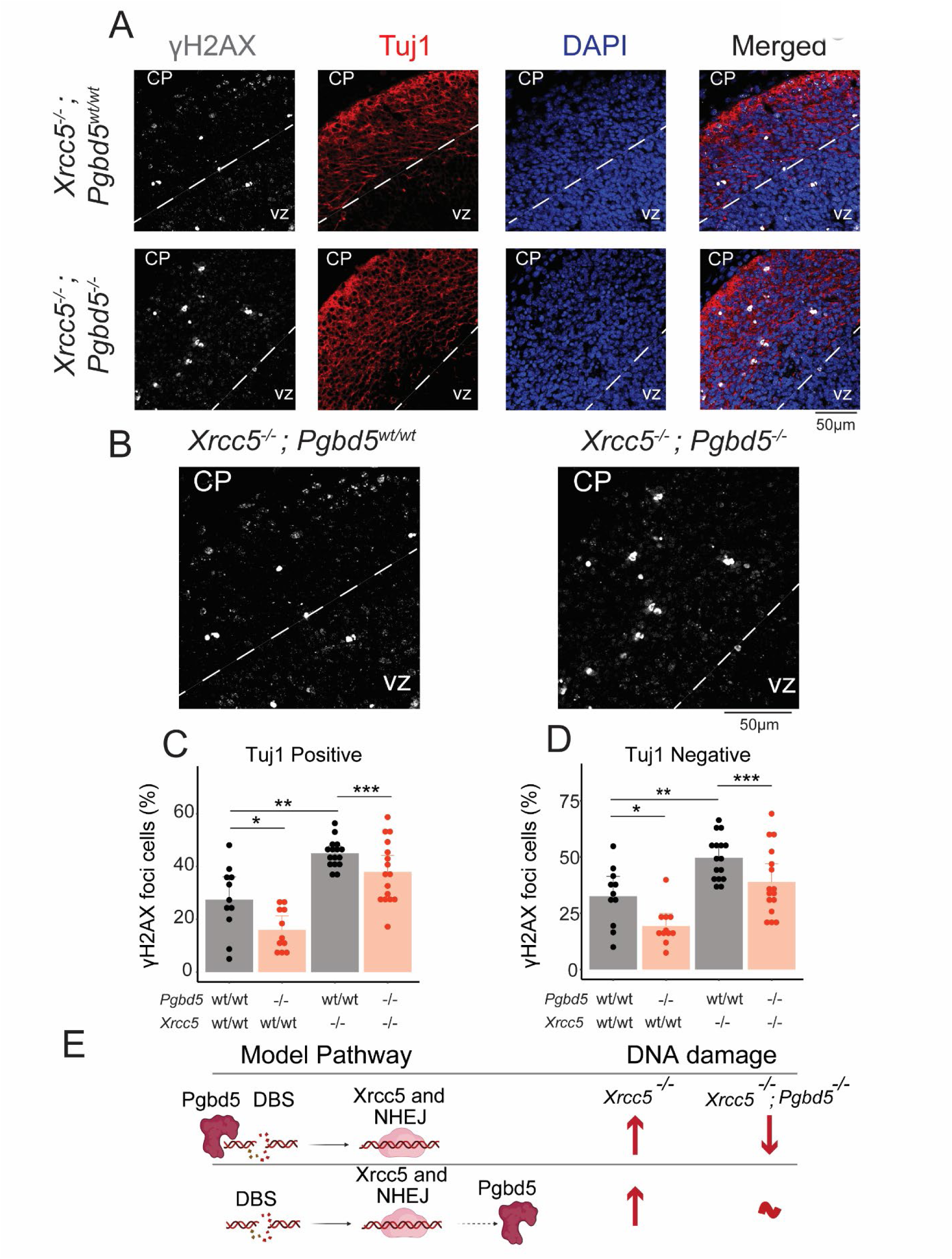
*Xrcc5* is required for *Pgbd5*-induced double-strand DNA break repair. **A**, Representative immunofluorescence micrographs of *Xrcc5^-/-^;Pgbd5^wt/wt^* (top) and *Xrcc5^-/-^;Pgbd5^-/-^* (bottom) 14.5-day old litter mate embryos stained for γH2AX. DAPI nuclear staining is shown in blue; γH2AX indicates sites of double-strand break repair (white), and Tuj1 marks postmitotic neurons (red); CP = cortical plate, VZ = ventricular zone. **B**, Enlarged representative immunofluorescence micrographs of *Xrcc5^-/-^;Pgbd5^wt/wt^*and *Xrcc5^-/-^;Pgbd5^-/-^* 14.5-day old litter mate embryos from panel A. **C-D**, Quantification of nuclear γH2AX in postmitotic neurons (Tuj1 positive) and proliferating neurons (Tuj1 negative). Bar plots showing percentages of cells with punctate γH2AX staining in Tuj1-positive (**C;** t-test \**p* = 2.3E-2, \*\**p* = 1E-3, and \*\*\**p* = 4.1E-2) and Tuj1-negative neurons (**D;** t-test \**p* = 0.012 and \*\**p* = 1.8E-3, \*\*\**p* = 0.024 for postmitotic and proliferating neurons, respectively). **E**, Schematic showing potential genetic interaction models between *Pgbd5* and *Xrcc5* in cortical neuronal developmental DNA break repair. Arrows denote relative levels of DNA damage.

First, we confirmed that *Xrcc5^-/-^* mice failed to produce normal T- and B-lymphocytes, as assayed by fluorescence-activated cell scanning (FACS) using CD4/CD8 and B220/IgM-specific antibodies, respectively, as compared to their wild-type or *Pgbd5^-/-^* littermates (Figs. S21I-J & S22). In agreement with prior studies, *Xrcc5^-/-^*mice showed a significant increase in the number of neurons with γH2AX breaks as compared to their wild-type littermates in post-mitotic neurons (t-test *p* = 4.1E-2; Fig. 4C) and proliferative progenitors (t-test *p* = 2.4E-2; Fig. 4D). Remarkably, we found that *Pgbd5^-/-^;Xrcc5^-/-^* mice had significantly reduced DNA damage as compared to their *Pgbd5^wt/wt^;Xrcc5^-/-^* littermates (t-test *p* = 4.1E-2 and 2.4E-2 for Tuj1-negative and positive neurons, respectively; Figs. 4C-D, S14C-D, and S17-20). Thus, Pgbd5-induced neuronal DNA damage repair requires Xrcc5 (Fig. 4E). Commensurate with the physiologic function of developmental neuronal DNA break repair, we found that *Pgbd5^-/-^;Xrcc5^-/-^* mice were similarly runted and failed to thrive as compared to their *Pgbd5^wt/wt^;Xrcc5^-/-^* littermates (Fig. S21A-H & S22), which was associated with increased neuronal cell death as measured by terminal deoxynucleotidyl transferase dUTP nick end labeling (TUNEL) specifically in E14.5 but not E12.5 brains (t-test *p* = 1.5E-2 and *p =* 0.89, respectively; Fig. S23). In all, these findings indicate that Pgbd5 is required for the developmental induction of DNA breaks and cortical brain development.

To determine whether Pgbd5 induces somatic DNA rearrangements during brain development, we used PCR-free paired-end Illumina whole-genome sequencing (WGS) of diverse anatomically dissected brain regions from multiple individual *Pgbd5*-deficient and wild-type littermate mice. Current single-cell DNA sequencing methods enable accurate detection of single nucleotide variation, but their requirements for DNA amplification prevent the detection of larger rearrangements, such as those expected from DNA nucleases (*34*). While bulk PCR-free DNA sequencing is not sufficiently sensitive to detect DNA rearrangements occurring in single neurons, we reasoned that if Pgbd5 functions as a somatic neuronal DNA nuclease, its developmental activity would yield recurrent somatic signals in multiple diverse wild-type but not *Pgbd5^-/-^*litter mate brains via involvement of shared loci and/or sequences in bulk cell sequencing.

We tested this conjecture by analyzing somatic DNA rearrangements observed using PCR-free paired-end Illumina WGS analysis of peripheral blood mononuclear cells (PBMC) isolated from 30-day old mice (mean genome coverage 90-fold, Fig. 5A). First, we validated that our analysis was not biased by sequencing coverage (Fig. S24A-B) and produced accurate detection of somatic DNA variants based on the allele frequencies in matched tissues (Fig. S24C-F). We analyzed the resultant sequencing data using recently developed methods optimized for the accurate detection of somatic genome variation (*35–37*).

**Fig. 5.**
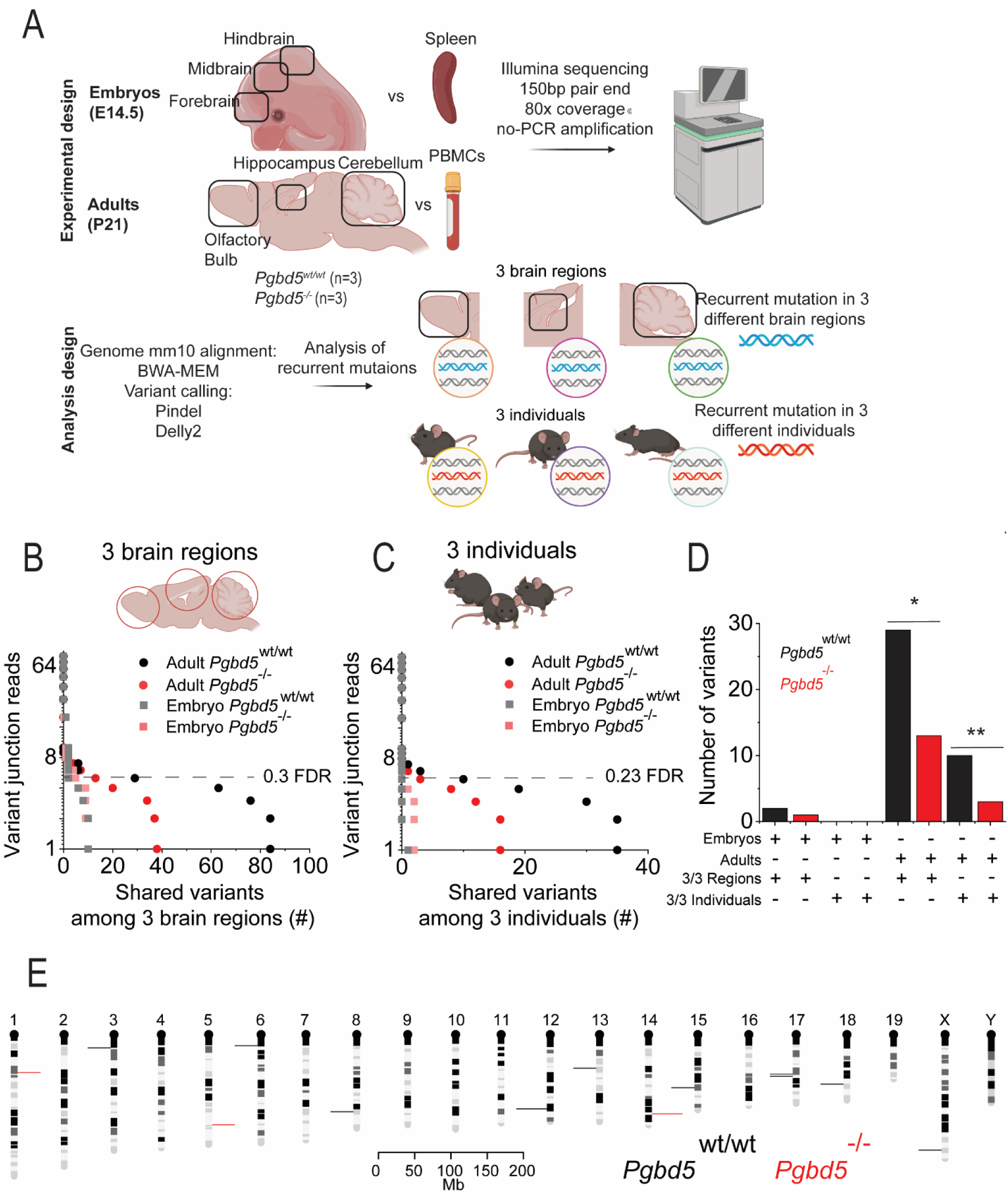
*Pgbd5* is required for recurrent somatic DNA rearrangements in developing mouse brain. **A,** Schematics for somatic whole-genome sequencing analysis of neuronal and non-neuronal tissues from three *Pgbd5^wt/wt^*and three *Pgbd5^-/-^* adult and embryonal littermate mice. **B-D**, Dot plots showing numbers of somatic structural variants at different variant junction read thresholds shared across cerebellum, hippocampus, and olfactory bulb brain regions (**B**) and three individuals (**C**) in adult and embryonal *Pgbd5^wt/wt^* (black circles and grey squares, respectively) and *Pgbd5^-/-^* (red circles and light-red squares, respectively). The overlap among structural variants was calculated using the breakpoint analysis (Fig. S26D): 5’ and 3’ DNA breakpoint ± 350bp requiring an overlap of at least 1%. There are significantly more recurrent somatic structural variants in *Pgbd5^wt/wt^* with support of at least 5 variant junction reads in the recurrent events shared among three individuals and three brain regions (*χ^2^-test *p* = 2.2E-145 and 2.8E-133, respectively). (**D**) Bar plot summarizing the results from **B** and **C** using the support threshold of at least 5 variant junction reads. Significant differences between the number of recurrent somatic DNA rearrangements between adult *Pgbd5^wt/wt^* and *Pgbd5^-/-^*shared among three individuals and three brain regions (**χ^2^-test *p* = 1.6E-17 and 1.3E-9, respectively. **E**, Mouse chromosome ideograms showing the locations of recurrent somatic DNA rearrangements in three individuals observed in *Pgbd5^wt/wt^* (black) and *Pgbd5^-/-^*(red) brains; bin = 1 million bases.

Consistent with the known somatic V(D)J DNA recombination activity of RAG1/2 in blood lymphocytes, we observed somatic deletions of the *Igkj1* and *Igkj2* loci (among other immunoglobulin receptor genes) with common breakpoints in multiple sequencing reads in PBMC as compared to matched brain tissue (mean variant fraction = 0.015; Fig. S25A). Lack of apparent somatic deletions of *Igkj1* and *Igkj2* or related immunoglobulin gene loci in fetal spleens of 14.5- day old mouse embryos as compared to their matched brain tissue confirmed the specificity of this approach (Fig. S25A), consistent with the known absence of RAG1/2 activity in fetal hematopoietic cells in mouse spleen (*38*). In contrast, we observed clonal deletions of *Pgbd5* exon 4 in both adult PBMCs and fetal spleens of *Pgbd5^-/-^* mice but not in their *Pgbd5^wt/wt^*littermates (Fig. S25B).

Using this comparative approach to detect developmental somatic DNA rearrangements by the domesticated RAG1/2 DNA recombinase in blood cells, we examined somatic genomic variation of brain tissues dissected from *Pgbd5^wt/wt^* and *Pgbd5^-/-^* littermate mice. We performed independent analyses to quantify somatic single nucleotide variants (SNVs) and DNA rearrangements such as deletions, and then tested for their recurrence by comparing genomic locations of somatic variants in individual mice or their anatomic brain regions, as explained in detail in the methods section. We observed no significant differences in somatic SNVs in the brain tissues of both 30-day old adult and 14.5-day old embryonal *Pgbd5^wt/wt^* as compared to their *Pgbd5^-/-^*litter mate mice, consistent with their equal chronological and biological age (median allele fraction = 0.096 and 0.094 for adult *Pgbd5^wt/wt^*and *Pgbd5^-/-^* mice, respectively; Fig. S23A-B).

In agreement with the stochastic nature of somatic nucleotide substitutions, most of which are due to DNA replication errors in proliferating tissues (*39*), we also found no genomic regions that recurrently accumulated somatic SNVs across different brain regions or different individual mice, which have an apparently random distribution across the mouse genome (Fig. S26A-B; Table S4). We then focused on the analysis of somatic deletions, insertions, and duplications in adult and embryonal *Pgbd5^wt/wt^* brains, as compared to their *Pgbd5^-/-^* litter mate controls (Fig. S27A-F). While we observed some differences in the various types of structural DNA rearrangements, there were no statistically significant differences in the total numbers of somatic DNA rearrangements between *Pgbd5^wt/wt^* and *Pgbd5^-/-^* brains, both in adults and embryos (Fig. S27A-F).

In contrast, individual adult *Pgbd5^wt/wt^* mice showed significantly more recurrent somatic DNA rearrangements both among different individual mice and their cerebella, hippocampi, and olfactory bulbs, as compared to their *Pgbd5^-/-^* litter mates. This was detected using both recurrence of somatic structural rearrangement breakpoints and their complete overlaps (24 versus 12, and 10 versus 2, respectively; χ^2^-test *p* = 1.3E-9 and 1.6E-17, respectively; Figs. 5A-D & S26C-G). Importantly, there were no significant differences in the recurrence of somatic DNA rearrangements in 14.5-day old embryonal brain tissues isolated from *Pgbd5^wt/wt^* and *Pgbd5^-/-^* litter mate embryos (Fig. 5D & S26G), consistent with the onset of Pgbd5/Xrcc5-dependent DNA break repair during this developmental period.

Finally, recurrent somatic DNA rearrangements shared among different individual mice and brain regions showed distinct genomic distributions (Figs. 5E & S26H; Table S4). Manual inspection of sequencing reads of a subset of DNA rearrangements was consistent with their somatic induction in brain tissues in *Pgbd5^wt/wt^*but not *Pgbd5^-/-^* littermate mice (Fig. S28A-C; Table S4). While the definition of physiologic Pgbd5 genomic targets and their rearranged sequences will require the development of improved single-cell genomic sequencing methods, we propose that the somatically rearranged genomic elements identified here represent signals of developmental physiologic Pgbd5 activity in normal brain development.

To elucidate the specific neuronal populations that may require Pgbd5 activity during brain development, we performed single-nucleus RNA-sequencing (snRNA-seq) combined with assay for transposase-accessible chromatin-sequencing (snATAC-seq) of nuclei isolated from the brain motor cortex of three 21-day old *Pgbd5^wt/wt^* and three *Pgbd5^-/-^* littermate mice (Fig. 6A, Table S5). Upon mapping the observed gene expression onto the developmental ontogeny of normal mouse cortex using two recently established brain atlases (*40, 41*), we clustered the gene expression states of detected nuclei. This identified specific Pgbd5-expressing neuronal populations, as compared to astrocytes, oligodendrocytes and immune cells, most of which lack Pgbd5 expression (Figs. 6B & S29A-C; Table S6). First, we confirmed that the *Pgbd5^wt/wt^* and *Pgbd5^-/-^* brain cortices had relatively equal cellular sampling, consistent with their preserved overall morphologic organization (Fig. S4). We found no significant differences in the apparent proportions of annotated cell types between *Pgbd5^wt/wt^*and *Pgbd5^-/-^* brain cortices (Fig. S30A-B).

**Fig. 6.**
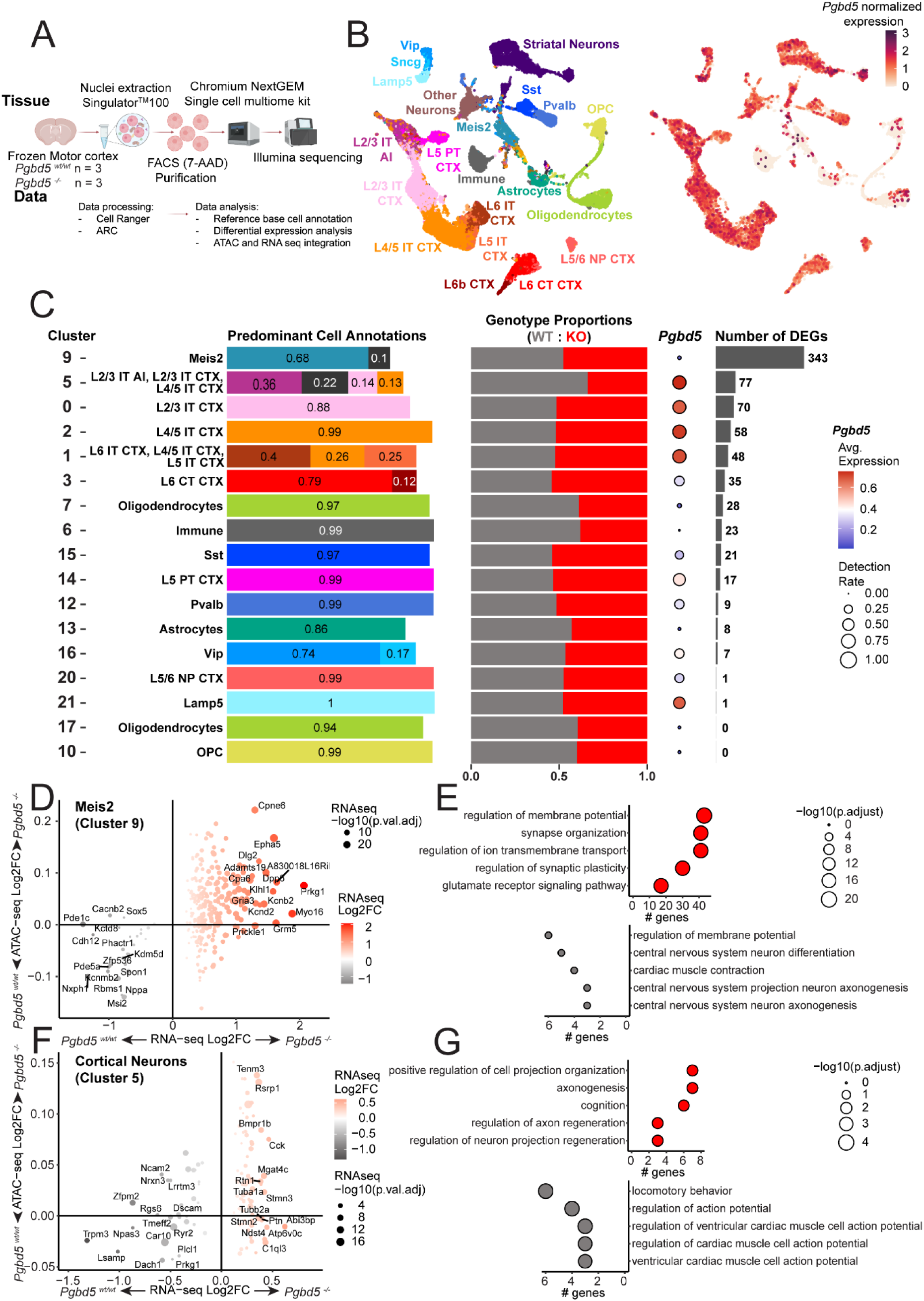
*Pgbd5* deficiency alters gene expression in distinct cortical neurons. **A**, Schematic of experimental procedures for analysis of combined single-nucleus RNA and ATAC sequencing of brain motor cortices from three *Pgbd5^wt/wt^* and three *Pgbd5^-/-^* littermate mice. **B**, Uniform manifold approximation and projection (UMAP) plots of single nuclei gene expression from brain motor cortices of *Pgbd5^wt/wt^* and *Pgbd5^-/-^* littermates, colored by their classification with respect to the reference atlas of normal mouse brain cortex (left; *n* = 18,107 and 14,359, respectively). Right UMAP is colored by *Pgbd5* expression (normalized white to dark red). **C**, Cell clusters with greater than 200 nuclei corresponding to cell populations of cortical origin in *Pgbd5^wt/wt^* (grey) and *Pgbd5^-/-^* (red) mice, excluding ‘Striatal Neurons’ and ‘Other Neurons’. From left to right: proportions of predominant cell type annotations per cluster; proportion of each genotype per cluster; expression (dot color) and detection rate (dot size) of *Pgbd5* expression in wildtype cells of each cluster; number of differentially expressed genes (DEG; log_2_FC > 0.25, adjusted *p* < 0.05) between the genotypes per cluster. **D-G,** Differential expression and promoter accessibility analysis in clusters corresponding to Meis2 cluster 9 (**D-E**) and cortical cluster 5 neurons (**F-G)**. Bubble plots showing changes in gene expression correlated with changes in chromatin accessibility at the corresponding gene promoter regions (+/- 2.5kb from TSS). Only genes with significant changes in expression (adjusted *p* < 0.05) are plotted in Meis2 cluster 9 (**D**) and cortical intratelencephalic (IT) neurons (**F**). Top GO pathways ranked by the number of genes showing adjusted *p*-values in bubble size for DEGs upregulated in knockout (top) and wildtype (bottom) cells of the corresponding cluster in Meis2 cluster 9 (**E**) and cortical IT cluster 5 neurons (**G**).

In contrast, there were significant differences in gene expression of specific populations of neurons with both relatively high and low Pgbd5 expression between *Pgbd5^wt/wt^* and *Pgbd5^-/-^* brains (Fig. 6C). This included the large population of high Pgbd5-expressing intratelencephalic (IT) glutamatergic pyramidal neurons of layers 2/3, 4/5, and 6 that project to other cortical areas and the striatum (*42*), as well as the smaller population of low Pgbd5-expressing Meis2 GABAergic interneurons that are present in the cortical white matter and likely represent projection neuron precursors (*43, 44*) (Fig. 6C). Interestingly, pyramidal tract (PT) neurons, which are the other major cortical pyramidal neurons that project to subcortical structures, as well as cortical GABAergic Pvalb, Sst, Vip interneurons, had relatively few differentially expressed genes, consistent with the distinct function of Pgbd5 in specific neuronal populations (Fig. 6C). Thus, loss of Pgbd5 induces distinct changes in the organization and gene expression of cortical neurons.

We next assessed changes in the chromatin accessibility of promoter regions of differentially expressed genes between *Pgbd5^wt/wt^* and *Pgbd5^-/-^* cells. Significantly affected cortical neuronal populations included glutamatergic cluster 5 layers 2/3 and 4/5 IT and cluster 9 GABAergic Meis2 neurons (Figs. 6F-G & S32A-D). We also observed substantial correlations between differential gene expression and chromatin accessibility of distinct sets of genes (Figs. 6D & 6F). The concordance between Pgbd5-dependent gene expression and chromatin accessibility suggests that Pgbd5 deficiency leads to the dysregulation of specific gene expression within neuronal populations. Gene ontology pathway analysis of differentially expressed genes from the cortical neuronal populations revealed multiple sets of genes involved in the regulation of neuronal membrane potentials, synapse organization, ion channel signaling, and neuronal and axonal projection regeneration, among other neuronal functions (Figs. 6E-G & S31A-B). At least in part, this may explain the phenotype of human PGBD5 deficiency, including developmental delay, intellectual disability, ataxia-dystonia, and in particular epilepsy, given its known imbalance of excitatory and inhibitory neuronal activity (*45*). In all, these results indicate that Pgdb5 is required for the function of specific excitatory and inhibitory cortical neurons.

Although somatic genetic mosaicism has been documented extensively in diverse tissues, and somatic genetic diversification during neuronal development was originally proposed more than 50 years ago (*10*), the existence of physiologic somatic DNA rearrangements in vertebrate brain development has not been proven so far. Here, we demonstrate that an evolutionarily conserved domesticated DNA transposase-derived PGBD5 is an unanticipated cause of double strand DNA breaks in normal neuronal and mammalian brain development. We provide evidence that PGBD5 is required for normal brain development in humans and mice, where its genetic inactivation constitutes the PGBD5 deficiency syndrome, characterized by developmental delay, intellectual disability, language and motor impairments, seizures, and reductions in corpus callosum and cerebellar size. This function likely requires the nuclease activity of PGBD5, as evident from studies of mice engineered to express an enzymatically impaired Pgbd5 nuclease mutant.

We observe that Pgbd5 is responsible for recurrent somatic DNA breaks in mouse brains, which explains the long-standing observations of the requirement of NHEJ DNA repair for mammalian brain development. Similar to RAG1/2-dependent somatic genetic diversification during normal lymphocyte development, mammalian neuronal development also requires the evolutionarily conserved NHEJ DNA repair factor XRCC5/Ku80, which we now show to be associated with PGBD5-dependent neuronal DNA breakage. While we cannot exclude the formal possibility that PGBD5 induces DNA breaks to promote cortical neuronal death occurring during the same developmental period, we provide evidence that Pgbd5-induced somatic DNA rearrangements affect recurrent neuronal chromosomal loci. In all, this study establishes the human PGBD5 deficiency syndrome and identifies distinct neuronal populations and Pgbd5-dependent gene expression programs required for normal mammalian brain development and function. These findings and diverse engineered mouse models set a foundation for the identification of molecular mechanisms of PGBD5 and its substrates for neuronal genetic diversification and self-organization in brain development.

PGBD5-mediated somatic neuronal DNA rearrangements may offer a genetic mechanism for neuronal group selection and developmental apoptosis, which are known to affect a large subset of cells produced during mammalian neuronal development (*46–48*). Many studies have implicated DNA replication as a cause of somatic genetic brain mosaicism. However, this mechanism does not explain how DNA breaks and repair occur in post-mitotic neurons. The data presented here offer a plausible mechanism by which physiologic somatic DNA rearrangements induced by PGBD5 may contribute to somatic neuronal diversification and cellular selection as progenitor neuroblasts differentiate and exit the cell cycle and migrate from the ventricular zone to the mantle layer, where post-mitotic neurons exhibit NHEJ-dependent DNA damage and apoptosis (*2, 3, 7*). Indeed, independent concurrent study by Gustincich and Sanges and colleagues has also identified Pgbd5 as a cause of developmental neuronal DNA breaks in mice, with Pgbd5 being required for normal brain cortical neuronal migration and differentiation (*49*). Since RAG1/2 targets distinct genomic loci in developing B- and T-lymphocytes, PGBD5 targets may also depend on neuronal differentiation and function, and in the case of the brain cortex, the specific neuronal populations identified in this study (Figs. 6D-G and S31-32). Future studies will be needed to define PGBD5 functions in various brain regions including the hippocampus and medial temporal regions, given the prominent seizure phenotype of PGBD5 deficiency.

While we favor the conclusion that PGBD5 acts directly on DNA (*20, 21, 24*), additional biochemical and structural studies will be needed to define the exact enzymatic mechanisms of PGBD5 cellular activities and their developmental regulatory factors, including the possibility that PGBD5 promotes somatic DNA rearrangements through recruitment of other nucleases and chromatin remodeling factors. It is also possible that PGBD5 has additional nuclease-independent functions in nervous system development, such as those mediated by interactions with chromatin and other cellular factors. Finally, we cannot exclude non-central nervous system contributions to the developmental defects observed in PGBD5-deficient mice and humans, as PGBD5 is likely expressed in other neuronal tissues such as neuroendocrine and peripheral nervous system.

We must emphasize that PGBD5-dependent DNA rearrangements are not solely responsible for the physiologic requirement for DNA damage repair in nervous system function. For example, post-mitotic neurons also require XRCC1-dependent base excision/single-strand break repair due to single-strand DNA breaks induced by developmental cytosine demethylation (*46*). Recent studies have also shown that brain aging may involve additional somatic genetic processes (*47, 48*). While we used amplification-free DNA sequencing, specificity and sensitivity of bulk Illumina sequencing have intrinsic limitations, and further studies using amplification-free single-cell analyses will be needed to establish specific somatic neuronal DNA rearrangements and their functions, as recently shown by mapping recurrent mosaic copy number variation in human neurons (*17*).

Developmentally controlled DNA rearrangements have been discovered in diverse biological processes. For example, in addition to the function of the domesticated DNA transposase RAG1/2 in immunoglobulin receptor gene diversification in vertebrate lymphocytes (*50*) , the Spo11 DNA recombinase initiates recombination in eukaryotic meiosis (*51*), the Kat1 DNA transposase controls the yeast mating switching (*52*), and the PiggyMac DNA transposase mediates somatic DNA elimination during macronucleus development in ciliates (*53*). PGBD5-dependent mammalian neuronal genome rearrangements suggest that other evolutionarily conserved DNA transposases may be domesticated as developmental somatic nucleases. This would provide molecular mechanisms for genetic diversification during physiologic somatic tissue and organ development. In turn, dysregulation of these processes can cause deleterious somatic mutations, leading to disease. In the case of RAG1/2 and PGBD5, their dysregulation causes somatic oncogenic DNA rearrangements in blood cancers and solid tumors affecting children and young adults (*54*). Thus, dysregulation of PGBD5 functions during brain development may also contribute to the somatic DNA rearrangements in specific neurodevelopmental disorders.

## Supporting information

Supplementary Materials for A transposase-derived gene required for human brain development

## Acknowledgments

We thank Andrew Kung, Alejandro Gutierrez, Michael Kharas, Marc Mansour, Anton Henssen, Maria Gil Mir, Hao Zhu, Gabriella Casalena and all our laboratory members for helpful suggestions, and Sandeep Reddy, Qiangqiang Zang, Songhai Shi, Adria Pares-Palacin, Michael G. Ploof, Rodrigo Gularte Merida, Montserrat Puiggros, Jan Korbel, Tony Papenfuss, Guillaume Bourque, Patricia Goerner Potvin, Nicolas Robine, Ronan Chaligne, MSK Brain Tumor Center, Molecular Cytology, Integrated Genomics, Single-Cell Analytics, Bioinformatics, Mouse Genetics, and Animal Imaging core facilities for technical assistance, and Maria Jasin for the gift of *Ku80*-knockout mice. AK is a Scholar of the Leukemia & Lymphoma Society and acknowledges generous support of multiple funders listed below.

## Funding

National Institutes of Health grant R01 CA214812 (AK)

National Institutes of Health grant P30 CA008748 (AK)

St. Baldrick’s Foundation (AK)

Burroughs Wellcome Fund (AK) Rita Allen Foundation (AK)

Pershing Square Sohn Cancer Research Alliance and The G. Harold and Leila Y. Mathers Foundation (AK)

Starr Cancer Consortium (AK)

Plan Nacional, Agencia Estatal, Spain, PID2020-119797RB-I00 (DT)

National Institutes of Health grant R35 CA253126 (RR, JZ)

Canadian Institutes of Health Research (CIHR) grant PJT-190271 (CLK) Compute Canada Resource Allocation Project (WST-164-AB) (CLK)

## Author contributions

Conceptualization: AK, LJZ, MT, MCK

Methodology: AK, LJZ, ERF, MPF, NDS, RLG, SH

Investigation: LJZ, SAC, RLG, MY, ERF, MPF, DC, PD, AN, HM, JZ, PB, CR, TCE, AP, CN, EBP, PC, PD, RKO, HTA, RM, HH, HAC, MNA, GZ, ME, MBH, HB, CJGM, AP, XZ, NDS, MH, RR, DT, YB, CLK, MCK, MT, AK

Visualization: AK, LJZ

Funding acquisition: AK, MCK, MT

Project administration: AK Supervision: AK, LJ

Writing – original draft: AK, LJZ, MT, RLG, MY, HM

Writing – review & editing: All authors

## Data and materials availability

All data are openly available via Zenodo (10.5281/zenodo.13291236), with sequencing data available from the NCBI Sequence Read Archive (PRJNA876210) as well as single-nucleus RNA/ATAC-seq (GSE272642). Genetically engineered mouse strains are available from the Jackson Laboratory (*Pgbd5^fl/fl^*, *Pgbd5^D236A^*, and *Pgbd5^3xFlag-HA-P2A-eGFP^* stock numbers 037535, 038881, and 039713, respectively).

## Supplementary Materials

Materials and Methods

Supplementary Text

Figs. S1 to S32

Tables S1 to S9

References (*01–80*)

