## Supplementary Materials for A transposase-derived gene required for human brain development for "A transposase-derived gene required for human brain development"

[1]Molecular Pharmacology Program, Sloan Kettering Institute, Memorial Sloan Kettering Cancer Center, New York, NY, 10021; [2]Tow Center for Developmental Oncology, Department of Pediatrics, Memorial Sloan Kettering Cancer Center; New York, United States, 10021; [3]Pediatric Movement Disorders Program, Barrow Neurological Institute, Phoenix Children's Hospital and Departments of Child Health, Neurology, Genetics and Cellular & Molecular Medicine, Phoenix, AZ;[4] Department of Human Genetics, McGill University, Montreal, Quebec, Canada; [5]Barcelona Supercomputing Center (BSC), Barcelona, Spain, 08034; [6]Department of Pharmacology, Weill Cornell Medical College, New York, NY, 10021; [7]Program for Mathematical Genomics, Departments of Systems Biology and Biomedical Informatics, Columbia University, New York, NY; [8] Molecular Biology Program, Sloan Kettering Institute, Memorial Sloan Kettering Cancer Center, New York, NY, 10021; [9] Programs in Biochemistry, Cell, and Molecular Biology, Weill Cornell Graduate School of Medical Sciences, New York, NY 10065. [10]The Australian e-Health Research Centre, CSIRO, Brisbane, Australia; [11]Assistance Publique-Hôpitaux de Paris, Département de Génétique, Hôpital Pitié-Salpêtrière, Paris, France; [12]Research Group on Multimodal Analysis of Brain Function, University of Picardie Jules Verne, France; [13]Pediatric Neurophysiology Unit, Amiens Picardie University Hospital, France; [14]Phoenix Children's Hospital, Phoenix, Arizona; [15]Hacettepe University, Faculty of Medicine & Institute of Child Health, Department of Pediatric Metabolism, Ankara, Turkey; [16]Department of Neuromuscular Diseases, UCL Queen Square Institute of Neurology, London, United Kingdom; [17]Department of Pediatric Medicine, The Children's Hospital, University of Child Health Sciences, Lahore, Pakistan; [18]CENTOGENE GmbH, Rostock, Germany; [19]LR18SP04, Department of Child and Adolescent Neurology, National Institute Mongi Ben Hmida of Neurology, University of Tunis El Manar, Tunis, Tunisia; [20]Graduate Program in Genetics, University of Arizona, Tucson, AZ, 85721; [21] Department of Anesthesiology and Critical Care, Perelman School of Medicine, University of Pennsylvania; [22]Department of Physics and Computer Science, Xavier University of Louisiana, New Orleans, LA; [23]Physiologie de la reproduction et des comportements, UMR INRAe 0085 CNRS7247, Centre INRAE Val de Loire, France; [24]Institució Catalana de Recerca I Estudis Avançats (ICREA), Barcelona, Spain, [25]Lady Davis Institute for Medical Research, Jewish General Hospital, Montreal, Quebec, Canada, [26]Departments of Pediatrics, Pharmacology, and Physiology & Biophysics, Weill Medical College of Cornell University; New York, United States.

**This PDF file includes:**

Materials and Methods
Supplementary Text
Figs. S1 to S32
Tables S1 to S9

### Materials and Methods

<u>Patient recruitment, sequencing and assessment</u>

WES was performed first on Family 2 using whole-exome Illumina sequencing with mean depth of coverage of 40x. A homozygous 1bp deletion resulting in a frameshift in PGBD5 was identified chr1:230492889 (hg19) NM_001258311.2 c.509 GA>G p.(Phe170SerfsTer5). Four additional families were identified through GeneMatcher (*25*). Family 1 had a homozygous nonsense variant identified via WES as chr1:230425880C>A (hg38) NM_001258311.1 c.49G>T p.(Glu17*). Family 3 WES performed by Centogene identified chr1:230425791G>T (NM_001258311.2)c.138C>A (p.Phe170Serfs*5). Family 4 WES performed at UCL Queen Square Genomics identified as chr1:230323510del49 (NM_001258311.2)c.1442_1490del(p.Ile481ThrfsTer2). All variants had gnomAD frequency=0.Variants for Families 1, 2, and 4 were additionally verified by Sanger sequencing (Fig. S1C).

Family 1 Sanger sequencing performed by Integragen on genomic DNA isolated from peripheral blood using forward primer 5' GCTGGGAGGCACTGTGG and reverse primer 5' CAGCCCACGGAGAGTCTG. Family 2 Sanger sequencing performed by Functional Bio on DNA isolated from peripheral blood using Forward primer 5' TGGACAAGCTCTTACGTCCT and reverse primer 5' CCCAGGGATTGGAAATGCAG. Family 4 Sanger sequencing performed at UCL using Forward primer 5' ACCTAACATCCCAAGCCTGA and Reverse primer 3' GGGTCTCTGATGGGCAAGTT.

Clinical phenotypes were provided by collaborating physicians with additional information extracted from patient photos, videos, and MRI images. Facial features assessed by CGM. Videos from Families 1 and 4 reviewed by pediatric movement disorder neurologist (MCK). Informed consent was obtained by the treating clinician under their local IRB protocol.

<u>Gene expression analysis from publicly available data</u>

Expression data from normal tissues was obtained from Genotype-Tissue Expression (GTEx) Project for humans and RIKEN FANTOM5 project for mice.

Single cell brain transcriptomics data was obtained from Allen brain cell atlas (*56, 57*). Analysis was made using modifications on the python scripts provided by the Allen Institute (https://github.com/AllenInstitute/abc_atlas_access).

<u>Magnetic resonance imaging</u>

For patients, multiplanar, multisequence MRI of the brain without contrast: Sagittal T1 FLAIR, axial T2, sagittal T2 CUBE, sagittal FLAIR CUBE, axial FLAIR, axial T2*, axial DWI, coronal T1 IR 3 plane reconstructions. 1-1 acquired at ages 7 and 10 years,1-2 acquired at age 26 months, 5-1 at age 3 years, and 5-2 at 21 months. Neuroimaging findings were also assessed and measured by a board-certified neuroradiologist (PC) (*58*). Corpus callosum (*58*) and cerebellar measurements (*59*) were compared to age and sex matched controls to determine if they fell below a threshold of the 3$^{rd}$ percentile.

Several image processing steps were performed on the T1-weighted brain MRIs (*55*) including registration to the Colin 27 Average Brain Atlas, correcting image bias using the N4 algorithm, followed by intensity normalization and image de-noising, using anisotropic diffusion. Skull stripping was performed using an in-house algorithm developed in Python. In this approach, intradural CSF was identified using thresholding and morphological operations, following which the lateral ventricles were isolated based on their spatial location, allowing the volume of the lateral ventricles (in mL) to be extracted. Cerebral brain tissues (grey matter, white matter) were then isolated based on their MR intensities using the expectation maximization (EM) and Markov random field (MRF) approach. From the cortical grey matter segmentation, three measures of cortical shape were measured (cortical thickness, curvature and sulcal depth) to quantify shape abnormalities. Measures were converted to a z-score (a measurement of standard deviation from normal population) from healthy cortical shape measures measured from the corresponding cortical region compared to the Child Mind Institute Healthy Brain Network cohort of 564 typically developing children (TDC) (Equation 1), based on cortical regions from the Automated Anatomical Labeling (AAL) atlas.

$$(Eqn\ 1) \qquad z - score_{subject} = \frac{(x_{subject} - \mu_{TDC})}{\sigma_{TDC}}$$

One MRI from each of the two patients in Family 1 passed the quality checks for initial MRI data quality and processed segmentations. For each participant, z-scores of grey matter volume, white matter volume, ventricle asymmetry (Equation 2), was extracted.

$$(Eqn\ 2) \qquad Ventricle\ asymmetry = \frac{(vol_{left} - vol_{right})}{(vol_{left} + vol_{right})}$$

For mice, high-resolution Mn-enhanced mouse brain 3D images were acquired on a 9.4-Tesla BioSpec scanner equipped with 114-cm gradient coil (maximum gradient strength 530 mT/m; Bruker Biospin Corp., Billerica, MA). A Bruker ID 4 cm quadrature volume coil was used for RF excitation and detection. Twenty-four hours before imaging, mice were injected intraperitoneally with $MnCl_2$ at a dose of 0.5 mmol/kg. During imaging sessions mice were anesthetized with 1-2% isoflurane gas in air and were positioned prone in the scanner. Animal body temperature was maintained with a circulating warm water bath and animal respiration was monitored with an animal physiological monitoring system (SA Instruments, Inc., Stony Brook, New York). First, T2-weighted scout brain images along with 3 orthogonal orientations were acquired. Then, 3D T1-weighted mouse brain images along the trans-axial orientation were acquired covering the whole brain using FLASH (Fast Low Angle Shot) gradient echo sequence with the following acquisition parameters: repetition time 22 ms, echo time 3.6 ms, and an isotropic spatial resolution of 100 µm and total imaging time of 2 hours.

<u>Plasmid transfection</u>

HEK293T cells were transfected with pD649-IRES-GFP (Empty vector), pD649-3xFLAG-PGBD5-IRES-GFP encoding wildtype *PGBD5*, or plasmids encoding *PGBD5* c.49#G>T, p.(Glu17*) mutation from family 1 or *PGBD5* c.509del, p.(Phe170Serfs*5) mutation from family 2. Briefly, 100,000 HEK293T cells were seeded in a 6-well plate on day 0. The next day cells were transfected with 1 µg of expression plasmids using TransIT-LT1 (MirusBio MIR 2304) following manufacturer's instructions. Sixteen hours post transfection, media was replaced with fresh media. Seventy-two hours post transfection cells, were trypsinized and pelleted for protein extraction.

<u>Western Immunoblotting</u>

Cells were lysed using Covaris sonication in RIPA lysis buffer. Extracted protein was then quantified using a Pierce BCA assay (ThermoFisher Scientific A65453). Briefly, 30 µg of protein extract was separated using NuPAGE 4-12% gradient gel (Invitrogen NP0322PK2) and electrophoresis was performed at 120 V. Proteins were then transferred to a PVDF membrane (Millipore Sigma, IPVH00010) using 20% methanol transfer buffer at 30 V for 1.5 hours at 4 C. The membrane was then blocked with 5% non-fat milk in TBST and then incubated overnight at 4C with one of the following primary antibodies: FLAG antibody, (Millipore Sigma F1804); GFP antibody, (ThermoFisher Scientific MA5-15256), ACTIN antibody (Cell Signaling Technology 3700). The blots were then incubated with HRP-conjugated secondary antibody (Sigma-Aldrich NA931) for 1 hour at room temperature and imaged after incubation with SuperSignal West Atto (ThermoFisher Scientific A38555) using the IQ800 imager (GE).

Animal handling

All animal procedures were performed following the guidelines of the Institutional Animal Care and Use Committee (IACUC) and approved by the Research Animal Resource Center of the Memorial Sloan Kettering Cancer Center. All mice were housed in groups with up to five animals per cage with a 12-hour light/dark cycle, starting at 06:00 am. Food and water were available ad libitum.

Genetic engineering of *Pgbd5*-deficient mice

To generate *Pgbd5*-deficient mice, we used the dual recombinase-mediated cassette exchange strategy (*26*). First, *tm1e* vector targeting exon 4 of mouse *Pgbd5* (Knockout Mouse Programme) was electroporated in C57BL/6 embryonic stem cells, followed by their microinjection into Balb/c blastocysts (Ingenious Targeting Laboratory). Resulting *tm1a* chimeras with a high percentage black coat color were mated to C57BL/6 FLP mice expressing the Flp recombinase to remove the Neo cassette to generate *tm1c Pgbd5*-floxed mice, as confirmed using genotyping with SC2 and SC4 primers (SC2: GAGAGCACCGTTGGTGCATATCAG, SC4: AGAGTATGAGCGGGAGAGGAGCAG). *Pgbd5*-floxed mice were crossed to B6.FVB-Tg(EIIa-cre)C5379Lmgd/J (EIIa-Cre) mice to generate *tm1d Pgbd5* deficient mice, as confirmed by genotyping with SC2 and SC4 primers. *Pgbd5*-deficient mice were backcrossed to C57BL/6J mice for six generations. $Pgbd5^{fl/fl}$ mice are available from the Jackson Laboratory (strain 037535).

Genetic engineering of *Pgbd5* catalytic deficient mice

To generate *Pgbd5*-catalytic deficient mice, we used CRISPR/Cas9-assisted genome editing by zygote microinjection in C57BL/6J (The Jackson Laboratory, ME, USA). Briefly, mice of 3-6 weeks were used as zygote donors. Fertilized eggs were recovered at pronuclear staged from oviducts of copulated females. Zygotes were microinjected in a drop of KSOM medium. Injection cocktails consisted of Cas9 protein (100 ng/µl; PNABio), in vitro synthesized Cas9 mRNA (50 ng/µl; kit, company), crRNA(s) (50 ng/µl each; Integrated DNA Technologies), tracrRNA (200 ng/µl; Integrated DNA Technologies) and donor single-stranded DNA(s) (20 ng/µl each; Integrated DNA Technologies). Cocktails were mixed just before microinjection and kept on ice. Guide RNA sequences were designed by CRISPR tools (IDT, https://www.idtdna.com/site/order/designtool/index/CRISPR_SEQUENCE and ChopChop https://chopchop.cbu.uib.no/). The crRNA and tracrRNA sequences were as follows; crRNA 1 for D/A on Exon 2, sequence (AGCCACTCTGCAGGGAGTCG), crRNA 2 for D/A on Exon 3, sequence (ACATGAACCCCTGATTGACG); tracrRNA. The microinjection setup was composed of an inverted microscope (TE200-U, Nikon), microinjectors (CellTram vario, Eppendorf;

Femtojet, Eppendorf) and manipulators (TransferMan 4r, Eppendorf). Injection cocktails were injected into pronuclei, and after injection, zygotes or overnight-cultured 2-cell stage embryos were surgically implanted into the oviduct of B6/CBA F1 females primed for pseudopregnancy by mating with vasectomized males. Offspring born from the implanted embryos (the founders) were screened for the presence of the insertion of the donor sequences by PCR of genomic DNA extracted from toe clips. PCR primers used are as follows; Primer #1 D/A Exon 2 (AGGCTTCTATAGCAACCGGAGCC); Primer #2 Exon 2 (TGCATGCATGGACCTGCGTGTGG); Primer #3 Exon 3(AGACTCCTGGTCAGAGAAGTCAG); Primer #4 D/A Exon 3 (TGATGAACCCGGTTGAAGAGC). *Pgbd5*$^{D236A}$ catalytic-deficient mice are available from the Jackson Laboratory (Strain 038881).

Genetic engineering of *Pgbd5*$^{3xFlag\text{-}HA\text{-}P2A\text{-}eGFP}$ knock-in mice

*Pgbd5*$^{3xFlag\text{-}HA\text{-}P2A\text{-}eGFP}$ mice were generated by targeting iTL BF1 (C57BL/6 FLP) embryonic stem (ES) cells (inGenious Targeting Laboratory). A targeting vector was designed to comprise a *3xFlag-HA-P2A-eGFP* in the upstream of the stop codon in exon 7 (Fig. S4A). Targeted ES cells were microinjected into the Balb/c blastocysts. Resulting chimeras were mated to C57BL/6N mice to generate germline transgenic mice. The resulting mice were backcrossed to C57BL/6J mice to eliminate the *FLP* allele. Probe sets were designed to detect *7WT* (72bp) and *7MD* (254bp) (Fig. S4A) for genotyping animals (Transnetyx). *Pgbd5*$^{3xFlag\text{-}HA\text{-}P2A\text{-}eGFP}$ knock-in mice are available from the Jackson Laboratory (Strain 039713).

Immunofluorescence analysis of *Pgbd5*$^{3xFlag\text{-}HA\text{-}P2A\text{-}eGFP}$ expression

Under deep anesthesia, 3 months old *Pgbd5*$^{3xFlag\text{-}HA\text{-}P2A\text{-}eGFP}$ homozygous and C57BL/6J wild-type mice were perfused with intracardiac 0.9% saline followed by 4% paraformaldehyde (PFA)/ 0.1M phosphate buffer (PB). Brains were dissected, and further fixed in 4% PFA/0.1M PB overnight, cryoprotected in 30% sucrose, and embedded in OCT. Blocks were sectioned sagittally in 10 micrometer sections using a cryostat (Leica). Sections were stored at -20°C. eGFP signal together with DAPI was imaged using LSM800 confocal microscope (Zeiss). Thereafter, antigen retrieval was done in sodium citrate buffer (10 mM sodium citrate, 0.05% Tween20, pH6.0) for one hour at 99°C. Anti-NeuN rabbit monoclonal antibody (D4G40: Cell Signaling Technology) was used in combination with an Alexa Fluor 647 F(ab')$_2$ fragment donkey anti-rabbit IgG (H+L) (Jackson Immuno Research). Two confocal images for eGFP and NeuN were co-registered using Photoshop (Adobe). For staining glial fibrillary acidic protein (GFAP) and TMEM119, anti-GFAP (GA5; Millipore Sigma) and anti-TMEM119 antibody (28-3; Abcam) were used without antigen retrieval, respectively. Images were captured simultaneously for eGFP and GFAP, or eGFP and TMEM119.

Mouse genotyping

DNA was extracted from tail clips using PureLink Genomic DNA kit (K182000, Invitrogen) following manufacturer's instructions. PCR was performed using SC2 and SC4 primers and Platinum PCR SuperMix High Fidelity (12532016, Invitrogen) reagents, following a protocol of 30 cycles that consists of denaturation at 98C for 10 sec, annealing at 55C for 30 sec and elongation at 72C for 30 sec. After the 30 cycles, an additional elongation step at 72C for 10 min was performed.

Brain weight assessment

Mice were sacrificed following IACUC specifications at 60 days old. After, brain (from olfactory bulbs to medulla oblongata was extracted and weighted on a precision balance scale.

Behavioral studies

All tests were conducted using 8–12 week-old mice. During all behavioral tests, the investigators who performed the tests were blinded to experimental genotypes. Behavioral tests were conducted during the light phase, between 10:00 hours and 16:00 hours. On test days, animals were transported to the dimly illuminated behavioral laboratory and left undisturbed for one hour before testing.

Locomotor assay

The locomotor assay used a 15×21-inch black box, divided into 12 even-sized (4 × 3 inch) rectangles. The time spent and distance traveled in the two rectangles at 150 lux of light illumination were recorded and analyzed using Med Associates Inc automated tracking system, according to manufacturer's instructions (Med Associates Inc). At least 5 mice per group were used in this assay.

Elevated plus maze

The elevated plus maze used a cross maze with 12×2-inch arms with 50 lux of light illumination. Animals were introduced to the middle portion of the maze facing an open arm and allowed to freely explore for 10 minutes. Times spent and distance traveled in the open and closed arms were measured by an automated video-tracking system (Noldus Information Technology). Data were analyzed for distance traveled in the open arm, because it is least confounded by repeated entry. At least 5 mice per group were used in this assay.

Rotarod test

An accelerating rotarod was used with 3 cm cylinders (47650, Ugo Basile). Each experiment included a training phase which included four trials with 15 min between trials. During training, animals were placed on the rod and allowed to run until the speed reached 5 rpm, then the rod was accelerated from 5 to 40 rpm over the course of 300 sec, followed by two trials for 60 seconds at 4 rpm, with 10 minutes between all trials. Mice able to remain on the rod for the training phase were assayed, following 30 minutes of rest in a cage. The final assay did not include the training phase. Only alternate rod positions were used to minimize the confounding effect of neighboring animals. At least 9 mice per group were used in this assay.

Grip strength assay

Mice were assessed for grip strength performance immediately after the rotarod test. Grip strength was measured as tension force using the force gauge (1027SM Grip Strength Meter with Single Sensor, Columbus Instruments). To assess forelimb grip strength measurement, mice were held by the base of their tails over the top of the grid so that only front paws were able to grip the grid platform T-bar. With the torso in a horizontal position, mice were pulled back steadily until the grip was released, while measuring the grip force. Grip strength measurements for both sets of limbs were performed similarly, with the torso held parallel to the grid for forelimb and hindlimb measurements. Measurements were performed 5 times with 5 min resting periods. At least 9 mice per group were used in this assay.

Seizure assessment

For seizure analysis, we utilized the Racine scale, as described previously (*60*). Briefly, mice were manually suspended by their tails while rotated 10 times for 5 seconds, after which they were placed individually into empty cages for observation. Each mouse was observed for 2 minutes.

<u>Waterbath tail immersion</u>

Using a water bath heated at 47°C, 1 cm of the tail tip was submerged and time to tail flick was measured. Five mice per group were used in this assay.

<u>Tissue perfusion and fixation</u>

Mice were anesthetized using ketamine and xylazine, followed by midline sternotomy. Upon cannulation of the left heart ventricle and connection to the Masterflex C/L peristaltic pump (Antylia Scientific), mice were perfused for two minutes with PBS, followed by six minutes with 4% (w/v) paraformaldehyde in PBS. Dissected tissues were fixed in 4% methanol-stabilized paraformaldehyde (15714-S, Electron Microscopy Sciences) in PBS for 16 hours at $4^0$C. Upon anesthesia of pregnant dams and externalization of the uterus, embryos were dissected and immobilized on 6 cm plates using pin needles (Silicone Dissecting Pad Kit 501986, World Precision Instruments). Embryos were perfused with 0.5 ml of PBS, followed by 1 ml of 4% methanol-stabilized paraformaldehyde (15714-S, Electron Microscopy Sciences) in PBS. Fixed tissues were washed in deionized water twice and stored in 70% aqueous ethanol followed by paraffin embedding or incubated in a 30% (w/v) sucrose in PBS and frozen in Clear O.C.T. compound (4585, Thermo Fisher Scientific).

<u>Immunofluorescence and Histology</u>

Fixed specimens (at least 3 per group) were sectioned using a microtome (Leica RM2155) in 5 µm sections. Slides were deparaffinized and rehydrated using equal volumes of xylene (100%) 3 times followed by a graded alcohol series (100-50%), followed by washing with deionized water. Antigen retrieval was performed using two methods: for PGBD5 antibody Proteinase K at 10U/ml, or the other antibodies citric acid-based solution (H-3300, Vector) in a 95°C water bath for 30 min, followed by 5 min immersion in room temperature solution. Sections were then washed twice with PBS (phosphate buffer saline 0.1M, pH 7.5) and once with PBST (PBS with 0.25% Triton X-100). Slices were subsequently blocked for 30 min at room temperature (RT) using blocking solution: 10% normal donkey serum (NSD, 017-000-121, Jackson ImmunoResearch Laboratories), 1% bovine serum albumin (BSA, A2153, Sigma-Aldrich) in PBST. Primary antibodies (Table S7) were incubated for 16 hours at $4^0$C in blocking solution. Slides were washed 4 times for 10 min using PBST at room temperature. Secondary antibodies (Table S7) were incubated for an hour in the dark at room temperature in 1% NSD and 1% BSA in PBST. Slides were washed 4 times in the dark using PBST for 10 min. Slides were then stained with Hoescht-33342 (H1399, Invitrogen) for 5 min. Slides were washed one last time for 5 min in PBS and coverslips were mounted at medium light using ProLong Diamond antifade mountant (P36962, ThermoFisher). Sagittal sections of cerebellum were stained with hematoxylin and eosin.

<u>Terminal deoxynuleotidyl transferase dUTP nick end labeling (TUNEL) assay</u>

Same deparaffinization and rehydration protocol from immunofluorescence was used. Click-iT TUNEL assay was performed following manufacturer's instructions (C10617, Invitrogen).

<u>RNA *In situ* hybridization (RNA-ISH)</u>

Paraffin-embedded tissue sections were cut at 5 μm and kept at 4°C. Samples were loaded into Leica Bond RX, baked for 30 mins. at 60°C, dewaxed with Bond Dewax Solution (Leica, AR9222), and pretreated with EDTA-based epitope retrieval ER2 solution (Leica, AR9640) for 15 mins. at 95°C (no proteolytic retrieval). For fluorescent *in situ* hybridization (FISH), the probe for mouse *Pgbd5* (Advanced Cell Diagnostics (ACD), Cat# 561948) was hybridized for 2hrs at 42°C. Mouse *PPIB* (ACD, Cat# 313918) and *dapB* (ACD, Cat# 312038) probes were used as positive and negative controls, respectively. The hybridized probes were detected using RNAscope 2.5 LS Reagent Kit – Brown (ACD, Cat# 322100) according to manufacturer's instructions with some modifications (DAB application was omitted and replaced with Fluorescent Alexa 488/Tyramide (Invitrogen, B40953bak) for 20 mins. at RT). FISH slides were loaded in Leica Bond RX for IF staining. Samples were pretreated with EDTA-based epitope retrieval ER2 solution (Leica, AR9640) for 20 mins. at 95°C. Mouse monoclonal antibody Tuj1 (Biolegend, MIMS-435P) was incubated for 1h at RT. Samples were then incubated with Leica Bond Post-Primary reagent (Rabbit anti-mouse linker) (included in Polymer Refine Detection Kit (Leica, DS9800)) for 8 min, followed by incubation with Leica Bond Polymer (anti-rabbit HRP) (included in Polymer Refine Detection Kit (Leica, DS9800)) for another 8 min. Alexa Fluor tyramide signal amplification reagents (CF® 594/ tyramide conjugates (Biotium 92174) were used for signal visualization. After staining, slides were washed in PBS and incubated in 5 μg/ml 4',6-diamidino-2-phenylindole (DAPI) (Sigma Aldrich) in PBS (Sigma Aldrich) for 5 min, rinsed in PBS, and mounted in Mowiol 4–88 (Calbiochem). Slides were kept overnight at -20°C before imaging.

For RNA *in situ* hybridization, BaseScope hybridization probes specific for Pgbd5 exon 4 were generated as per manufacturer's instructions (ACD, catalog 1181898-C1). Upon cardiac perfusion and fixation in 4% PFA/0.1M PB overnight, dissected brains were washed twice with 30% sucrose/PBS and incubated overnight at 4°C. After cryopreservation in sucrose, brains were embedded in OCT (Thermo), and blocks were sectioned sagittally in 10 micrometers sections using a rotary microtome cryostat (Leica). Sections were stored at −80°C. Cryosections were baked for one hour at 60°C and fixed in 4% PFA for 15 min followed by washing in PBS. After dehydration, epitope retrieval treatment with ER2 for 5 min at 95°C and subsequent Protease III treatment for 15 min at 40°C were performed. The probe set was hybridized for two hours at 42°C. Signal amplification steps were performed according to the manufacturer's protocol. Fast Red (Leica Bond Polymer Refine Red Detection kit DS9390) was used as the chromogen. Hematoxylin was used as a counterstain. Mouse Ppib (ACD, catalogue 701078) and Bacillus subtilis dapB (ACD, catalog 701018) probe sets were used as positive and negative controls, respectively.

Flow cytometry analysis

Thymus and spleen were collected using standard dissection. Lymphocytes were extracted by mechanical dissociation followed by erythrocyte lysis using RBC lysis buffer (Biolegend cat.420301). Cells were filtered using a 70-μm mesh (Fisherbrand cat.22-363-528). Cells were blocked on ice for 30 min using mouse IgG isotype control antibodies (Invitrogen cat. 10400C) at 1:1000 dilution in 2.5% FBS in PBS. Cells were aliquoted at $1x10^6$ cells/ml. Antibodies were added at the specified concentrations (Table S7) and incubated for 30 min on ice protected from light. Nuclear staining was done using DAPI. Flow cytometry analysis was performed using BD LSRFortessa cell analyzer and FCS Express 6 software.

Microscopy and image acquisition

Confocal imaging was performed using a Leica TCS SP5 microscope with plan-apochromat 20x/0.75NA objective lens with 4x digital zoom. Images were quantified using blinded observers. Epifluorescence and bright field images were acquired using 20x/0.8NA objective from the Pannoramic P250 Flash microscope (3Dhistech).

Whole-genome sequencing

Matched litter mates (3 per group) were dissected, then, tissue was extracted, flash frozen and kept at -80°C till further processing. For adults, olfactory bulb, hippocampus, cerebellum and peripheral mononuclear cells (PBMC) were obtained. For embryos, forebrain, midbrain, hindbrain and liver were collected. After PicoGreen quantification and quality control by Agilent BioAnalyzer, 137-500 ng of genomic DNA was sheared using a LE220-plus focused ultrasonicator (Covaris catalog # 500569) and sequencing libraries were prepared using the KAPA Hyper Prep Kit (Kapa Biosystems KK8504) kit with the following modifications to the manufacturer's instructions. After adapter ligation, libraries were subjected to 0.5X size selection using AMPure XP beads (Beckman Coulter catalog # A63882). DNA quantity was estimated by real-time PCR using sequencing adapter primers, and libraries were mixed at equimolar ratios for sequencing. Samples were sequenced using NovaSeq 6000 in a 150bp/150bp paired-end mode, with the NovaSeq 6000 SBS v1 kit and S4 flow cells (Illumina).

Sequenced data alignment and processing

All FASTQ files corresponding to the same sample were merged. Sequencing reads were aligned to the mouse reference genome (GRCm38/mm10) downloaded from (https://genome.ucsc.edu) using bwa-mem algorithm (*61*). We used bammarkduplicates to mark the duplicated reads. For the alignment summary metrics, we used Alfred v0.1.16 (*62*).

Variant calling

Variant calling was performed using Pindel version 2.2.3 (*37, 63*) and Delly2 (*35*) with the following parameters: -c 0.05, -a 0.05 and -m 15. Following the merging of results, duplicate calls were filtered using a similarity window of 300 bp, keeping those with the highest detection quality, variant allele frequency. Only variants with features exceeding default PASS quality filter were included for analysis.

Identification and analysis of somatically rearranged genes

For the detection of genes affected by observed variants, we used the BEDTOOLS package (*64*) and the annotation of NCBI genes for mouse downloaded from (https://genome.ucsc.edu). To study the effect of the different subsets of variants, we used the ENSEMBL Variant Effect Predictor (https://www.ensembl.org/Tools/VEP) (*65*) with default settings.

Identification and analysis of somatic genome rearrangement regions

For the detection of genomic intervals, the regions have been created dynamically through the list of mutations contained in each group, with a static window size of 3 Mb, a minimum of two mutations/window, and excluding groups of mutations that come from larger windows using Intersect in bedtools (*64*).

Recurrence analysis of polyclonal somatic DNA rearrangements

A series of R-scripts were written to first parse and filter the VCF files for events, which are available from https://github.com/kentsisresearchgroup/P5BrainReorg. The filtered events were then checked for overlaps and clusters of mutually overlapping groups were output. For the events called by the Delly2 algorithm, the overlaps were computed in two ways. Method 1 looked at the actual genomic intervals and two events were defined to overlap if either the length of both was less than 100,000 bp or the percent overlap was greater than 0.1%, where the percent overlap was computed as the length of the overlap divided by the average size of the two events. Method 2 examined the overlap between the breakpoints of the somatic variants. The ends were defined as a region of 701 bp centered on each end of the event. It was these edge regions that were then intersected for overlaps this time using the rule that the percent overlap was greater than 1%.

Once all overlapping events were identified, clusters were determined by looking for groups of events that all overlapped with each other. That is each element in the cluster had to overlap with every other element, as defined as a maximal clique in graph theory. We used the R package *igraph* to both construct the overlap graph and find all maximal cliques in the graph. We then filtered for sets with 3 or more events and output the table of clusters. To examine how the clusters varied on the filtering parameters this entire procedure was repeated multiple times each time adjusting one of the following parameters: normalized total read counts (RC/EventSize), high-quality variant junction reads (RV), variant allele frequency (RV/[RR+RV], where RR is the high-quality reference junction reads).

For SNV and small insertions/deletions called by Mutect2, a similar procedure was used to identify overlaps for the precise same event in multiple samples. Filtering was done on the following variables: allele depth, variant allele frequency and total depth. Since Mutect2 does not report total depth per sample, we computed an estimate of total depth by taking the maximum value of the following three estimates: RD/(1-AF), AD/AF and AD+RD, where RD is the depth of the reference reads, AD is the depth of the alternate reads and AF was Mutect2's reported allele frequency. We also computed a consistent variant allele frequency of AD/(AD+RD).

To manually inspect that these rearrangements we extracted split reads and discordant pair-end reads using SAMBLASTER (*66*). We then used SparK with default -pr -cf -o -gl , and mm10 reference gene gtf file for -gtf and -gs "yes" parameters to visualize and plot the split reads found in the regions of interest (*67*).

<u>Single-nucleus RNA and ATAC sequencing</u>

Brains from 21 days old *Pgbd5*$^{wt/wt}$ (n=3) and *Pgbd5*$^{-/-}$ (n=3) littermates were extracted. Motor cortices were dissected, flash frozen and stored at –80 C. Nucleus extraction was performed according to the protocol from Masilionis et al. (*68*) using the Singlulator100 (S2 Genomics). Extracted nuclei were stained with 7-aminoactinomycin D (Invitrogen) and FAC-sorted for further nuclei purification. Ten thousand nuclei were targeted for library construction using ChromiumNextGEM Multiome ATAC + Gene Expression kit (10x Genomics).

Cell Ranger ARC v2.0.0 (10X Genomics) was used ('count' option with default parameters) to filter and align raw reads, identify transposase cut sites, detect accessible chromatin peaks, call cells and generate raw count matrices for scMultiome samples. Alignment was performed using the mm10 reference genome build coupled with the Ensembl 98 gene annotation. Reads that mapped to the intronic regions were excluded for the RNA modality.

Quality control (QC) and subsequent data processing steps were performed using Signac v1.3.0(*69*) and Seurat v4.3.0.(*70*) QC metrics for RNA and ATAC modalities were calculated independently but were jointly used to filter cells. A combination of thresholds was established for each sample based on hard cut-offs or on the distribution of each metric within the sample (thresholds specified in Table S5). In the RNA modality, cells were filtered on the number of genes, unique molecular identifiers (UMIs), and mitochondrial content. In the ATAC modality, cells were filtered on the number of peaks detected, transcription start site enrichment, and nucleosome signal.

<u>Cell type annotation in single nuclei RNA data</u>

Annotation of cell types was performed using a consensus approach between seven different reference-based annotation tools, using two references separately. Six machine learning-based prediction tools (ACTINN (*70*), scAnnotate (*71*), SciBet (*72*), SingleR (*73*), scClassify(*74*), and Support Vector Machines) and one model based on maximum Spearman correlation (as previously described (*41*)) were trained on a developmental murine atlas of the forebrain (Jessa et al.(*41*)) or a postnatal murine atlas of the isocortex with high neuronal diversity (Yao et al.(*40*)). Labels predicted by each tool were aggregated into major cell type classes according to the ontology found in Table 6S6, and a consensus label annotation was assigned to each cell using a majority vote approach between tools (with at least two tools agreeing). Predictions based on the developmental atlas (Jessa et al.(*41*)) were used to assign broad cell types, while predictions based on Yao et al.(*40*) were used to annotate neuronal types.

Normalization, dimensionality reduction, and doublet detection in single cell multiome data

For the RNA modality, libraries were scaled to 10,000 UMIs per-cell and log normalized. UMI counts and mitochondrial content were regressed out from normalized gene counts and the residual z-scored gene-wise. Dimensionality reduction was performed using principal-component analysis (PCA) on the top 2,000 most variable features. For the ATAC modality, peaks were called using MACS2 (v2.2.7.1)(*75*) using the CallPeaks function from Signac library with default parameters. ATAC reads were quantified in each peak per cell and a resulting count matrix was generated. Dimensionality reduction was performed using latent semantic indexing (LSI) (*76*).

A weighted nearest neighbor graph was constructed between all cells using the first 30 principal components of the RNA data and the top 6 dimensions of the LSI reduction from the ATAC data with the following default parameters: 20 multimodal neighbors, 200 approximate neighbors and L2 normalization enabled. This weighted nearest neighbor graph was used as input for clustering using a shared nearest-neighbor (SNN) algorithm(*70*) based on the Louvain algorithm on a k-nearest neighbor graph with k = 20 and resolution 0.2.

For each sample, doublets were identified using scDblFinder v1.8.0 (*77*) with the recommended cluster-based approach and with the RNA modality counts.

Joint sample visualization in single nuclei RNA data

For visualizing multiple samples, samples were first merged by genotype, generating a joint sample object for KO and WT separately, as follows. Corresponding RNA libraries were merged without batch correction, followed by scaling, normalization, and dimensionality reduction as described in the previous section except for the regression of variables. The first 30 principal components were used as input for projection into two dimensions (uniform manifold approximation and projection (*78*)) and for clustering (SNN) with k = 20 and resolution 0.5. These genotype joint objects were used to perform post-clustering QC (described in the next section).

After post-clustering QC, all samples were merged in a single joint object (in the same manner as the merging of genotype joint objects) to perform comparison of cell populations, differential gene expression analysis, and differential promoter activity analysis between KO and WT samples.

Post-clustering QC

QC of cell populations and clusters in each genotype joint object was performed to avoid an imbalance in cell populations between KO and WT samples. First, clusters consisting of doublets with a proportion greater than 70% were completely removed. Cells called as doublets were filtered in the remaining clusters.

Next, cell type annotations were inspected at the cluster level as follows. Cluster annotations consisting of a mixture of conflicting cell lineage labels were manually inspected by assessing their gene signatures. Gene signatures were derived by identifying differentially expressed genes in each cluster compared to all other cells using Seurat's FindAllMarkers function (Wilcoxon rank sum test). For each cluster, positive markers were sorted by adjusted p-value, mitochondrial (defined as having gene symbols starting with "mt-") and ribosomal genes (defined as having gene symbols matching "Rps", "Rpl", "Mrps", and "Mrpl") were filtered out, and the top 100 genes were used as a signature. Some clusters were then reannotated as "Striatal Neurons" or "Other Neurons" by assessing the expression of the cluster gene signatures in a whole adult mouse brain atlas(*56*).

Finally, clusters driven by a single sample (non-reproducibility across replicates likely due to sampling issues), were removed only if a similar cell type population was not observed in the opposing genotype integration. Clusters which were manually annotated and retained after post-clustering QC can be observed in tables S8 (WT) and S9 (KO).

Comparison of cell population proportions

To identify significant differences in cell population proportions between different genotypes, a permutation test was applied using the scProportionTest package v0.0.0.9(*79*) with a total of 10,000 permutations. Only cells of known cortical origin (excluding "Striatal Neurons", "Other Neurons", and "No Consensus" cells) were considered for comparison of cell population proportions.

Differential expression in single nuclei RNA data

Differential expression analysis was performed in a per-cluster basis within the joint sample object. Only clusters with greater than 200 cells were considered. First, in each cluster, if the proportion of KO or WT cells was below 40%, random downsampling of cells was performed to balance the number of cells per genotype. Next, a Wilcoxon rank sum test was applied through the FindMarkers function from the Seurat package (logFC.threshold = 0, min.pct = 0.05). Gene ontology pathway analysis was performed on each cluster's DEGs with adjusted p-values < 0.05 using the enrichGO function from the clusterProfiler package v4.0.0(*80*), using all detected genes in the integration as the universe (OrgDb = org.Mm.eg.eb, keyType = "SYMBOL", ont = "BP").

Differential promoter activity in single nuclei ATAC data

Quantification of chromatin accessibility was performed on a per-gene basis by counting fragments that overlapped the gene promoter region, defined as 5-kb bins centered on the transcription start site. Next, differential promoter activity analysis was performed in a per-cluster basis. Using the same clusters as in the differential expression analysis, random downsampling of cells per genotype was performed if necessary due to imbalanced cell numbers. Finally, differential promoter activity between genotypes was calculated using a Wilcoxon rank sum test through the FindMarkers function from the Seurat package (logFC.threshold = 0, min.pct = 0.05).

Statistical methods

Statistical analyses were performed using OriginPro2018 and R. Statistical significance was determined using two-tailed t-tests for continuous pair variables, two-tailed Fisher exact and $\chi^2$ tests for discrete variables, and one and two-way ANOVA with Bonferroni and Tukey's test for multiple comparisons.

#### Supplemental Tables

##### Table S1 (separate file)

See file Table_S1.xlsx

##### Table S2

<u>Dysmorphic facial feature analysis by CGM.</u> + indicates the presence of an abnormal feature, - the absence of the feature, and N/A indicates the feature was not able to be assessed from provided photos in the supplementary clinical information.

| Feature | Family 1 | | Family 2 | | Family 3 | | Family 5 | | #affected/ total |
|---|---|---|---|---|---|---|---|---|---|
| Patient # | 1.1 | 1.2 | 2.1 | 2.2 | 3.1 | 3.2 | 5.1 | 5.2 | |
| Facial asymmetry | - | + | - | NA | - | - | - | + | 2/7 |
| Down slanting palpebral fissures | - | - | - | - | - | - | + | + | 2/8 |
| Telecanthus | + | + | - | NA | + | + | + | + | 6/7 |
| Long Palpebral fissures | + | - | NA | - | - | - | + | + | 3/7 |
| Almond shaped palpebral fissures | - | + | NA | NA | - | - | - | - | 1/6 |
| Sparse lateral eyebrows | + | + | - | - | - | - | - | - | 2/8 |
| Medial eyebrow flare | + | + | - | - | - | - | + | + | 4/8 |
| Posteriorly rotated ears | + | + | NA | + | NA | - | NA | NA | 3/4 |
| Fleshy earlobe | + | + | + | + | + | + | + | + | 8/8 |
| Narrow nasal base | + | + | + | + | - | - | - | - | 4/8 |
| widened nasal tip | + | + | - | - | - | - | + | + | 4/8 |
| low insertion of columella | + | + | + | + | - | - | - | - | 4/8 |
| Long philtrum | - | + | - | - | + | - | - | + | 3/8 |
| Short philtrum | - | - | - | - | - | - | + | - | 1/8 |
| Deep philtrum | - | + | + | + | + | + | + | + | 7/8 |
| Cupid-bow | + | + | + | + | + | + | + | + | 4/8 |
| Thick lower lip | + | + | - | - | - | - | + | + | 4/8 |

| | | | | | | | | | |
|---|---|---|---|---|---|---|---|---|---|
| Down turned corners of the mouth | NA | + | + | + | - | + | + | + | 6/7 |
| Tented upper lip | NA | + | - | - | - | - | - | + | 2/7 |
| Retrognathia | + | + | - | - | NA | - | NA | NA | 2/5 |
| Short chin | + | + | - | - | + | + | + | + | 6/8 |

**Table S3 (separate file)**

See file Table_S3.txt

**Table S4 (separate file)**

See file Table_S4.xlsx

**Table S5 (separate file)**

See file Table_S5.xlsx

**Table S6 (separate file)**

See file Table_S6.xlsx

**Table S7**

| Antibody | Vendor | Catalog number | Concentration |
|---|---|---|---|
| PGBD5 | Abbexa | Abx019151 | 1:500 |
| γH2AX | Abcam | ab11174 | 1:500 |
| TUJ1 | BioLegend | 801202 | 1:500 |
| CD4 | eBioscience | 11-0042-82 | 1:500 |
| CD8 | eBioscience | 25-0081-81 | 1:1000 |
| B220 | BioLegend | 103221 | 1:100 |
| IgM | eBioscience | 12-5790-82 | 1:750 |
| α-mouse-Alexa647 | Invitrogen | A32728 | 1:1000 |
| α-rabbit-Alexa555 | Invitrogen | A21422 | 1:1000 |
| ACTIN | Cell Signaling | 3700 | 1:1000 |
| FLAG | Millipore | F1804 | 1:1000 |
| GFP | Thermo-Fisher | MA5-15256 | 1:1000 |

**Table S8 (separate file)**

See file Table_S8.xlsx

**Table S9 (separate file)**

See file Table_S9.xlsx

1

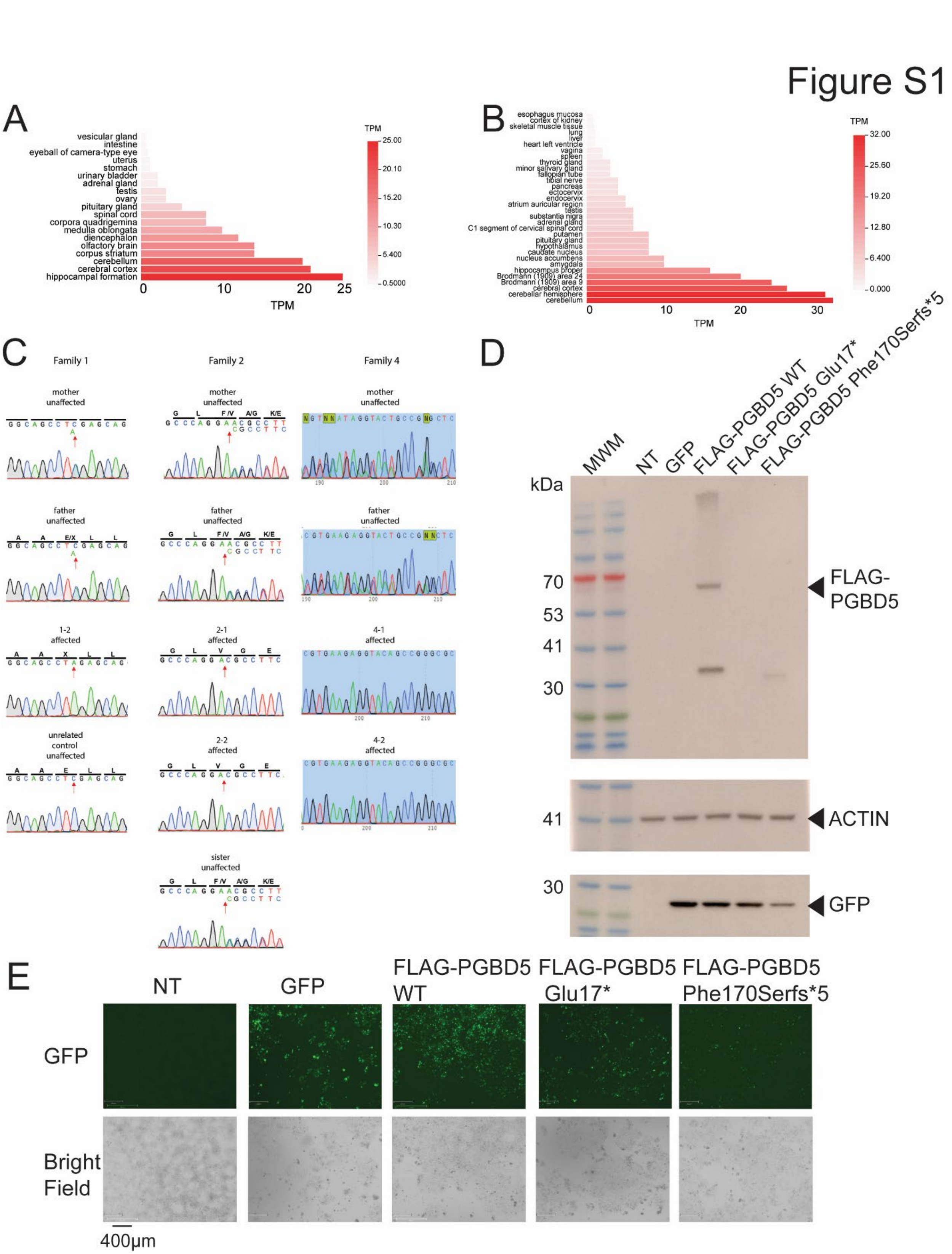
Figure S1
A
vesicular gland
intestine
eyeball of camera-type eye
uterus
stomach
urinary bladder
adrenal gland
testis
ovary
pituitary gland
spinal cord
corpora quadrigemina
medulla oblongata
diencephalon
olfactory brain
corpus striatum
cerebellum
cerebral cortex
hippocampal formation
TPM
B
esophagus mucosa
cortex of kidney
skeletal muscle tissue
lung
liver
heart left ventricle
vagina
spleen
thyroid gland
minor salivary gland
fallopian tube
tibial nerve
pancreas
ectocervix
endocervix
atrium auricular region
testis
substantia nigra
adrenal gland
C1 segment of cervical spinal cord
putamen
pituitary gland
hypothalamus
caudate nucleus
nucleus accumbens
amygdala
hippocampus proper
Brodmann (1909) area 24
Brodmann (1909) area 9
cerebral cortex
cerebellar hemisphere
cerebellum
TPM
C
Family 1
Family 2
Family 4
mother unaffected
father unaffected
1-2 affected
unrelated control unaffected
2-1 affected
2-2 affected
sister unaffected
4-1 affected
4-2 affected
D
MWM
NT
GFP
FLAG-PGBD5 WT
FLAG-PGBD5 Glu17*
FLAG-PGBD5 Phe170Serfs*5
kDa
70
53
41
30
FLAG-PGBD5
ACTIN
GFP
E
NT
GFP
FLAG-PGBD5 WT
FLAG-PGBD5 Glu17*
FLAG-PGBD5 Phe170Serfs*5
GFP
Bright Field
400μm

2
3

**Fig. S1.** ***PGBD5*** **nonsense mutations disrupt protein expression in cells. A-B,** Heatmap showing *PGBD5* gene expression in **A** human and **B** mouse brain regions. Highest expression is observed in the hippocampus, cerebellum and cortex. **C,** Sanger sequencing of unaffected parents and siblings, and affected children showing homozygous mutation in families 1, 2 and 4. **D**, Western immunoblotting of HEK293T cells transfected with wildtype *PGBD5* and specific patient mutations using the pD649-3xFLAG-cDNA-IRES-GFP expression vector; MWM = molecular weight marker, NT = not transfected, GFP = pD649-GFP vector, FLAG-PGBD5 WT = 3XFLAG-PGBD5-IRES-GFP vector, FLAG-PGBD5 Glu17* = PGBD5 WT vector containing *PGBD5* c.49#G>T, p.(Glu17*) mutation from family 1, FLAG-PGBD5 Phe170 Serfs*5 = PGBD5 WT vector containing *PGBD5* c.509del, p.(Phe170Serfs*5) mutation from family 2. Top: blot with FLAG antibody showing expression of PGBD5 protein. Middle: blot of ACTIN as the loading control. Bottom: blot against GFP protein to control for vector expression. **E**, Representative images of GFP expression in HEK293T transfected with the various expression vectors in S1D.

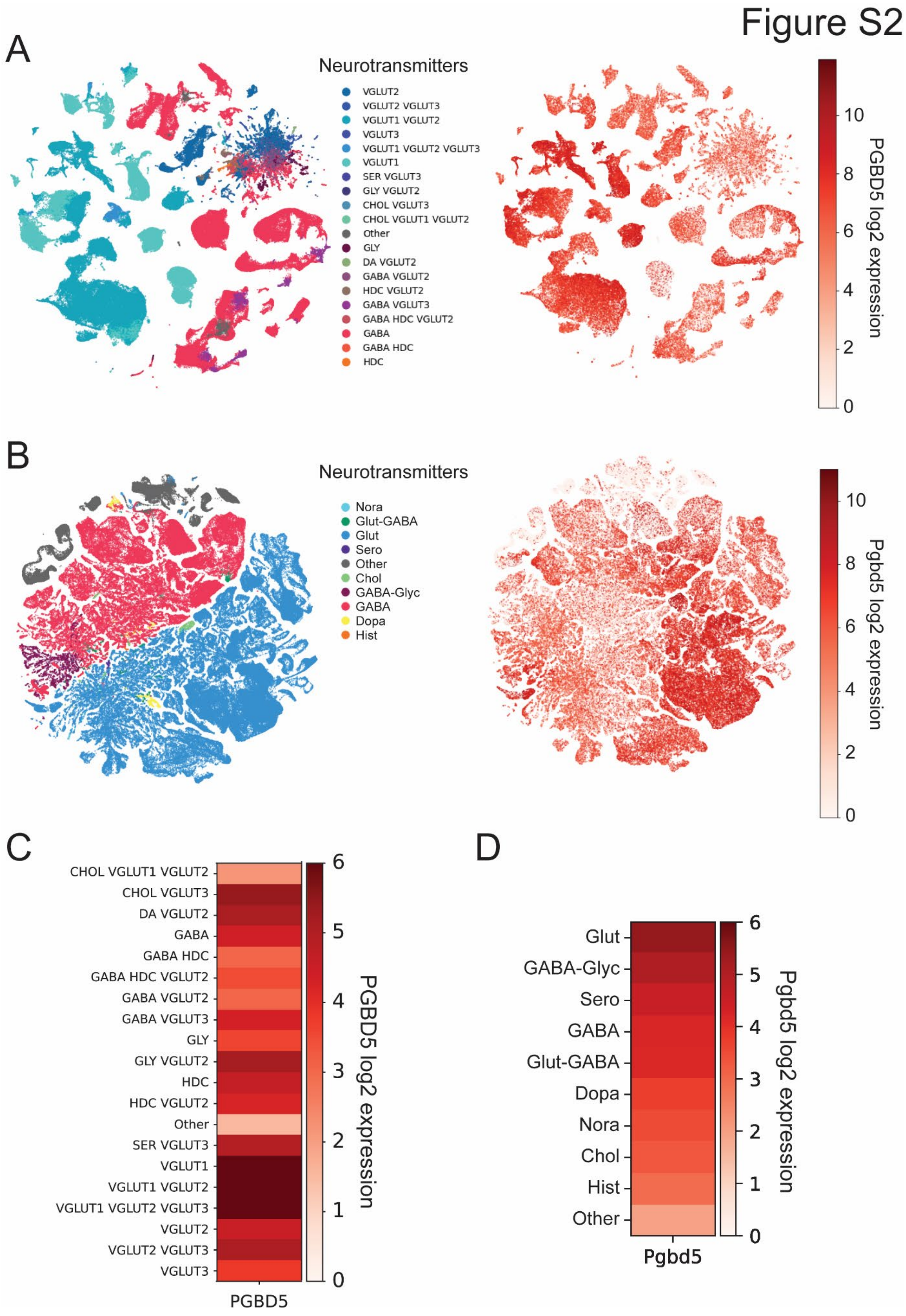
Figure S2
A
Neurotransmitters
VGLUT2
VGLUT2 VGLUT3
VGLUT1 VGLUT2
VGLUT3
VGLUT1 VGLUT2 VGLUT3
VGLUT1
SER VGLUT3
GLY VGLUT2
CHOL VGLUT3
CHOL VGLUT1 VGLUT2
Other
GLY
DA VGLUT2
GABA VGLUT2
HDC VGLUT2
GABA VGLUT3
GABA HDC VGLUT2
GABA
GABA HDC
HDC
PGBD5 log2 expression
B
Neurotransmitters
Nora
Glut-GABA
Glut
Sero
Other
Chol
GABA-Glyc
GABA
Dopa
Hist
Pgbd5 log2 expression
C
PGBD5 log2 expression
PGBD5
D
Glut
GABA-Glyc
Sero
GABA
Glut-GABA
Dopa
Nora
Chol
Hist
Other
Pgbd5 log2 expression
Pgbd5

1

**Fig. S2. Glutamatergic neurons have the highest expression of PGBD5. A**, Uniform manifold approximation and projection (UMAP) plot of single nuclei RNA seq from 3 million single cell transcriptomes spanning the whole adult human brain (*56,57*). Left, cells are colored by neurotransmitter identity using Allen brain atlas classification. Right, cells are colored by *PGBD5* $\log_2$ expression. **B**, UMAP from 4 million single cell transcriptomes spanning the whole adult mouse brain. Left, cells are colored by neurotransmitter identity using Allen brain atlas classification in the ABC project. Right, cells are colored by mouse *Pgbd5* $\log_2$ expression. **C-D**, Heatmaps of *PGBD5* $\log_2$ expression in different neurotransmitter classes (**C**) from human and (**D**) mouse brains. In both instances the glutamatergic neurotransmitter class shows the highest expression of PGBD5.

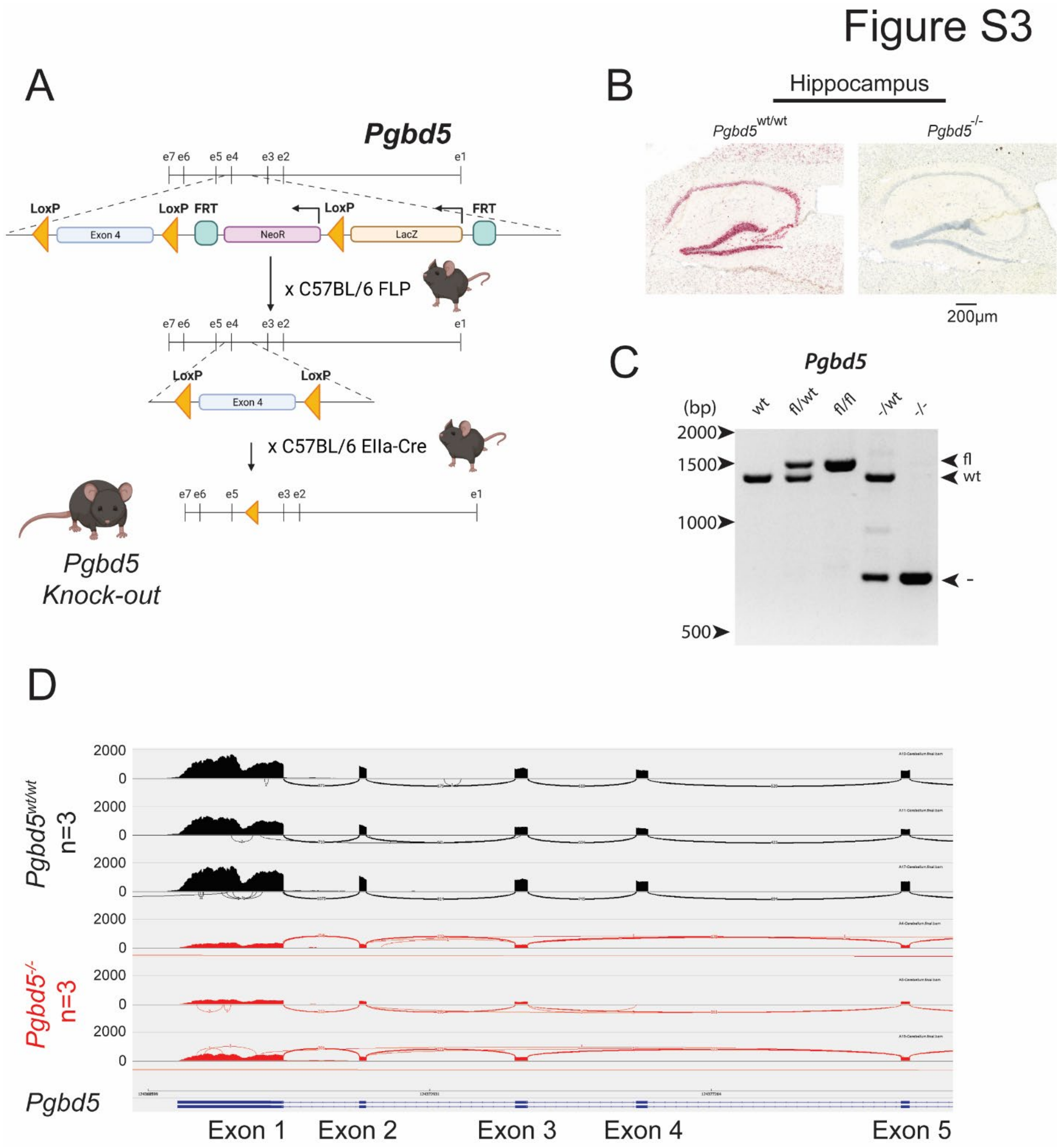


**Fig. S3. *Pgbd5*-deficient mice exhibit specific phenotypic deficits. A**, Schematic of genetically engineered *Pgbd5* locus and breeding strategy to generate *Pgbd5*$^{-/-}$ mice. Triangles and squares indicate *LoxP* and *FRT* sites, respectively, allowing for conditional excision of *NeoR* and *LacZ* targeting cassette, followed by conditional excision of exon 4 of *Pgbd5*. **B**, Representative in situ hybridization micrographs of the hippocampus of 60-day old mice, Pgbd5 (red) and DNA (blue), showing loss of *Pgbd5* expression in *Pgbd5*$^{wt/wt}$ as compared to *Pgbd5*$^{-/-}$ mice. Scale bar = 200 µm. **C**, Gel electrophoresis analysis of genomic PCR of genetically engineered mouse *Pgbd5* locus demonstrating floxed, wildtype, and knock-out alleles. **D**, Sashimi plots of three *Pgbd5*$^{wt/wt}$ and three *Pgbd5*$^{-/-}$ littermate mice showing *Pgbd5* transcripts as measured by RNA-seq of the cerebellum. *Pgbd5*$^{-/-}$ mice have reduced expression of *Pgbd5* lacking exon 4, associated with presumed nonsense-mediated decay.

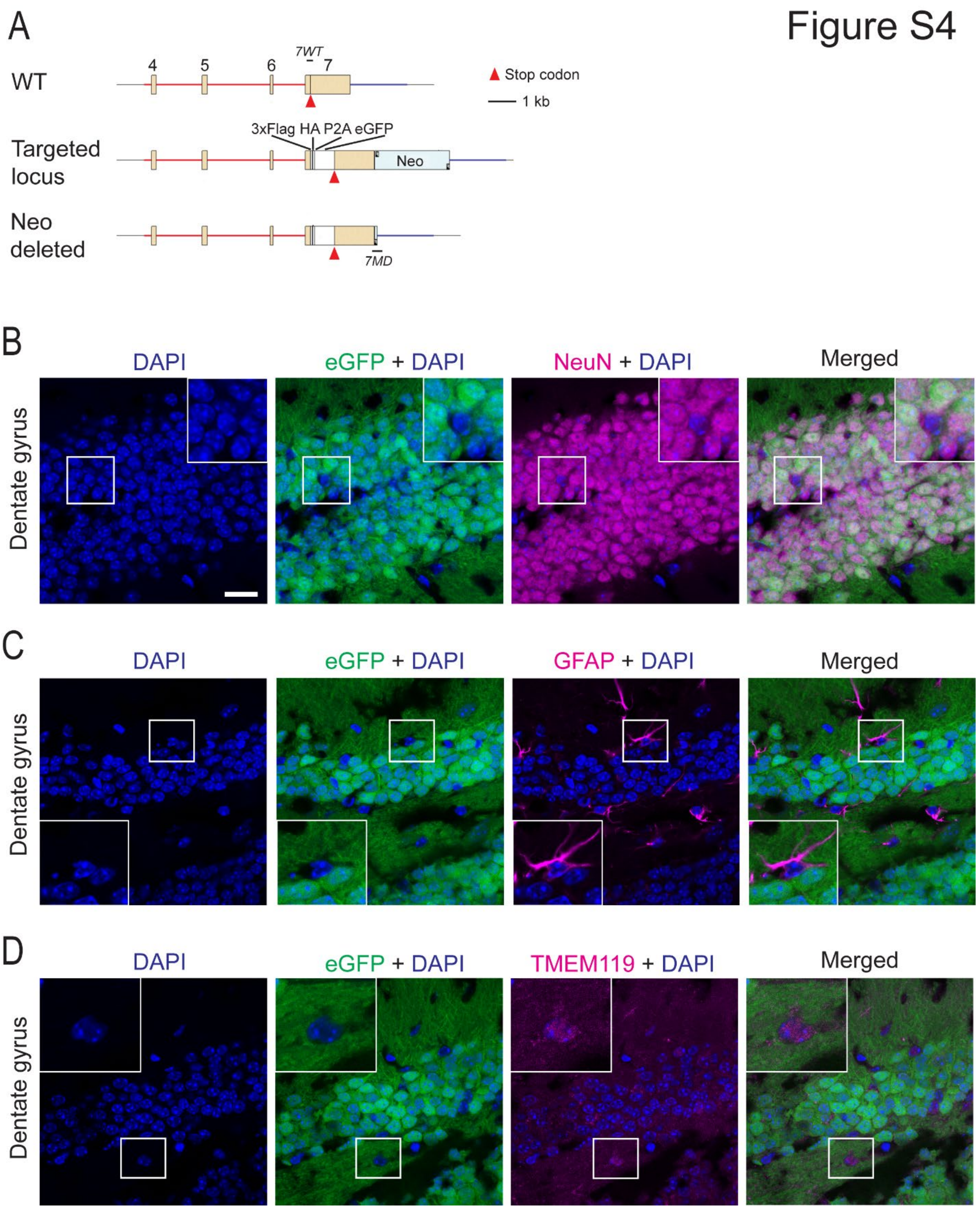

Figure S4
A
WT
Targeted locus
Neo deleted
4
5
6
7
7WT
7MD
3xFlag HA P2A eGFP
Neo
Stop codon
1 kb
B
C
D
DAPI
eGFP + DAPI
NeuN + DAPI
GFAP + DAPI
TMEM119 + DAPI
Merged
Dentate gyrus

**Fig. S4. Pgbd5 is specifically expressed in neurons but not astrocytes or microglia in the mouse dentate gyrus. (A)** Schematic showing the knockin engineering of the endogenous *Pgbd5* locus with $Pgbd5^{3xFLAG\text{-}HA\text{-}P2A\text{-}eGFP}$. **(B-D)** Immunofluorescence of adult mice dentate gyrus showing Pgbd5 expression in neurons (NeuN positive cells) (**B**) but not astrocytes (GFAP positive cells) (**C**) or microglia (TMEM119 positive cells) (**D**). eGFP expression only co-stains with NeuN positive cells indicating that Pgbd5 is exclusively expressed in neurons.

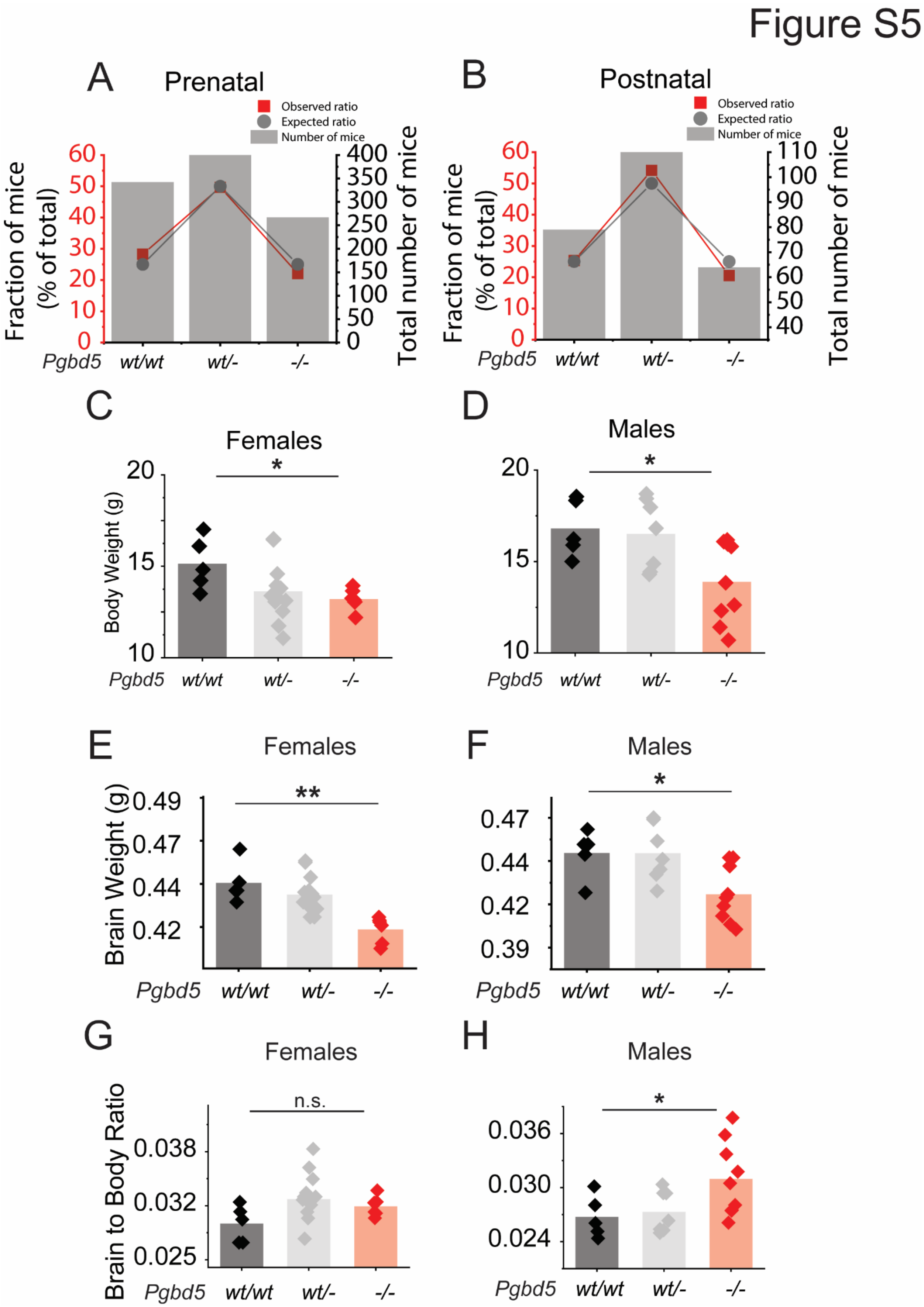
Figure S5
A
Prenatal
Observed ratio
Expected ratio
Number of mice
Fraction of mice (% of total)
Total number of mice
Pgbd5 wt/wt wt/- -/-
B
Postnatal
Observed ratio
Expected ratio
Number of mice
Fraction of mice (% of total)
Total number of mice
Pgbd5 wt/wt wt/- -/-
C
Females
Body Weight (g)
*
Pgbd5 wt/wt wt/- -/-
D
Males
*
Pgbd5 wt/wt wt/- -/-
E
Females
Brain Weight (g)
**
Pgbd5 wt/wt wt/- -/-
F
Males
*
Pgbd5 wt/wt wt/- -/-
G
Females
Brain to Body Ratio
n.s.
Pgbd5 wt/wt wt/- -/-
H
Males
*
Pgbd5 wt/wt wt/- -/-

**Fig. S5, Pgbd5 knock-out mice have reduced body weight. A-B**, Mendelian ratios of *Pgbd5* $^{wt/-}$ in-crosses of prenatal (**A**) and postnatal mice (**B**). **C-D**, Body weights of female (**C**) and male (**D**) mice are significantly reduced in *Pgbd5* $^{-/-}$ male mice (red); Two-way Anova $p$= 7.1E-3 * Tukey's $p$ = 0.098 in female and 8.3E-3 male mice. **E-J**, Total brain weights of female (**E**) and male (**F**) mice are significantly reduced in *Pgbd5* $^{-/-}$ mice (red); Two-way ANOVA $p$=1E-5 ** and * Tukey's $p$ = 1.8E-3 and 4.9E-3, respectively. **G-H**, Brain to body ratios of female (**G**) and male (**H**) mice, show significant change in *Pgbd5* $^{-/-}$ male mice (red); Two-way Anova $p$= 4.37E-2 * Tukey's $p$ = 0.45 in females and $p$= 0.028 in males. *Pgbd5* $^{wt/wt}$ $n$=5, *Pgbd5* $^{wt/-}$ $n$=8 males and $n$=10 females, *Pgbd5* $^{-/-}$ $n$=8 males and $n$=10 females.

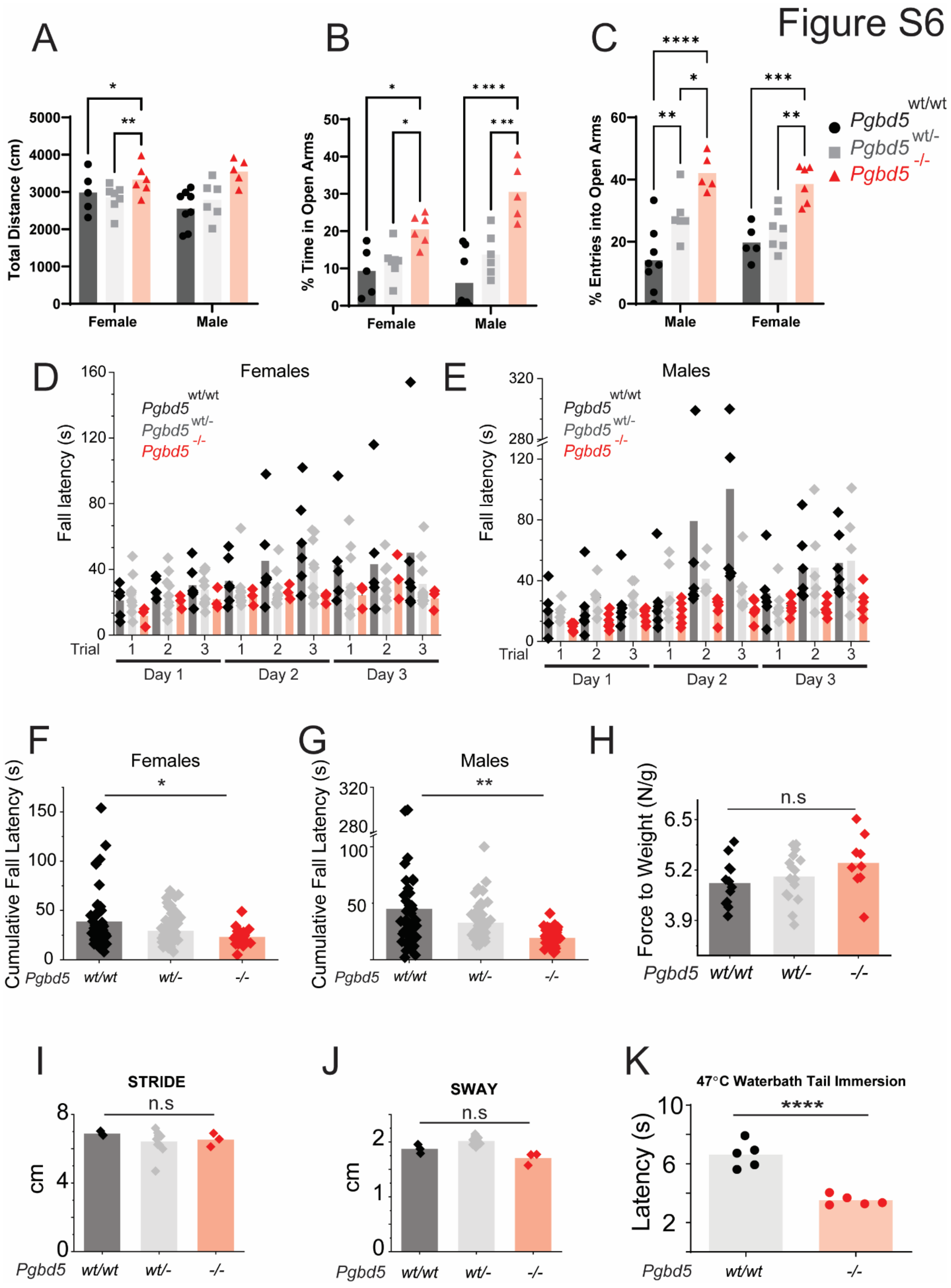
Figure S6
A
Total Distance (cm)
Female
Male
B
% Time in Open Arms
Female
Male
C
% Entries into Open Arms
Male
Female
Pgbd5 wt/wt
Pgbd5 wt/-
Pgbd5 -/-
D
Females
Fall latency (s)
Trial
Day 1
Day 2
Day 3
E
Males
Fall latency (s)
Trial
Day 1
Day 2
Day 3
F
Females
Cumulative Fall Latency (s)
Pgbd5 wt/wt wt/- -/-
G
Males
Cumulative Fall Latency (s)
Pgbd5 wt/wt wt/- -/-
H
Force to Weight (N/g)
n.s
Pgbd5 wt/wt wt/- -/-
I
STRIDE
n.s
cm
Pgbd5 wt/wt wt/- -/-
J
SWAY
n.s
cm
Pgbd5 wt/wt wt/- -/-
K
47°C Waterbath Tail Immersion
Latency (s)
Pgbd5 wt/wt -/-

**Fig. S6, Pgbd5-deficient mice exhibit impaired response to an anxiogenic stimulus and motor learning.** **A-**C, Elevated-plus-maze behavioral analysis demonstrates a significant increase in total distance traveled (**A**), percentage of time in open arms (**B**) and entries into open arms (**C**) of *Pgbd5*$^{-/-}$ mice (red). In total distance traveled (**A)** male *Pgbd5*$^{wt/wt}$ (black) versus *Pgbd5*$^{-/-}$ mice (red), two-way ANOVA *p*=1.9E-3 for phenotype and *p*=0.54 for sex, ** Tukey's test *p* = 1.6E-3; *Pgbd5*$^{-/wt}$ (grey) versus *Pgbd5*$^{wt/wt}$ (black), * Tukey's test *p* = 0.020. In % of time spent in open arms (**B**) male *Pgbd5*$^{wt/wt}$ (black) versus *Pgbd5*$^{-/-}$ mice (red), Two-way *p*=6.6E-3 Tukey's test *p* = 1.0E-4; *Pgbd5*$^{-/wt}$ (grey) versus *Pgbd5*$^{-/-}$(red), *** Tukey's test *p* = 3.0E-4. Females also showed significant reduction in time spent between *Pgbd5*$^{wt/wt}$ (black) and *Pgbd5*$^{-/-}$ mice (red), two-way ANOVA *p*= 4.1E-7 for phenotype and *p*= 0.17 for sex,* Tukey's test *p* = 0.016 and *Pgbd5*$^{-/wt}$ (grey) versus *Pgbd5*$^{-/-}$ (red), * Tukey's test *p* = 0.045. (**C)** All groups exhibited significant differences in time spent in open arms in male mice, *Pgbd5*$^{wt/wt}$ (black) versus *Pgbd5*$^{-/-}$ mice (red), two-way ANOVA *p*=9.9E-8 for phenotype and *p*= 0.75 for sex, **** Tukey's test *p* = 1.0E-4; *Pgbd5*$^{wt/wt}$ (black) versus *Pgbd5*$^{-/wt}$ mice (grey) ** Tukey's test *p* = 3.8E-3; and *Pgbd5*$^{-/wt}$ (grey) versus *Pgbd5*$^{-/-}$ mice (red), * Tukey's test *p* = 0.013. Females also showed significant reduction between *Pgbd5*$^{wt/wt}$ (black) and *Pgbd5*$^{-/-}$ mice (red), *** Tukey's test *p* = 7.0E-4; *Pgbd5*$^{-/wt}$ (grey) versus *Pgbd5*$^{-/-}$ (red), ** Tukey's test *p* = 3.4E-3. *Pgbd5*$^{wt/wt}$ *n*=5, *Pgbd5*$^{wt/-}$ *n*=8 males and *n*=10 females, *Pgbd5*$^{-/-}$ *n*=8 males and *n*=10 females. **D-G**, Rotarod test analysis demonstrates impaired motor learning reflected in the relatively short fall latency from successive day trials. Fall latency in seconds, in females (**D**) and males (**E**). (**F-G)** Cumulative fall latency shows significant motor learning impairment in both female (**F**) (* = One-way ANOVA *p*=9E-4, Tukey's test *p*=1.4E-3) and male (**G**) (** = One-way ANOVA *p*=7E-4, Tukey's test *p*=4E-4) *Pgbd5*$^{-/-}$ mice. **H**, Grip strength assay showing lack of significant effects in Pgbd5-deficient mice (Two-way ANOVA genotype *p* = 0.08 and sex *p*= 0.03, Tukey's test wt vs ko *p*=0.07). **I-J,** Pgbd5-deficient mice show no apparent differences in stride (**I**) and sway in cm (**F**) *Pgbd5*$^{wt/wt}$ *n*=9, *Pgbd5*$^{wt/-}$ *n*=16, *Pgbd5*$^{-/-}$ *n*=12. **K**, Latency in seconds from water bath tail immersion assay showing significant reduction indicating thermal hypersensitivity in *Pgbd5*$^{-/-}$ mice (red); **** t-test *p* = 1.0E-4.

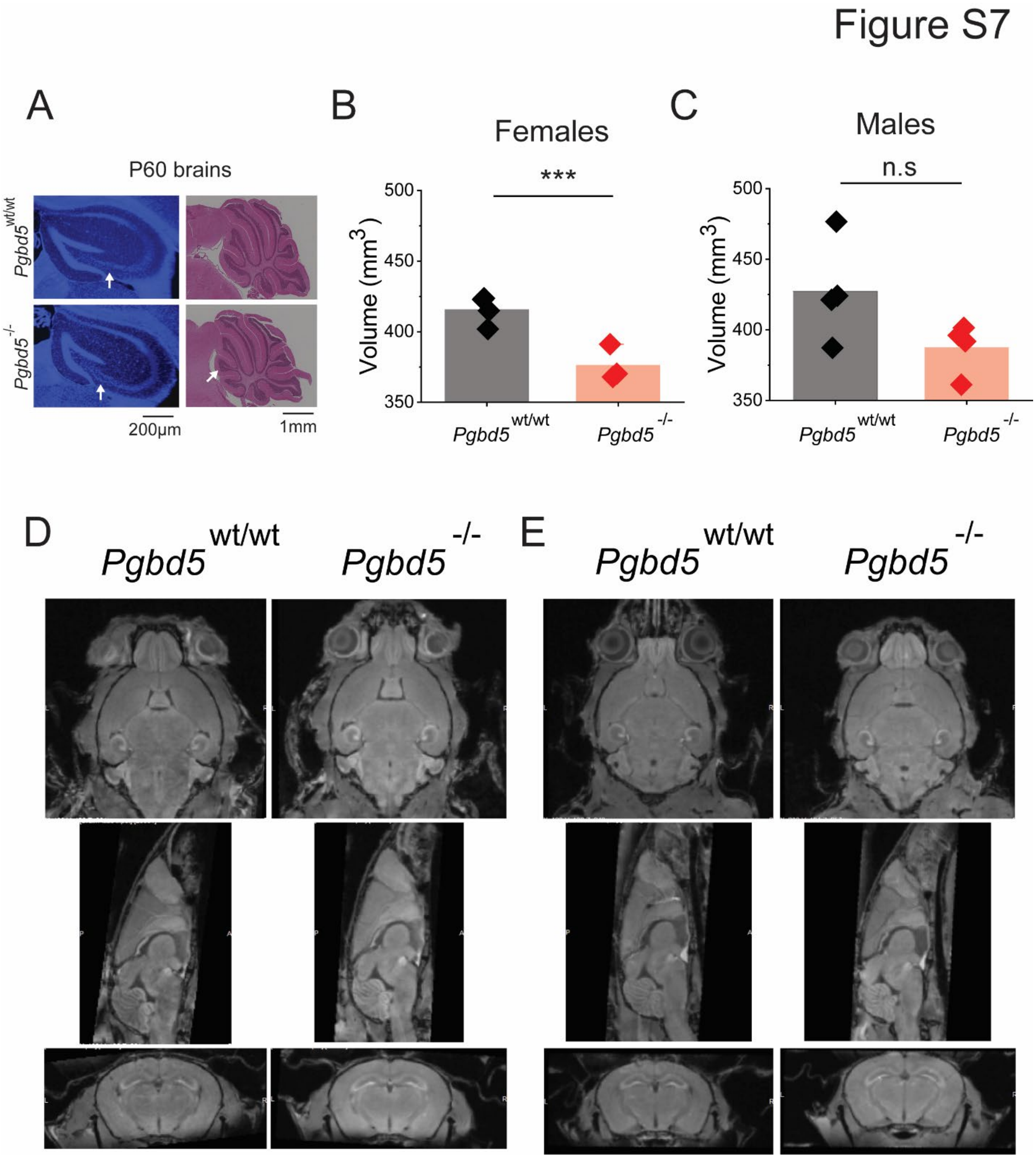


**Fig. S7. *Pgbd5*-deficient mice exhibit reduced brain volumes. A,** Representative fluorescence micrographs of coronal sections of P60 hippocampi *Pgbd5*$^{-/-}$ as compared to *Pgbd5*$^{wt/wt}$ litter mate mice, showing delaminated C3 (arrows). Scale bar = 200 μm and representative H&E micrographs of sagittal sections of P60 cerebellum *Pgbd5*$^{-/-}$ as compared to *Pgbd5*$^{wt/wt}$ litter mate mice, showing extra foliation in *Pgbd5*$^{-/-}$(arrow). Scale bar = 1mm. **B-C**, Cerebral volumes in female (**B**), and male (**C**) mice as measured by microMRI; *** t-test $p$ = 5.9E-3. **D-E**, Representative brain MRI images of male (**D**) and female (**E**) mice.

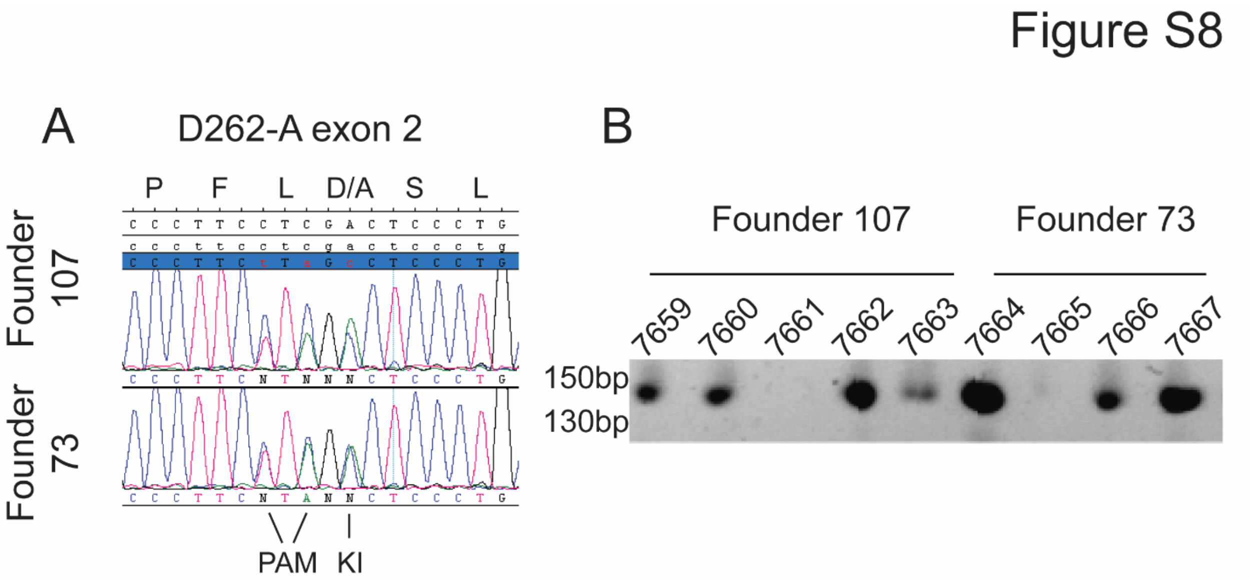


**Fig. S8. Pgbd5 catalytically inactive mice phenocopy full knock-out Pgbd5 mice. A**, Sanger sequence traces of the two founders showing heterozygosity of correct substitution of the PAM site and D262 codon. **B,** enzyme digestion of PCR amplicons of exon 2 D262 region shows correct transmission of both *Pgbd5*$^{ki/ki}$ founders upon backcrossing to C57BL/6.

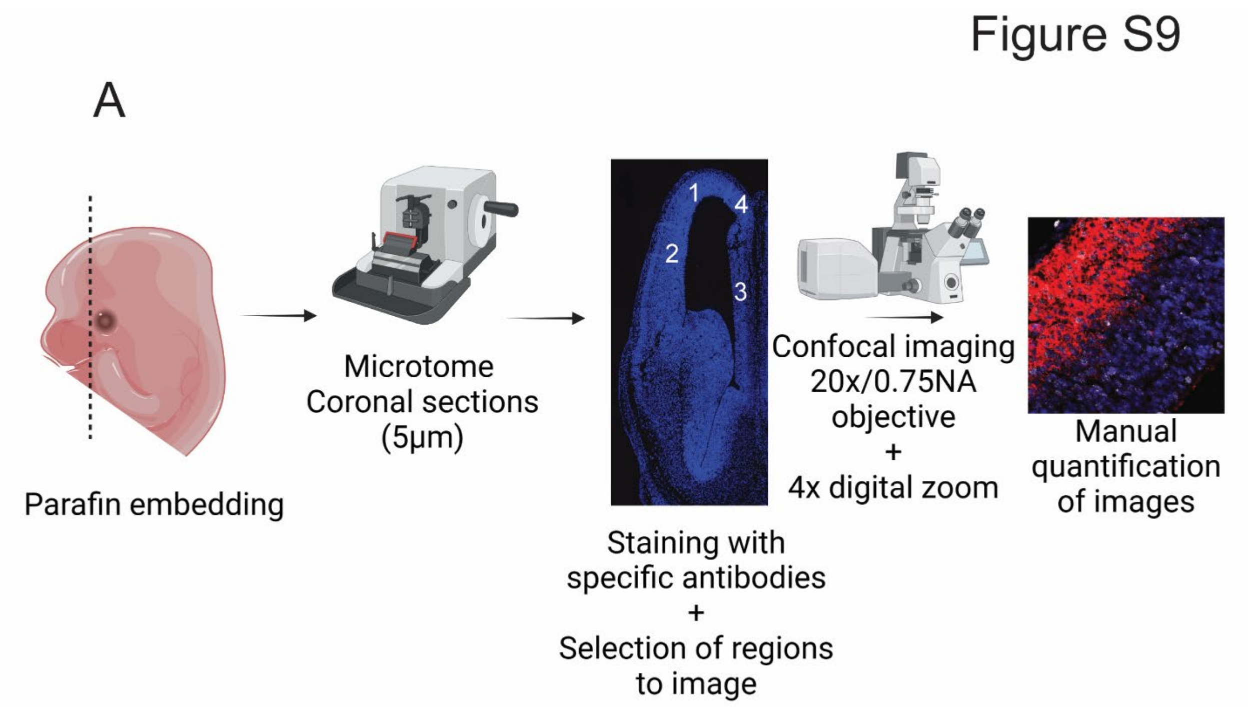


**Fig. S9. Schematics on selection of cortex regions and confocal imaging acquisition. A,** From left to right: Diagram showing E14.5 mouse head and axis for coronal sectioning, Microtome drawing representing how sections were made, a representative image of a E14.5 cortex showing the 4 different cortex-regions selected, the confocal microscope imaging process used in Fig S7-10 and Figure 3-4, representative image of pictures taken by the confocal that were manually quantified.

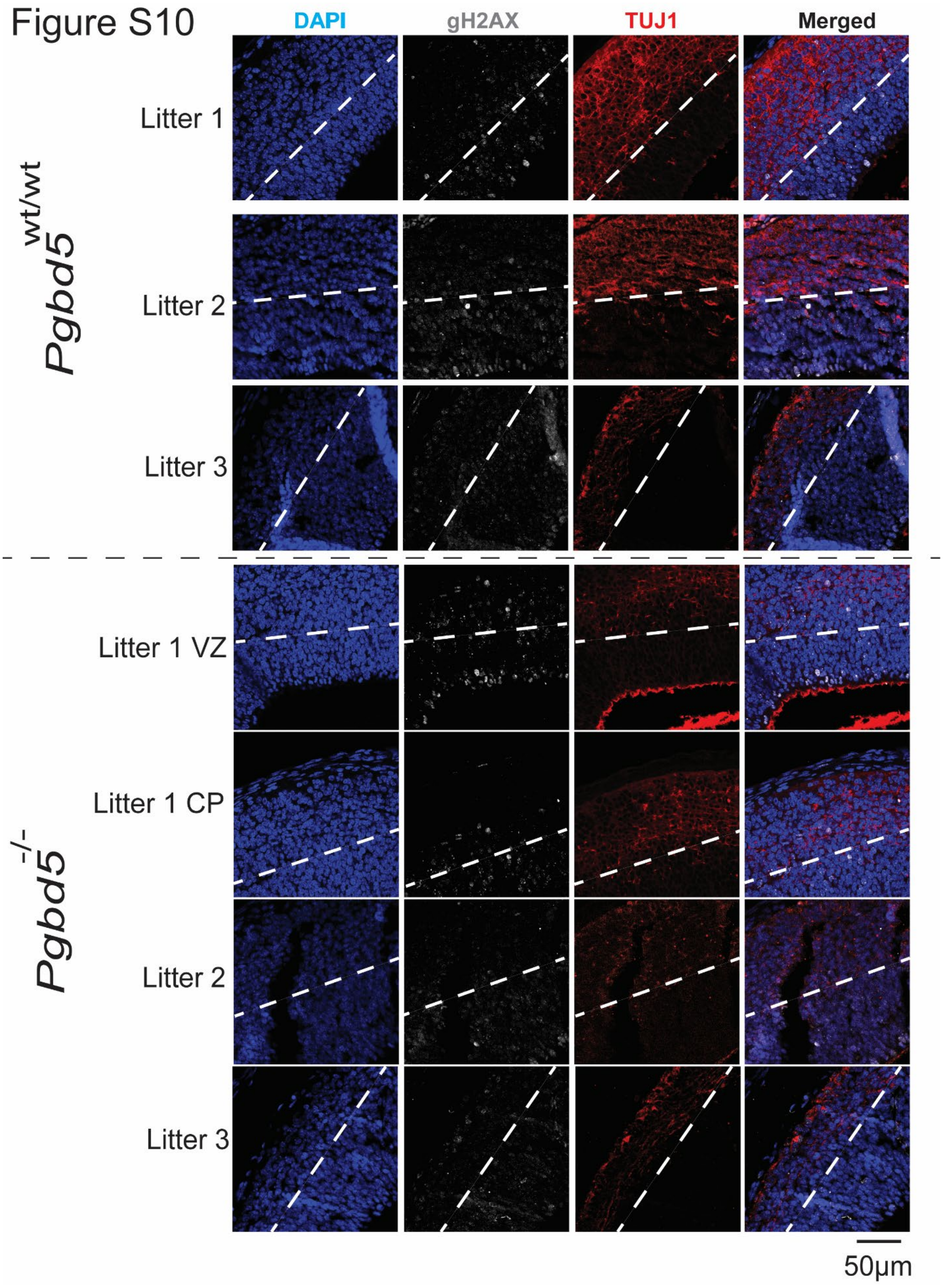

Figure S10
DAPI
gH2AX
TUJ1
Merged
Pgbd5 wt/wt
Litter 1
Litter 2
Litter 3
Pgbd5 -/-
Litter 1 VZ
Litter 1 CP
Litter 2
Litter 3
50μm


1

**Fig. S10. *Pgbd5*-deficient mice exhibit reduced DNA damage in region of interest 1.** Immunofluorescence photographs of brain region 1 from three different independent litters, stained for DNA using DAPI (blue), gH2AX (white), and TUJ1 (red), with specific litter mates indicated (VZ, ventricular zone; CP, cortical plate). Dashed lines indicate separation between ventricular zones and cortical plates. Scale bar = 50 mm.

Figure S11

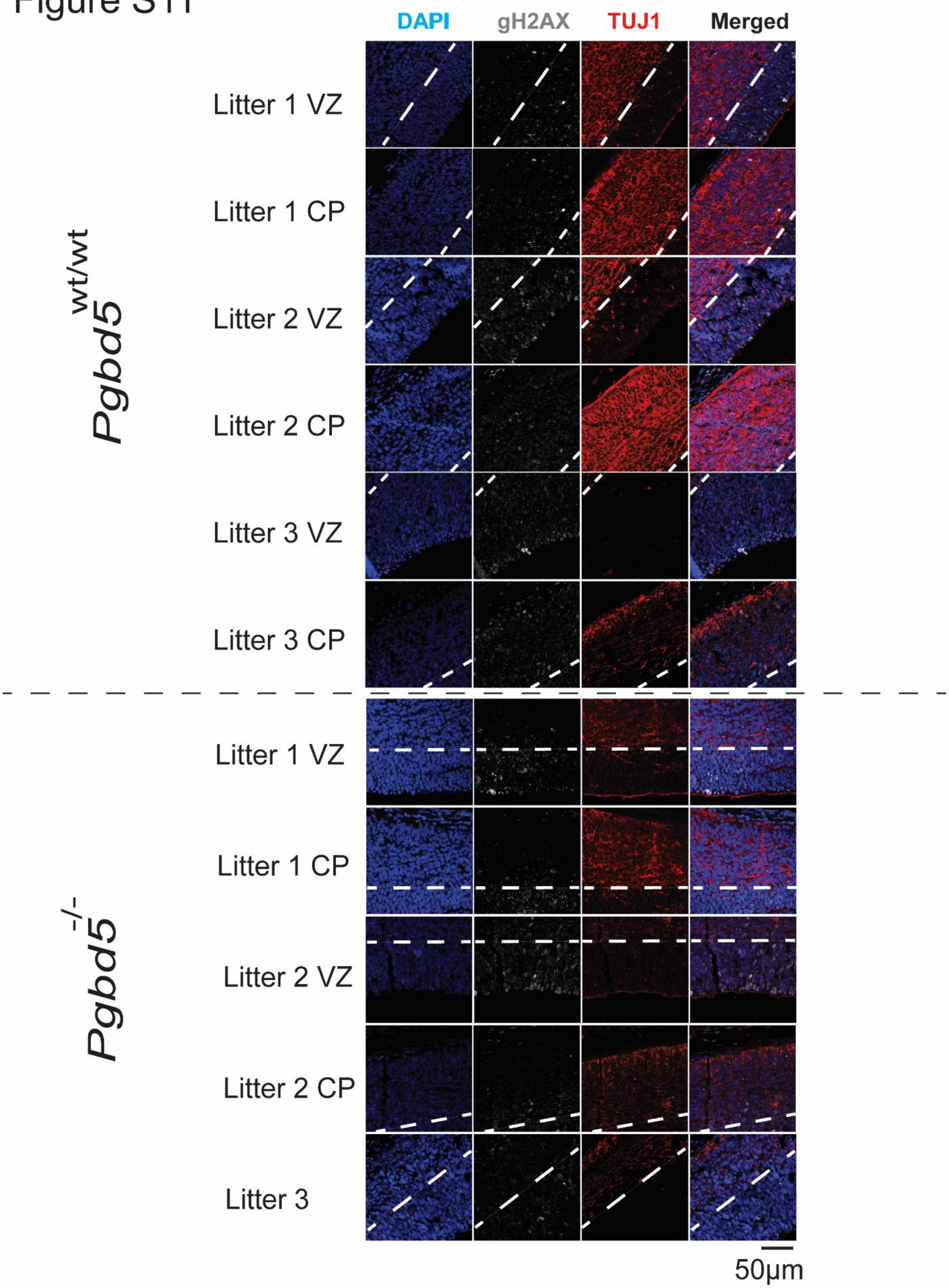
DAPI
gH2AX
TUJ1
Merged
Pgbd5 wt/wt
Litter 1 VZ
Litter 1 CP
Litter 2 VZ
Litter 2 CP
Litter 3 VZ
Litter 3 CP
Pgbd5 -/-
Litter 1 VZ
Litter 1 CP
Litter 2 VZ
Litter 2 CP
Litter 3
50µm

1

**Fig. S11. *Pgbd5*-deficient mice exhibit reduced DNA damage in region of interest 2.** Immunofluorescence photographs of brain region 2 from three different independent litters, stained for DNA using DAPI (blue), gH2AX (white), and TUJ1 (red), with specific litter mates indicated (VZ, ventricular zone; CP, cortical plate). Dashed lines indicate separation between ventricular zones and cortical plates. Scale bar = 50 mm.

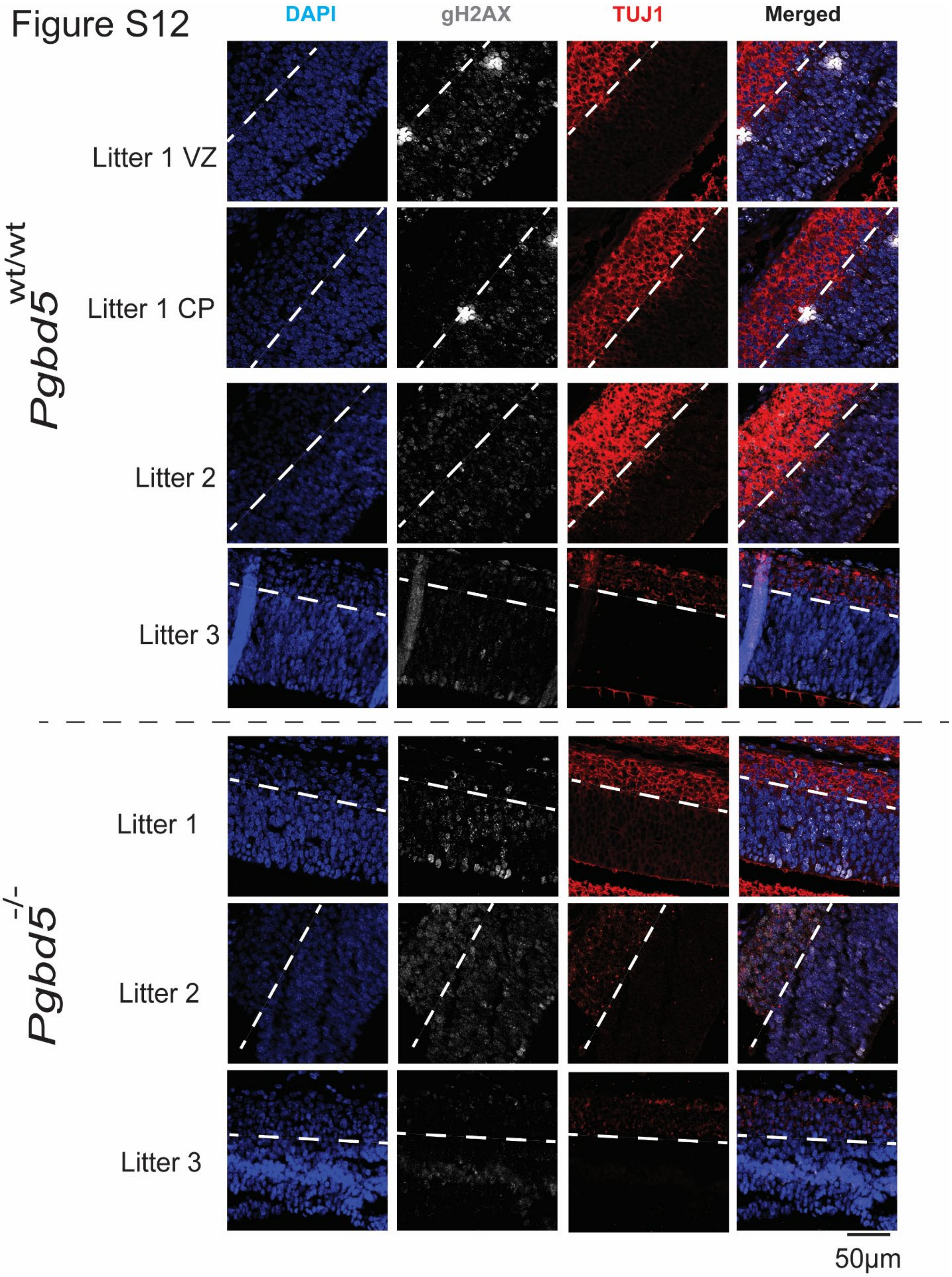

Figure S12
DAPI
gH2AX
TUJ1
Merged
Pgbd5 wt/wt
Litter 1 VZ
Litter 1 CP
Litter 2
Litter 3
Pgbd5 -/-
Litter 1
Litter 2
Litter 3
50μm

**Fig. S12.** ***Pgbd5*****-deficient mice exhibit reduced DNA damage in region of interest 3.** Immunofluorescence photographs of brain region 3 from three different independent litters, stained for DNA using DAPI (blue), gH2AX (white), and TUJ1 (red), with specific litter mates indicated (VZ, ventricular zone; CP, cortical plate). Dashed lines indicate separation between ventricular zones and cortical plates. Scale bar = 50 mm.

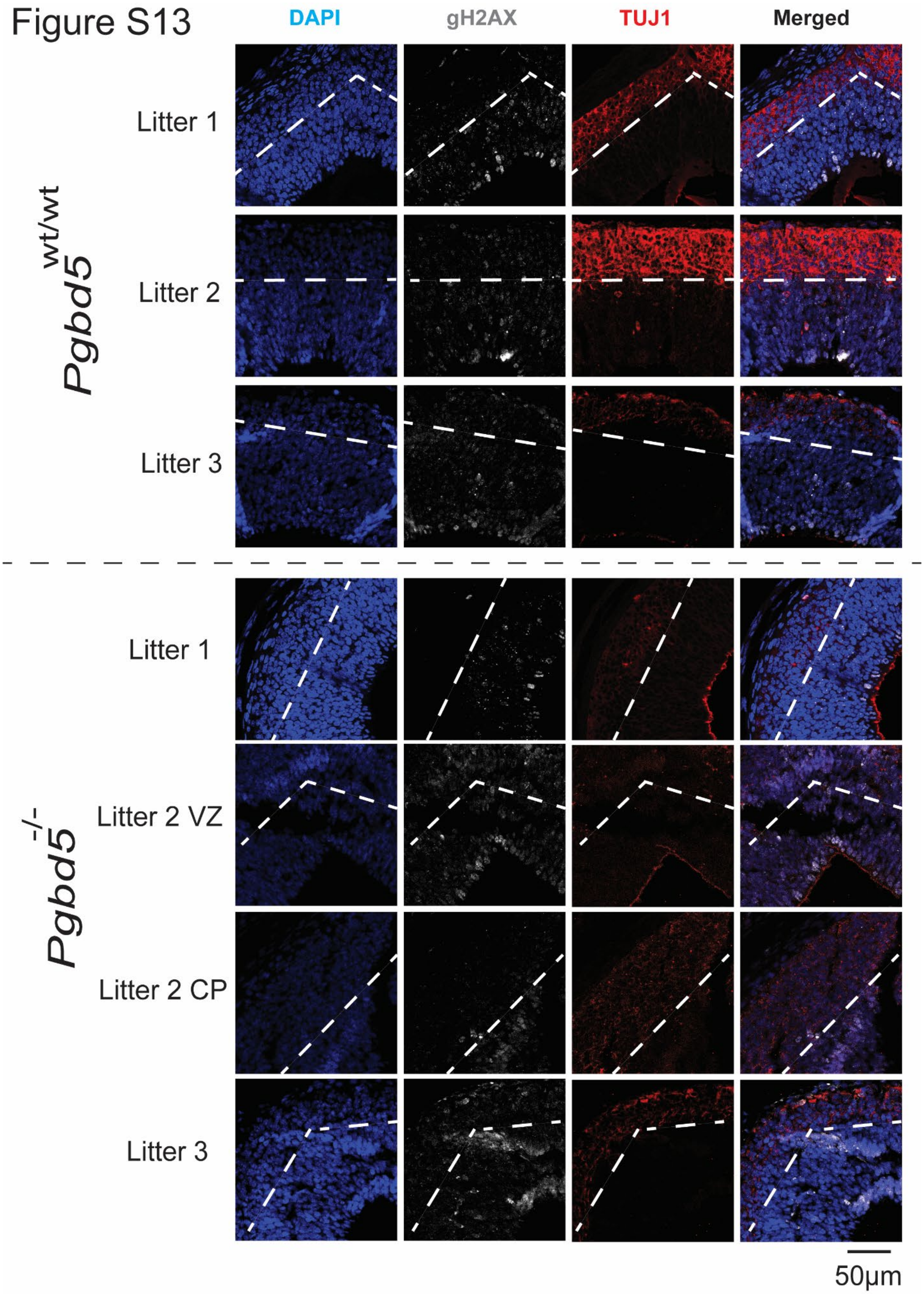
Figure S13
DAPI
gH2AX
TUJ1
Merged
Pgbd5 wt/wt
Litter 1
Litter 2
Litter 3
Pgbd5 -/-
Litter 1
Litter 2 VZ
Litter 2 CP
Litter 3
50µm

1

**Fig. S13. *Pgbd5*-deficient mice exhibit reduced DNA damage in region of interest 4.** Immunofluorescence photographs of brain region 4 from three different independent litters, stained for DNA using DAPI (blue), gH2AX (white), and TUJ1 (red), with specific litter mates indicated (VZ, ventricular zone; CP, cortical plate). Dashed lines indicate separation between ventricular zones and cortical plates. Scale bar = 50 mm.

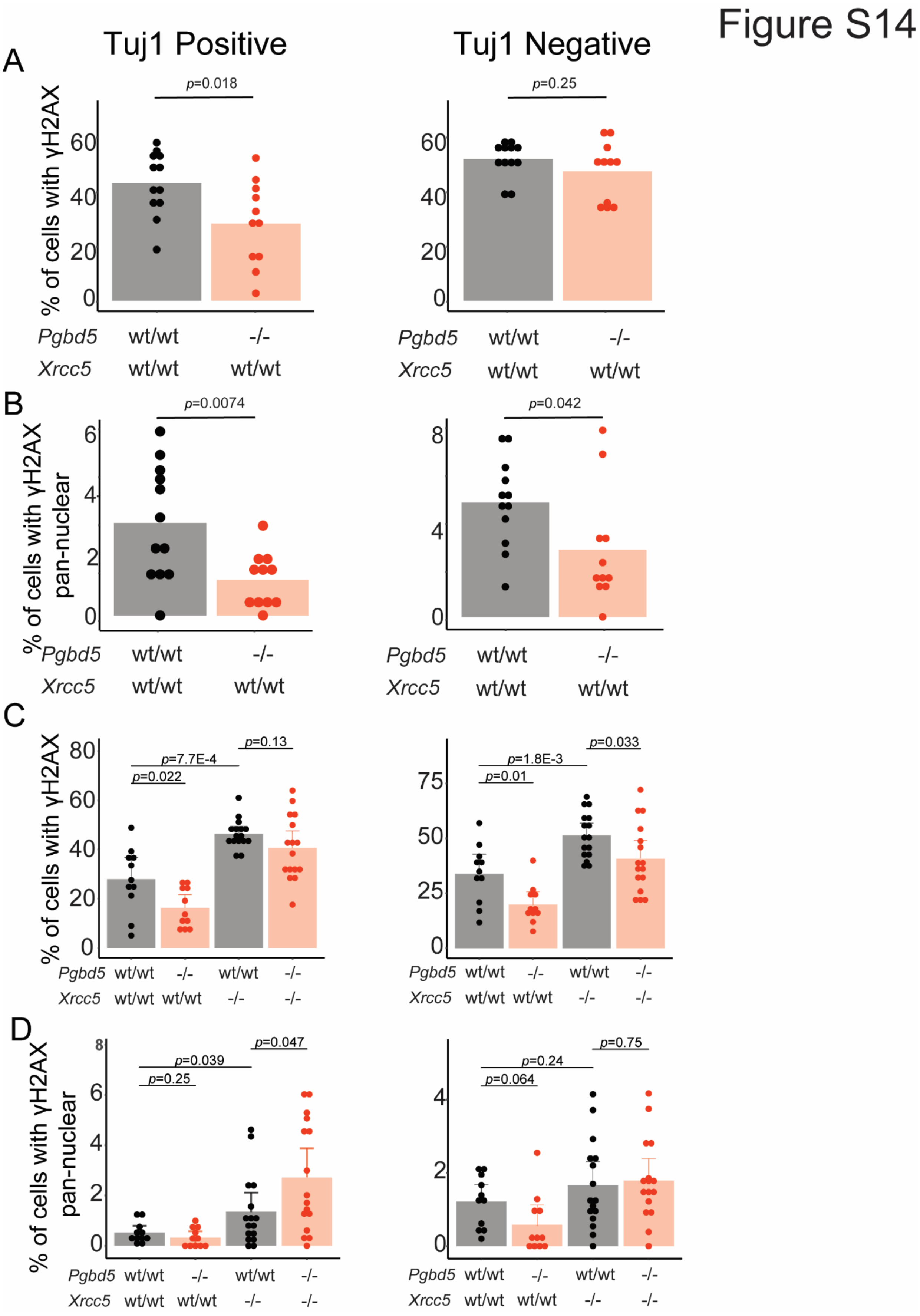

Figure S14
Tuj1 Positive
Tuj1 Negative
A
p=0.018
p=0.25
% of cells with γH2AX
Pgbd5 wt/wt -/-
Xrcc5 wt/wt wt/wt
B
p=0.0074
p=0.042
% of cells with γH2AX pan-nuclear
C
p=7.7E-4
p=0.022
p=0.13
p=1.8E-3
p=0.01
p=0.033
Pgbd5 wt/wt -/- wt/wt -/-
Xrcc5 wt/wt wt/wt -/- -/-
D
p=0.039
p=0.25
p=0.047
p=0.24
p=0.064
p=0.75


1

**Fig. S14. Double-mutant *Pgbd5*$^{-/-}$;*Xrcc5*$^{-/-}$ mice exhibit reduced DNA damage as compared to single-mutant *Pgbd5*$^{wt/wt}$;*Xrcc5*$^{-/-}$ mice, indicating the requirement of NHEJ DNA damage repair in forebrain of E14.5 embryos. A-B**, Bar plots of analysis of fraction of nuclei with punctate or pan-nuclear γH2AX in postmitotic (TUJ1 positive) and proliferative (TUJ1 negative) neurons in *Xrcc5*$^{wt/wt}$;*Pgbd5*$^{wt/wt}$ and *Xrcc5*$^{wt/wt}$;*Pgbd5*$^{-/-}$ (**A**) and *Xrcc5*$^{-/-}$;*Pgbd5*$^{wt/wt}$ and *Xrcc5*$^{-/-}$;*Pgbd5*$^{-/-}$ E14.5 mouse embryos (**B**), demonstrating a significant reduction of fraction of γH2AX-positive neurons in TUJ1-expressing cells (t-test $p = 0.018$). **C-D**, Bar plots of analysis of pan-nuclear γH2AX in postmitotic (TUJ1 positive) and proliferative (TUJ1 negative) neurons in *Xrcc5*$^{wt/wt}$;*Pgbd5*$^{wt/wt}$ and *Xrcc5*$^{wt/wt}$;*Pgbd5*$^{-/-}$ (**C**) and *Xrcc5*$^{-/-}$;*Pgbd5*$^{wt/wt}$ and *Xrcc5*$^{-/-}$;*Pgbd5*$^{-/-}$ (**D**; t-test $p = 0.0074$). Bars indicate median values.

Figure S15

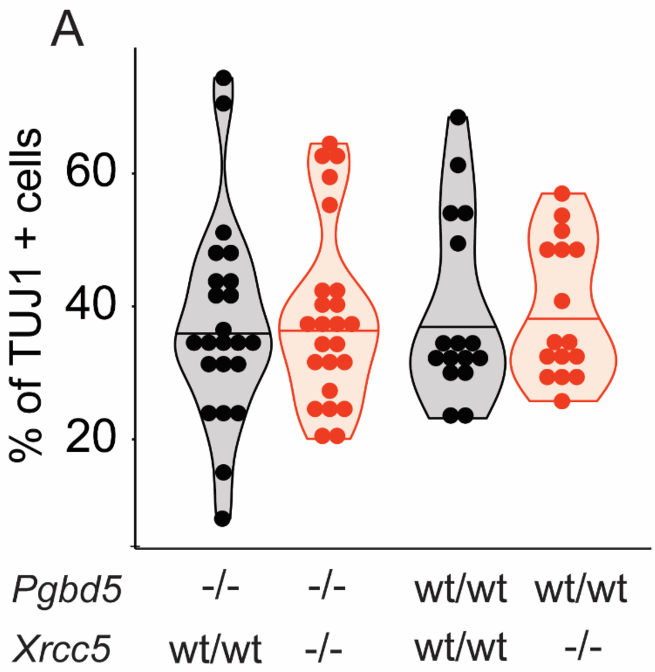


**Fig. S15.** ***Xrcc5*$^{-/-}$ and *Pgbd5*$^{-/-}$ mice do not alter postmitotic percentage of cells. A**, Violin plot for percent of TUJ1 positive cells in E14.5 developing cortex shows no significant difference across *Xrcc5* and *Pgbd5* genotypes indicating that neither of them affect the postmitotic embryonic layers of the developing cortex.

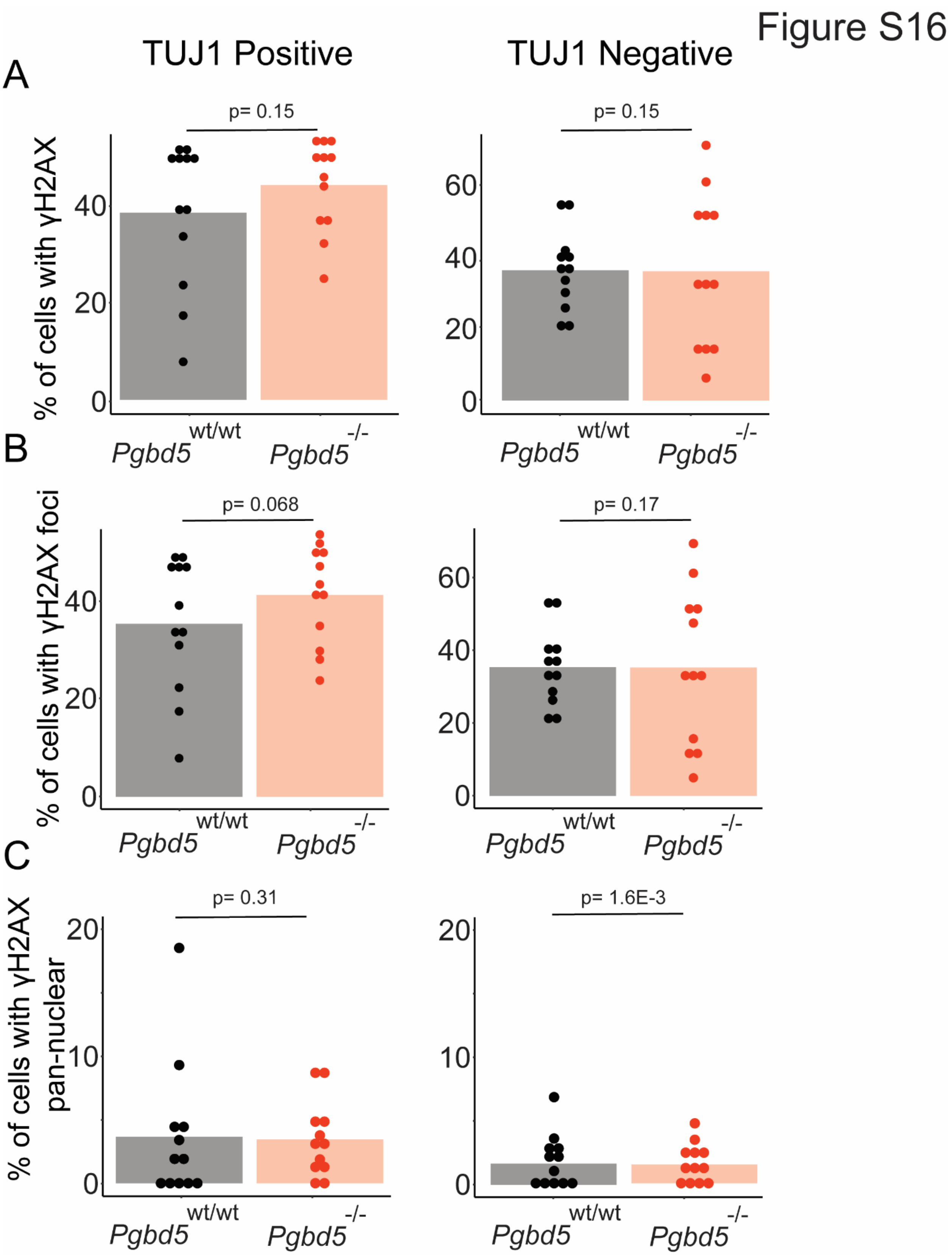
Figure S16
A
TUJ1 Positive
TUJ1 Negative
p= 0.15
p= 0.15
% of cells with γH2AX
Pgbd5 wt/wt
Pgbd5 -/-
B
p= 0.068
p= 0.17
% of cells with γH2AX foci
C
p= 0.31
p= 1.6E-3
% of cells with γH2AX pan-nuclear

**Fig. S16. *Pgbd5*-deficient E12.5 embryos lack measurable differences in brain DNA damage.** **A**, Bar plots of analysis of fraction of nuclei with punctate and pan-nuclear γH2AX in forebrain postmitotic (TUJ1 positive) and proliferative (TUJ1 negative) neurons in *Pgbd5*$^{wt/wt}$ (black) and *Pgbd5*$^{-/-}$(red). **B,** Bar plots of analysis of fraction of nuclei with punctate γH2AX in postmitotic (TUJ1 positive) and proliferative (TUJ1 negative) neurons in *Pgbd5*$^{wt/wt}$ (black) and *Pgbd5*$^{-/-}$(red). **C**, Bar plots of analysis of pan-nuclear γH2AX in postmitotic (TUJ1 positive) and proliferative (TUJ1 negative) neurons in *Pgbd5*$^{wt/wt}$ and *Pgbd5*$^{-/-}$. Bars indicate median values.

Figure S17

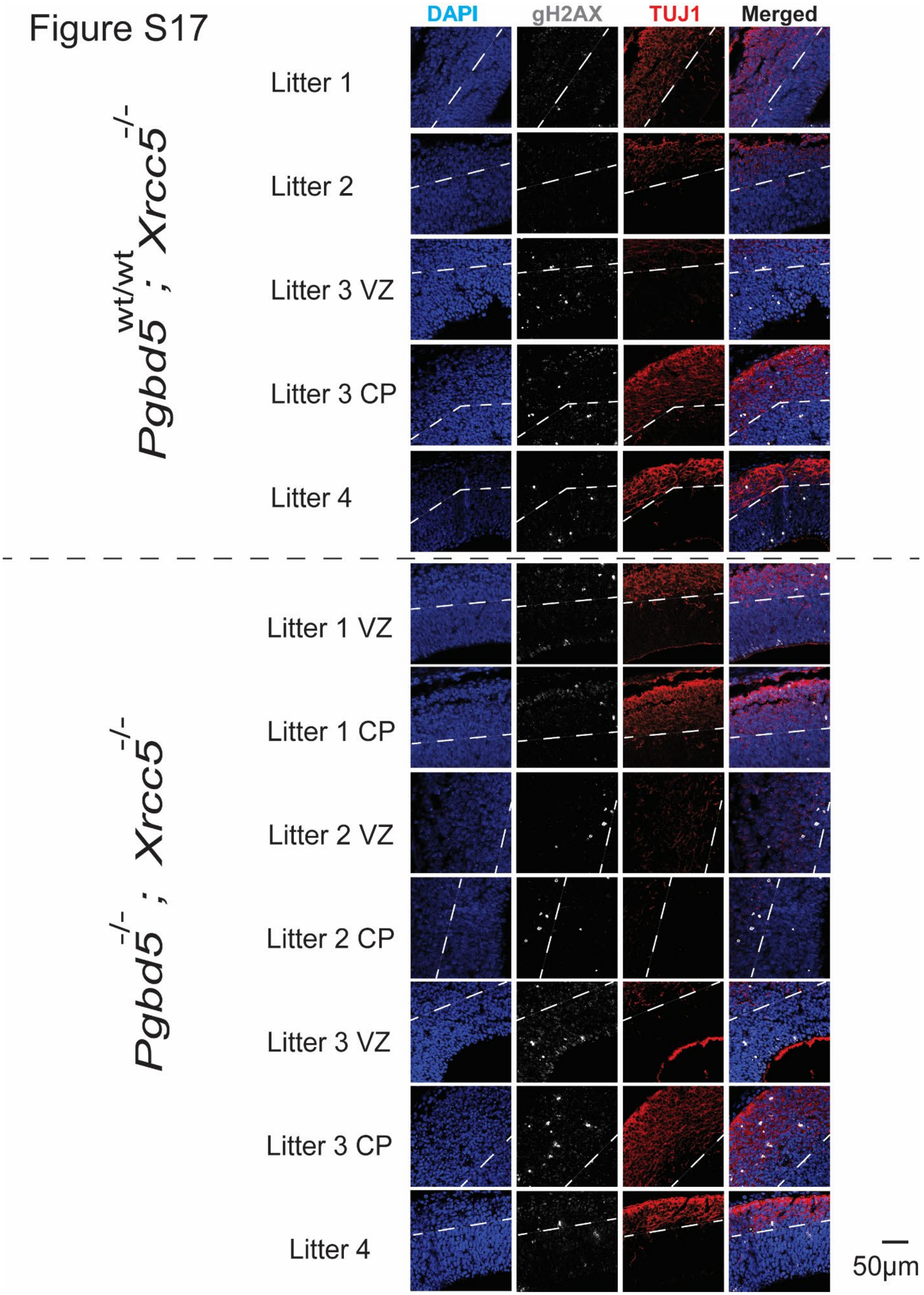
DAPI
gH2AX
TUJ1
Merged
Pgbd5 wt/wt ; Xrcc5 -/-
Litter 1
Litter 2
Litter 3 VZ
Litter 3 CP
Litter 4
Pgbd5 -/- ; Xrcc5 -/-
Litter 1 VZ
Litter 1 CP
Litter 2 VZ
Litter 2 CP
Litter 3 VZ
Litter 3 CP
Litter 4
50μm

1

**Fig. S17. *Xrcc5$^{-/-}$;Pgbd5$^{-/-}$* mice exhibit reduced DNA damage in region 1.** Representative immunofluorescence images of brain region 1 from three different independent litters, stained for DNA using DAPI (blue), gH2AX (white), and Tuj1 (red), with specific litter mates indicated (VZ, ventricular zone; CP, cortical plate). Dashed lines indicate separation between ventricular zones and cortical plates. Scale bar = 50 mm.

1 Figure S18

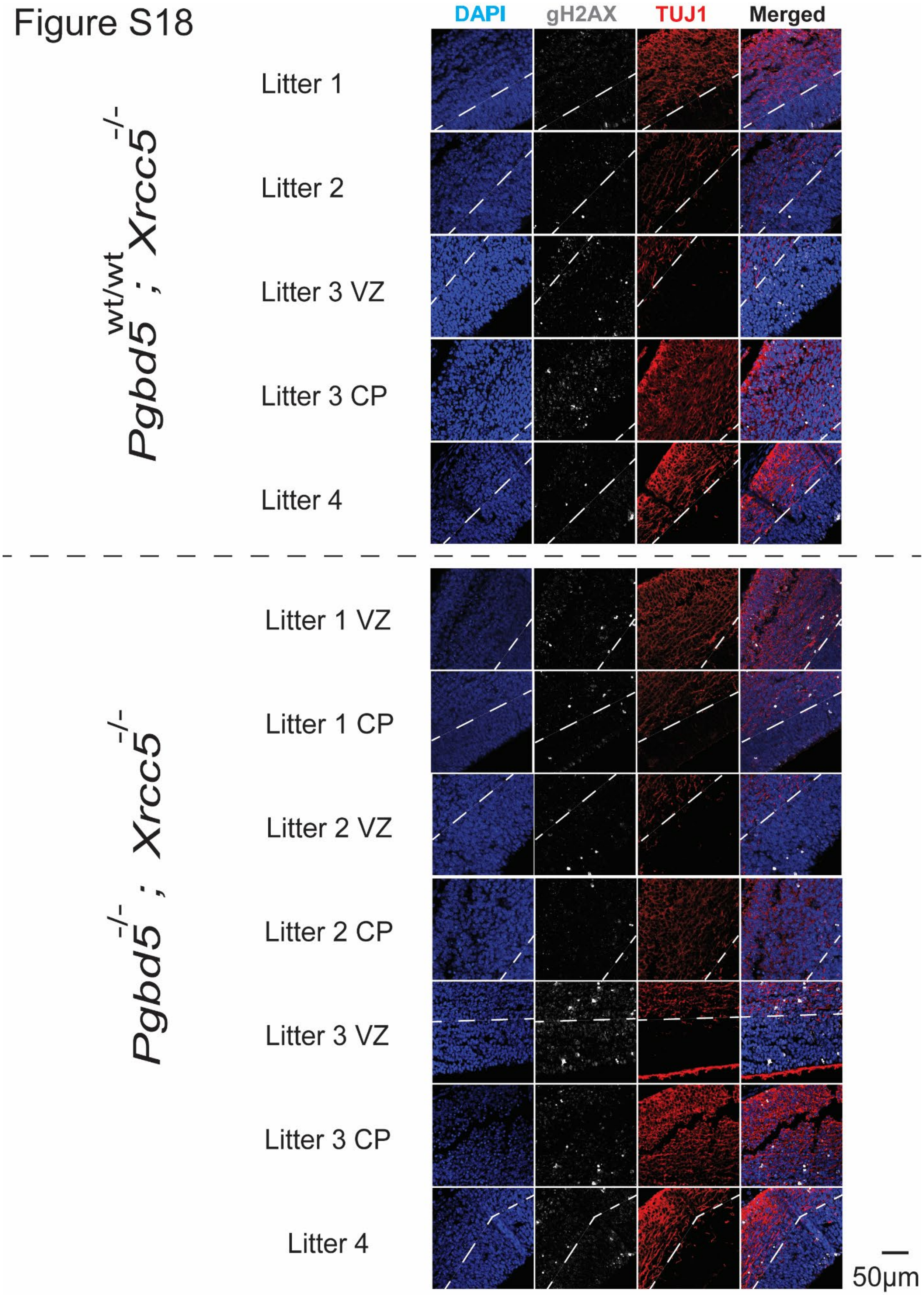
DAPI
gH2AX
TUJ1
Merged
Pgbd5 wt/wt ; Xrcc5 -/-
Litter 1
Litter 2
Litter 3 VZ
Litter 3 CP
Litter 4
Pgbd5 -/- ; Xrcc5 -/-
Litter 1 VZ
Litter 1 CP
Litter 2 VZ
Litter 2 CP
Litter 3 VZ
Litter 3 CP
Litter 4
50μm

**Fig. S18.** ***Xrcc5*$^{-/-}$*;Pgbd5*$^{-/-}$ mice exhibit reduced DNA damage in region 2.** Immunofluorescence photographs of brain region 2 from three different independent litters, stained for DNA using DAPI (blue), gH2AX (white), and Tuj1 (red), with specific litter mates indicated (VZ, ventricular zone; CP, cortical plate). Dashed lines indicate separation between ventricular zones and cortical plates. Scale bar = 50 mm.

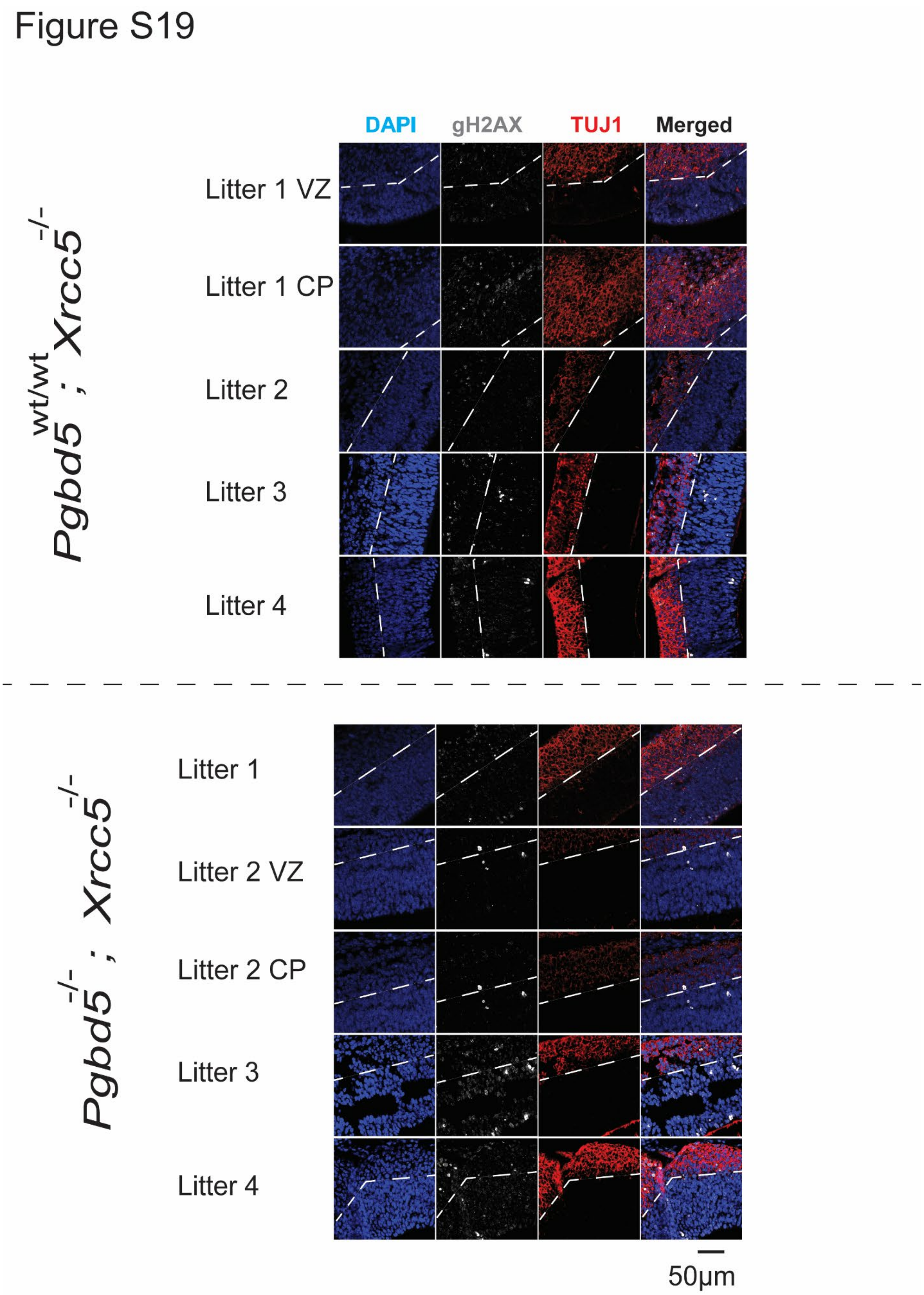
Figure S19
DAPI
gH2AX
TUJ1
Merged
Pgbd5 wt/wt ; Xrcc5 -/-
Litter 1 VZ
Litter 1 CP
Litter 2
Litter 3
Litter 4
Pgbd5 -/- ; Xrcc5 -/-
Litter 1
Litter 2 VZ
Litter 2 CP
Litter 3
Litter 4
50μm

1

**Fig. S19.** ***Xrcc5*$^{-/-}$*;Pgbd5*$^{-/-}$ mice exhibit reduced DNA damage in region 3.** Immunofluorescence photographs of brain region 3 from three different independent litters, stained for DNA using DAPI (blue), gH2AX (white), and Tuj1 (red), with specific litter mates indicated (VZ, ventricular zone; CP, cortical plate). Dashed lines indicate separation between ventricular zones and cortical plates. Scale bar = 50 mm.

### Figure S20

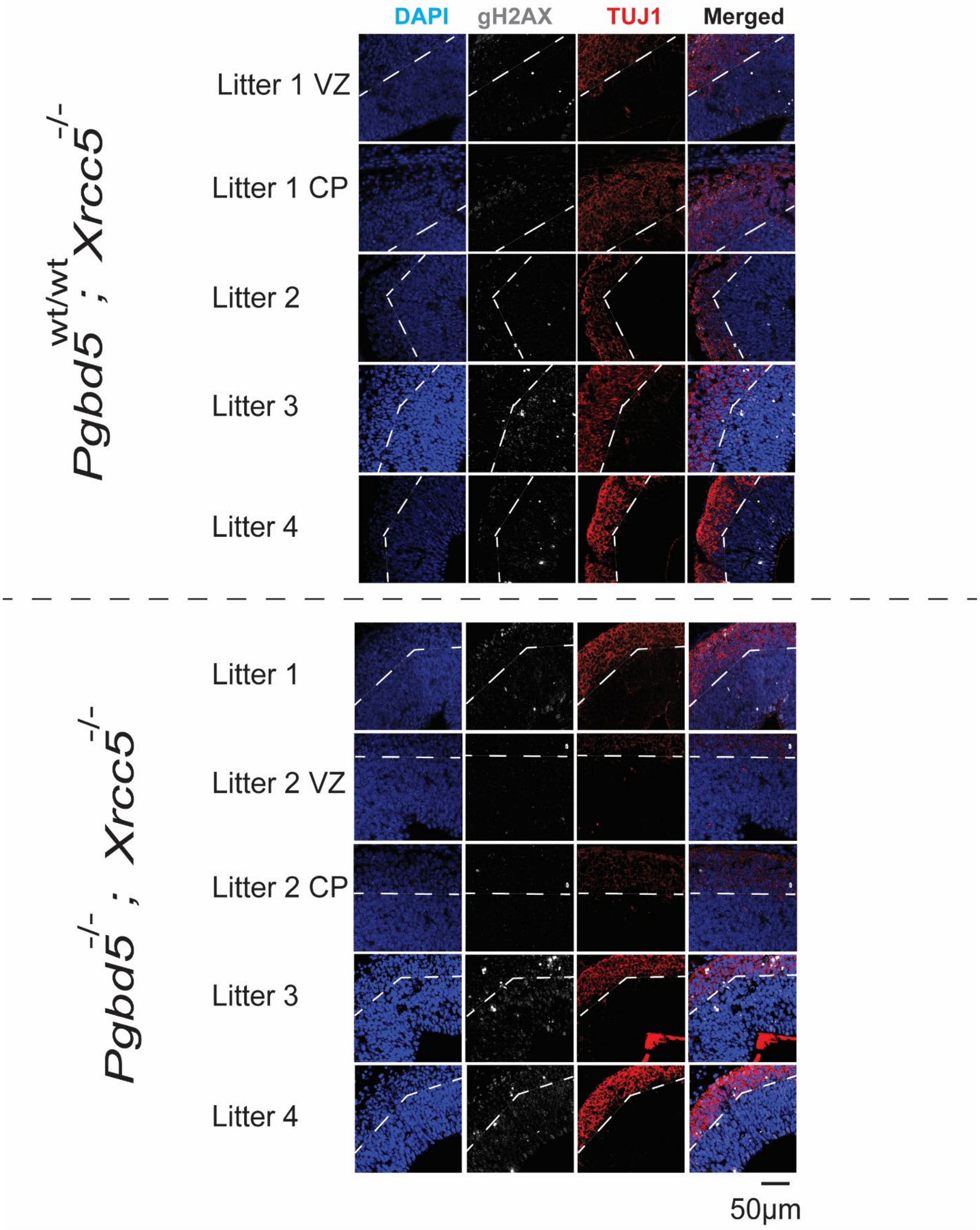
DAPI
gH2AX
TUJ1
Merged
Pgbd5wt/wt ; Xrcc5-/-
Litter 1 VZ
Litter 1 CP
Litter 2
Litter 3
Litter 4
Pgbd5-/- ; Xrcc5-/-
Litter 1
Litter 2 VZ
Litter 2 CP
Litter 3
Litter 4
50µm

**Fig. S20. *Xrcc5*$^{-/-}$*;Pgbd5*$^{-/-}$ mice exhibit reduced DNA damage in region 4.** Representative immunofluorescence images of brain region 4 from three different independent litters, stained for DNA using DAPI (blue), gH2AX (white), and TUJ1 (red), with specific litter mates indicated (VZ, ventricular zone; CP, cortical plate). Dashed lines indicate separation between ventricular zones and cortical plates. Scale bar = 50 mm.

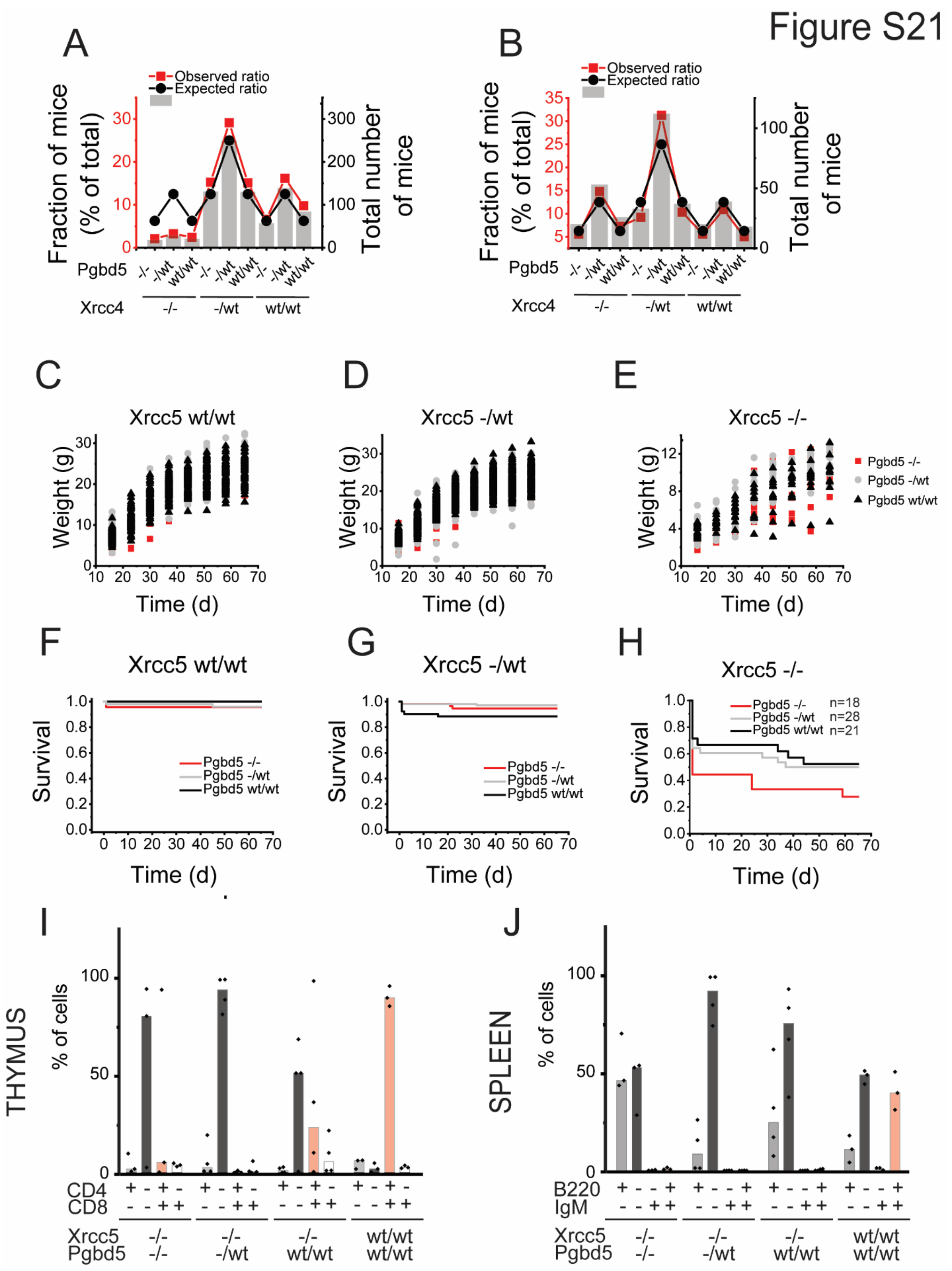
Figure S21
A
Observed ratio
Expected ratio
Fraction of mice (% of total)
Total number of mice
Pgbd5 -/- -/wt wt/wt -/- -/wt wt/wt -/- -/wt wt/wt
Xrcc4 -/- -/wt wt/wt
B
Observed ratio
Expected ratio
Fraction of mice (% of total)
Total number of mice
Pgbd5 -/- -/wt wt/wt -/- -/wt wt/wt -/- -/wt wt/wt
Xrcc4 -/- -/wt wt/wt
C
Xrcc5 wt/wt
Weight (g)
Time (d)
D
Xrcc5 -/wt
Weight (g)
Time (d)
E
Xrcc5 -/-
Weight (g)
Time (d)
Pgbd5 -/-
Pgbd5 -/wt
Pgbd5 wt/wt
F
Xrcc5 wt/wt
Survival
Pgbd5 -/-
Pgbd5 -/wt
Pgbd5 wt/wt
Time (d)
G
Xrcc5 -/wt
Survival
Pgbd5 -/-
Pgbd5 -/wt
Pgbd5 wt/wt
Time (d)
H
Xrcc5 -/-
Survival
Pgbd5 -/- n=18
Pgbd5 -/wt n=28
Pgbd5 wt/wt n=21
Time (d)
I
THYMUS
% of cells
CD4 + - + - + - + - + - + - + - + -
CD8 - - + + - - + + - - + + - - + +
Xrcc5 -/- -/- -/- wt/wt
Pgbd5 -/- -/wt wt/wt wt/wt
J
SPLEEN
% of cells
B220 + - - + + - - + + - - + + - - +
IgM - - + + - - + + - - + + - - + +
Xrcc5 -/- -/- -/- wt/wt
Pgbd5 -/- -/wt wt/wt wt/wt

**Fig. S21.** ***Pgbd5*** **deficiency does not rescue** ***Xrcc5*****-deficiency induced immunodeficiency and exacerbates** ***Xrcc5*****-deficiency induced runted phenotype.** **A-B**, Mendelian ratios of *Xrcc5*$^{wt/-}$*; Pgbd5*$^{wt/-}$ in-crosses of prenatal (**A**) and postnatal mice (**B**)**.** **C-E,** Whisker plot analysis of body weight (g, grams) of *Xrcc5*$^{wt/wt}$ (**C**), *Xrcc5*$^{wt/-}$ (**D**), *Xrcc5*$^{-/-}$ (**E**) and all the different *Pgbd5* genotype mice as a function of age (d, days). **F-H,** Kaplan-Meier survival analysis of *Xrcc5*$^{wt/wt}$ (**F**), *Xrcc5*$^{wt/-}$ (**G**), *Xrcc5*$^{-/-}$ (**H**) and all the different *Pgbd5* genotype mice. **I-J**, Fluorescence activated flow cytometry analysis of immature and mature T-cells (**I**) and B-cells (**J**), isolated from thymuses and spleens of 60-day old mice. Bars represent mean values of 3 independent biological replicates.

### Figure S22

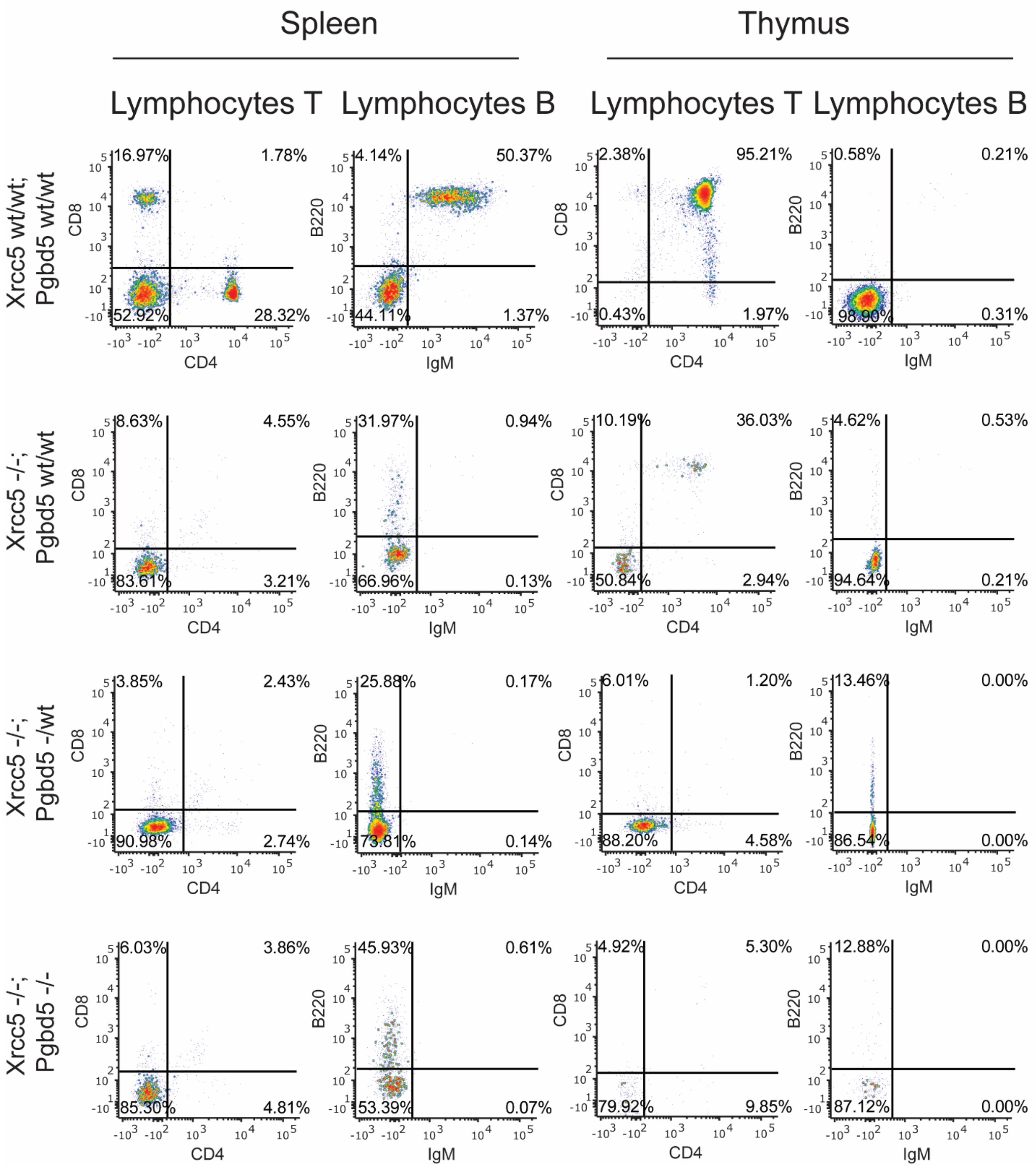
Spleen
Thymus
Lymphocytes T
Lymphocytes B
Lymphocytes T
Lymphocytes B
Xrcc5 wt/wt; Pgbd5 wt/wt
16.97% 1.78% 52.92% 28.32%
4.14% 50.37% 44.11% 1.37%
2.38% 95.21% 0.43% 1.97%
0.58% 0.21% 98.90% 0.31%
Xrcc5 -/-; Pgbd5 wt/wt
8.63% 4.55% 83.61% 3.21%
31.97% 0.94% 66.96% 0.13%
10.19% 36.03% 50.84% 2.94%
4.62% 0.53% 94.64% 0.21%
Xrcc5 -/-; Pgbd5 -/wt
3.85% 2.43% 90.98% 2.74%
25.88% 0.17% 73.81% 0.14%
6.01% 1.20% 88.20% 4.58%
13.46% 0.00% 86.54% 0.00%
Xrcc5 -/-; Pgbd5 -/-
6.03% 3.86% 85.30% 4.81%
45.93% 0.61% 53.39% 0.07%
4.92% 5.30% 79.92% 9.85%
12.88% 0.00% 87.12% 0.00%
CD8
CD4
B220
IgM

**Fig. S22.** ***Pgbd5*** **deficiency does not rescue** ***Xrcc5*****-deficiency induced immunodeficiency.** Representative fluorescence activated flow cytometry plots for spleen (left) and thymus (right) lymphocytes isolated from 60-day old mice, stained with specific T- (CD4, CD8) and B-cell (B220, IgM) markers, as indicated.

### Figure S23

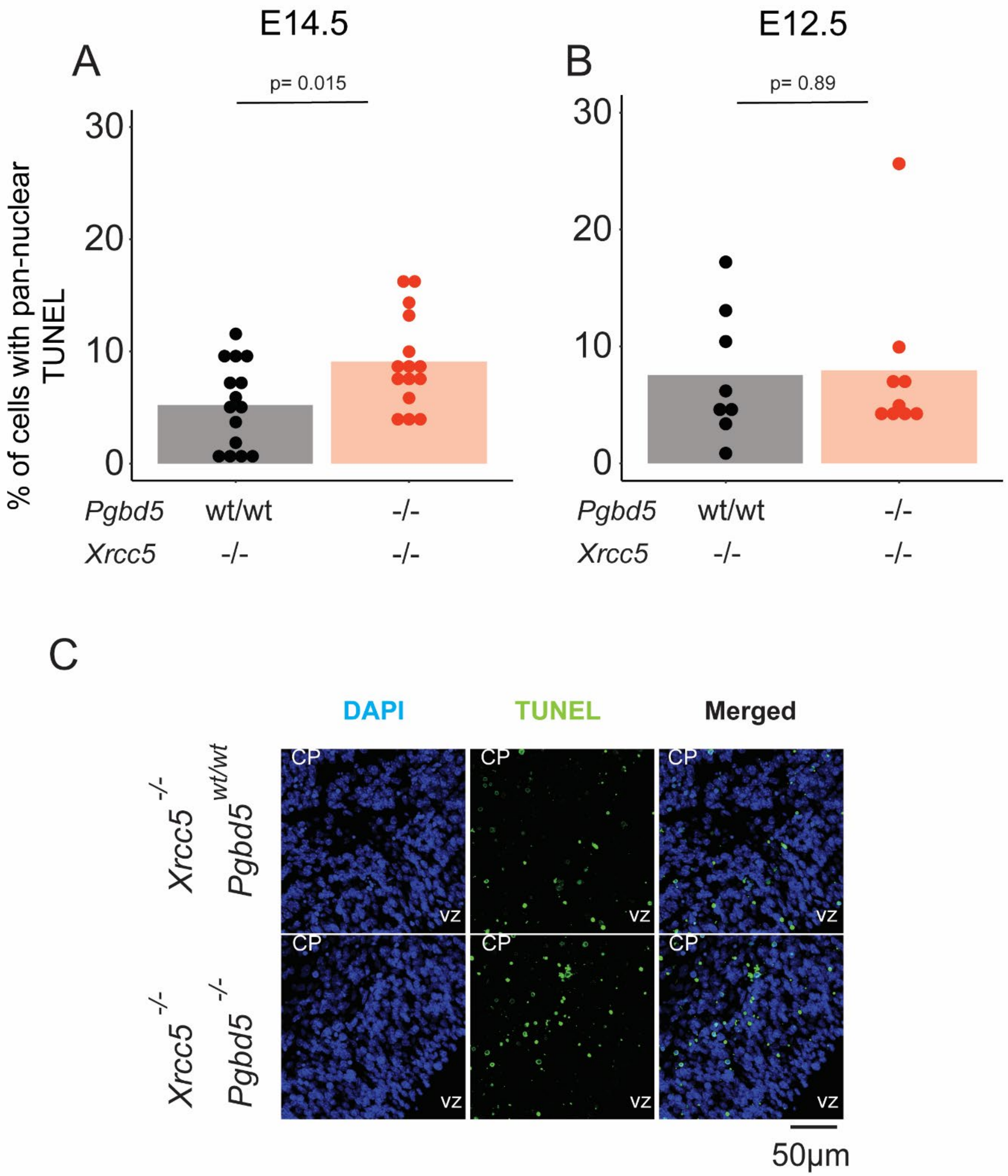
E14.5
A
p= 0.015
30
20
10
0
% of cells with pan-nuclear TUNEL
Pgbd5 wt/wt -/-
Xrcc5 -/- -/-
E12.5
B
p= 0.89
30
20
10
0
Pgbd5 wt/wt -/-
Xrcc5 -/- -/-
C
DAPI
TUNEL
Merged
Xrcc5 -/- Pgbd5 wt/wt
Xrcc5 -/- Pgbd5 -/-
CP
VZ
50μm

**Fig. S23. *Pgbd5* knock-out worsens the neuronal death exhibited in *Xrcc5* knock-out mice.** **A-B**, Neuronal apoptosis in forebrains of E14.5 mice measured using terminal deoxynucleotidyl transferase biotin-dUTP nick end labeling (TUNEL) shows a significant increase in E14.5 $Xrcc5^{-/-};Pgbd5^{-/-}$ embryos ($p$=1.5E-2) (**A**) but not in E12.5 ($p$=0.89) (**B**). **C**, Representative images of E14.5 $Xrcc5^{-/-};Pgbd5^{-/-}$ TUNEL.

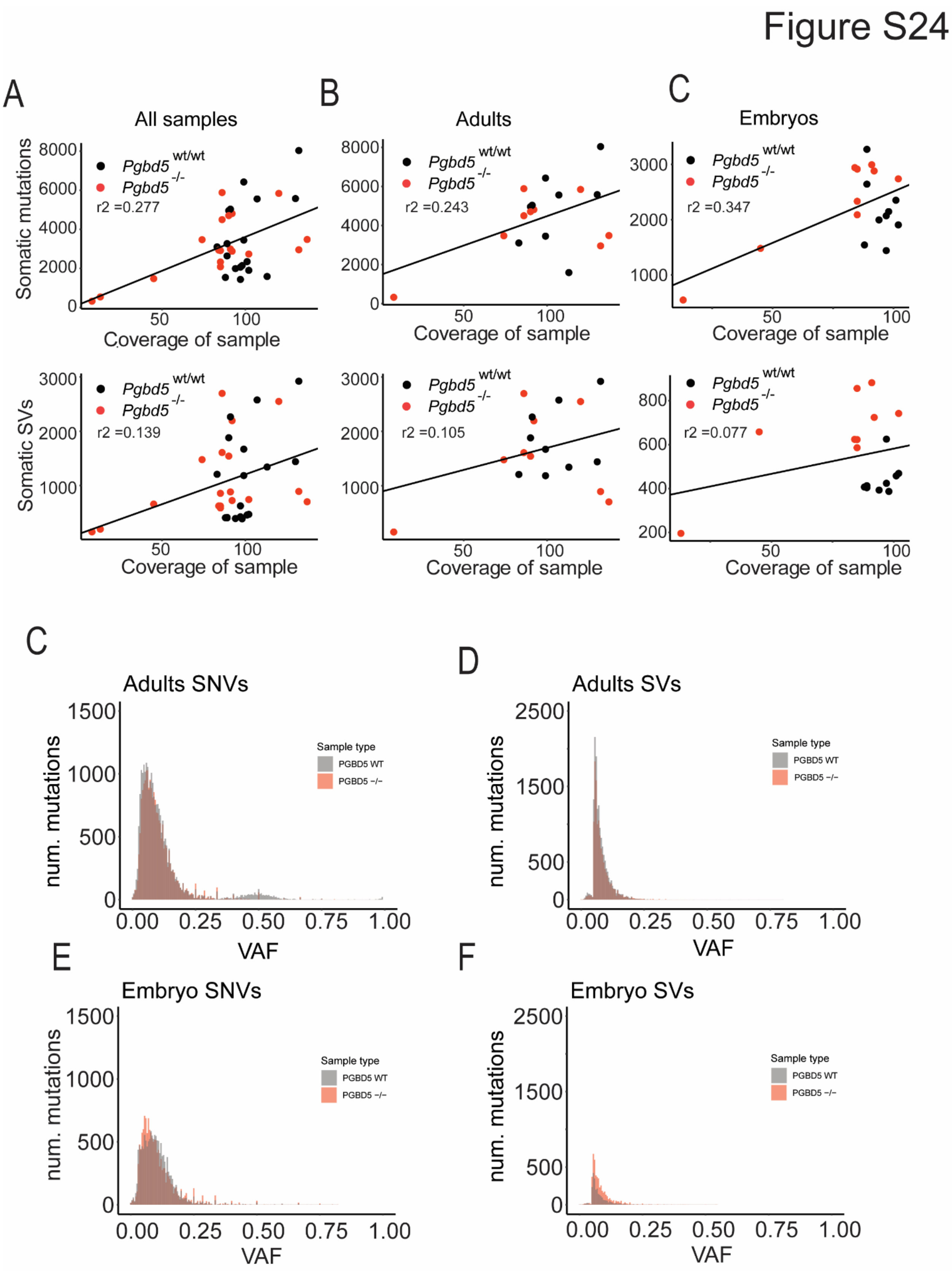
Figure S24
A
All samples
B
Adults
C
Embryos
Somatic mutations
Pgbd5 wt/wt
Pgbd5 -/-
r2 =0.277
r2 =0.243
r2 =0.347
Coverage of sample
Somatic SVs
r2 =0.139
r2 =0.105
r2 =0.077
C
Adults SNVs
D
Adults SVs
num. mutations
VAF
Sample type
PGBD5 WT
PGBD5 −/−
E
Embryo SNVs
F
Embryo SVs

**Fig. S24. WGS reliably detects infrequent variants. A-B**, Sample sequencing coverage analysis shows no significant correlation between detection of somatic variants and mean sequencing coverage for somatic SNVs (**A**) and structural variants (**B**). **C-D**, Variant allele frequency (VAF) histogram for adult single nucleotide variants (SNV) (**C**) and structural variants (SV) (**D**) in *Pgbd5*$^{wt/wt}$ and *Pgbd5*$^{-/-}$. **E-F**, Variant allele frequency histogram for embryo SNV (**E**) and SV (**F**) in *Pgbd5*$^{wt/wt}$ (grey) and *Pgbd5*$^{-/-}$(red) show that the mutations found are smaller than 0.25 indicating that they are somatic.

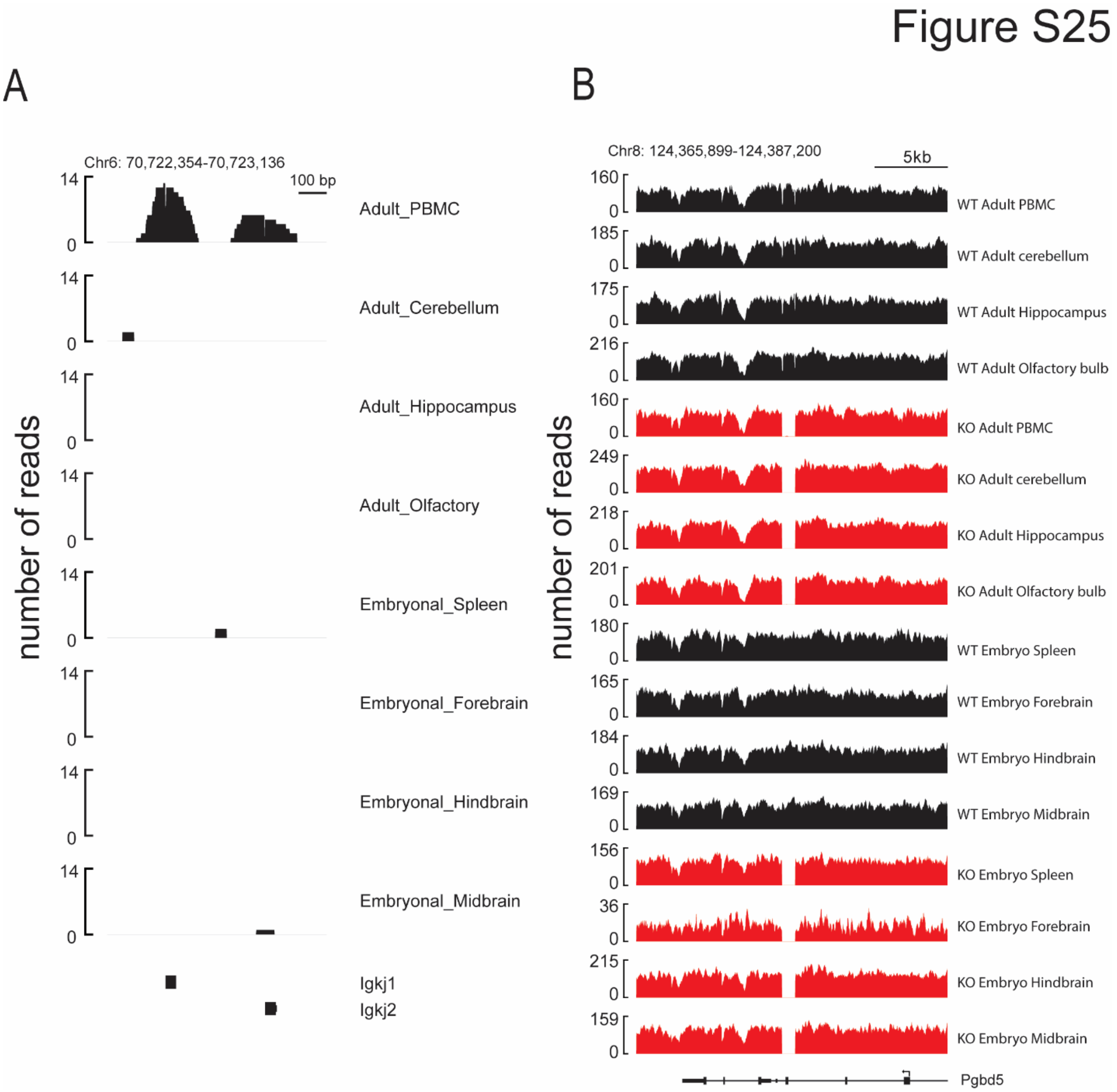


**Fig. S25. Reliable detection of immunoglobulin locus rearrangements.** **A**, Number of split reads of Igk1 and Igk2 locus in *Pgbd5*$^{wt/wt}$ adult mice show recombination only on peripheral blood mononuclear cells (PBMC) and embryo mice shows no recombination in any tissue. **B**, Split reads on Pgbd5 locus showing recombination of exon 4 across all analyzed tissues in *Pgbd5*$^{wt/wt}$ adult mice and embryo mice**.**

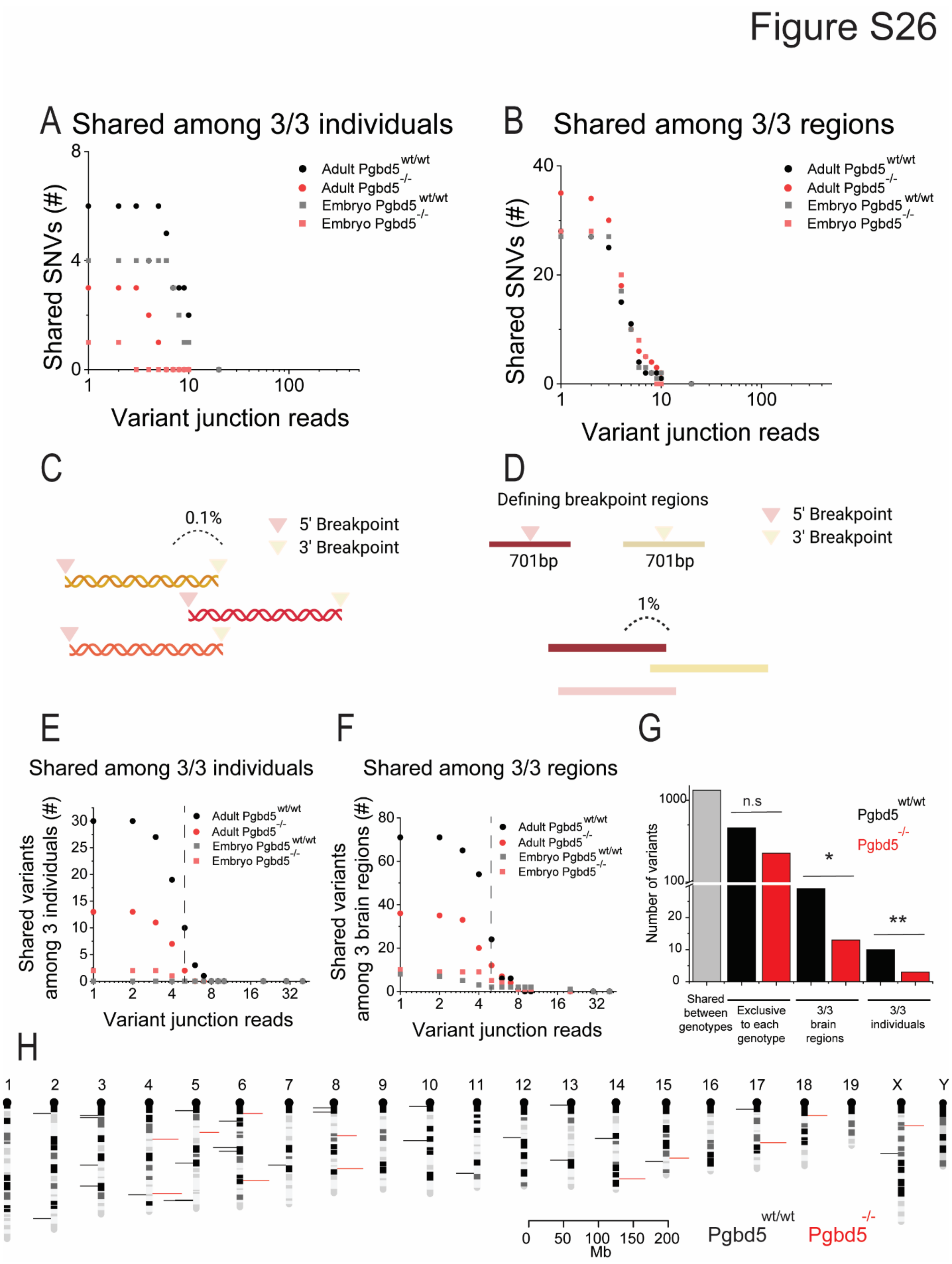
Figure S26
A Shared among 3/3 individuals
Shared SNVs (#)
Variant junction reads
Adult Pgbd5 wt/wt
Adult Pgbd5 -/-
Embryo Pgbd5 wt/wt
Embryo Pgbd5 -/-
B Shared among 3/3 regions
Shared SNVs (#)
Variant junction reads
C
0.1%
5' Breakpoint
3' Breakpoint
D
Defining breakpoint regions
701bp
701bp
1%
5' Breakpoint
3' Breakpoint
E
Shared among 3/3 individuals
Shared variants among 3 individuals (#)
Variant junction reads
F
Shared among 3/3 regions
Shared variants among 3 brain regions (#)
Variant junction reads
G
Number of variants
n.s
*
**
Pgbd5 wt/wt
Pgbd5 -/-
Shared between genotypes
Exclusive to each genotype
3/3 brain regions
3/3 individuals
H
Mb
Pgbd5 wt/wt
Pgbd5 -/-

**Fig. S26. Pgbd5 wildtype brains show recurrent structural variants among different individuals and brain regions. A-B,** Dot plots showing number of small nucleotide variants (SNV) at different variant junction reads thresholds shared across three individuals (**A**) and three brain regions (**B**) in adult and embryo *Pgbd5*$^{wt/wt}$ (black and grey respectively) and *Pgbd5*$^{-/-}$(red and light-red respectively). No significant difference was observed between *Pgbd5*$^{wt/wt}$ and *Pgbd5*$^{-/-}$ across the different number of supporting variant junction reads. **C**, Schematics showing the method for calculating overlap among structural variants looking at the actual genomic range (range analysis). In this case we only considered the variants that had at least an overlap of 0.1% among three individuals or brain regions. **D**, Schematics showing the method for calculating overlap among structural variants looking at the breakpoints. In this case, we defined a region of 701bp with each breakpoint on the center (breakpoint analysis). We then considered the variants that had at least an overlap of 1% among three individuals or brain regions. **E-F**, Dot plots showing numbers of recurrent somatic structural variants as a function of numbers of supporting sequencing reads shared among three individual mice (**E**) and three brain regions (**F**) in adult and embryonal brain tissues for *Pgbd5*$^{wt/wt}$ (black and grey, respectively) and *Pgbd5*$^{-/-}$ litter mate mice (red and pink, respectively). The overlap among structural variants was calculated using the range analysis: only the variants with 0.1% or more overlap were considered. Dashed line denotes allele fraction of 5 supporting reads, which minimizes most stochastic and artifactual somatic variants, as evident from their elimination in *Pgbd5*$^{-/-}$ tissues ($\chi^2$ test $p$ =1.6E-17 and 1.3E-9 for recurrence among different individuals and brain regions, respectively). **G**, Bar plot summarizing the results from Fig. 5**A** and **B** using the threshold of 5 reads or more. Significant differences between the number of recurrent rearrangements between *Pgbd5*$^{wt/wt}$ and *Pgbd5*$^{-/-}$ in three individuals ($\chi^2$ $p$ =1.6E-17) and three brain regions ($\chi^2$ test $p$ =1.3E-9) are observed. No significant difference is observed in non-recurrent somatic rearrangements. **H**, Mouse chromosomal ideograms showing the locations of recurrent somatic DNA rearrangements in three brain regions observed in *Pgbd5*$^{wt/wt}$ (black) and *Pgbd5*$^{-/-}$ (red) brains binned in 1 million base intervals.

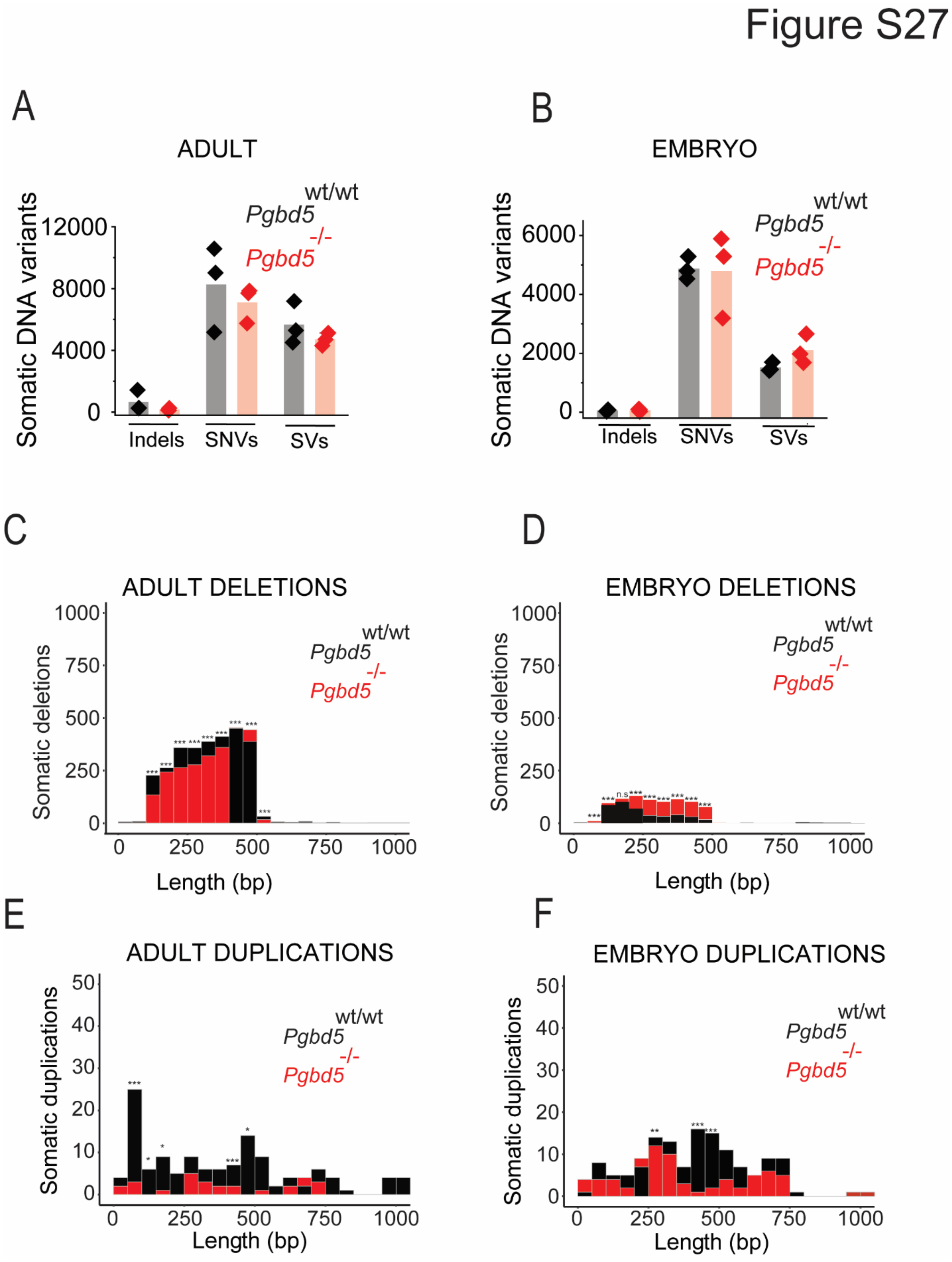
Figure S27
A
ADULT
Somatic DNA variants
12000
8000
4000
0
wt/wt
Pgbd5
-/-
Pgbd5
Indels
SNVs
SVs
B
EMBRYO
6000
4000
2000
C
ADULT DELETIONS
Somatic deletions
1000
750
500
250
Length (bp)
D
EMBRYO DELETIONS
n.s
E
ADULT DUPLICATIONS
Somatic duplications
50
40
30
20
10
F
EMBRYO DUPLICATIONS

**Fig. S27. Pgbd5 wildtype exclusive mutations are recurrent across different brain regions and individuals.** **A**, Distribution of variants in *Pgbd5*$^{wt/wt}$ (black) and *Pgbd5*$^{-/-}$ (red) adult mice show no significant difference. **B**, Distribution of variants in *Pgbd5*$^{wt/wt}$ (black) and *Pgbd5*$^{-/-}$ (red) in embryo mice show no significant difference**. C-D**, Histogram of medium-small size deletions (100-500bp) in adult and embryo *Pgbd5*$^{wt/wt}$ (grey) and *Pgbd5*$^{-/-}$(red) mice, Pearson's $\chi^2$ test shows a significant *p*-value (1.082E-02) for 100-500bp deletions (**C**) and *Pgbd5*$^{wt/wt}$ (grey) and *Pgbd5*$^{-/-}$ (red) embryos, Pearson's $\chi^2$ tests show a significant *p*-value (3.392E-03) for 100-500bp duplications (**D**). **E-F,** Histogram of medium-small size duplications (100-500bp) in adult and embryo *Pgbd5*$^{wt/wt}$ (grey) and *Pgbd5*$^{-/-}$(red) mice, Pearson's $\chi^2$ test shows a tendency *p*-value (0.35) for 250-500bp duplications in adult mice (**E**) and *Pgbd5*$^{wt/wt}$ (grey) and *Pgbd5*$^{-/-}$(red) embryos, Pearson's $\chi^2$ test shows a trend *p*-value (0.12) for 150-500bp duplications in embryos (**F**). SNVs= small nucleotide variants, SVs= structural variants.

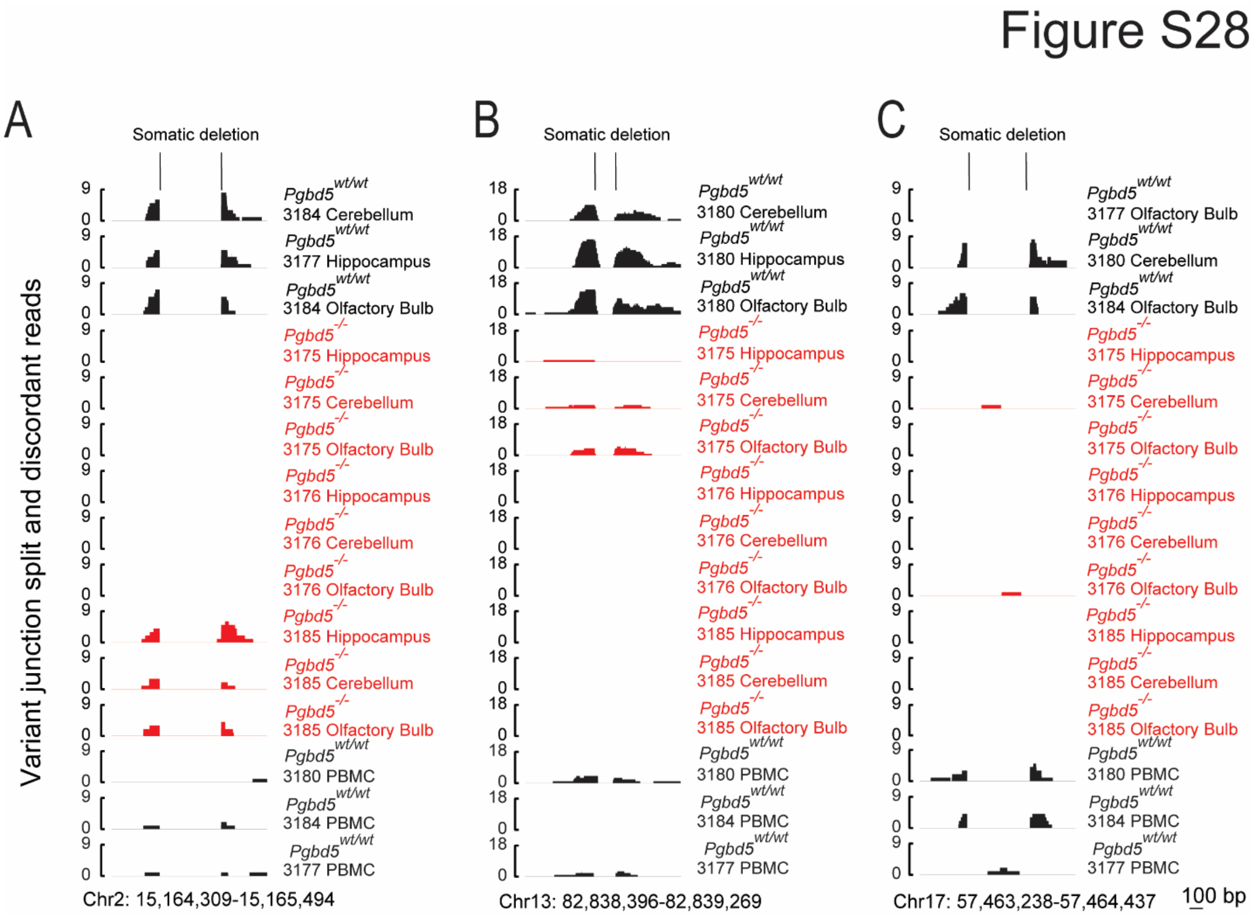


**Fig. S28. Three recurrent mutations in Pgbd5 wildtype mice passed the manual inspection as non-artifacts. A-C**¸ Histograms of variant junction split and discordant reads used by Delly2 to determine structural variants. Validated loci chr2:15164309-15165494 (**A**), chr13:82838396-82839269 **(B),** chr17:57463238-57464437 **(C).** In black, three recurrent Pgbd5wt/wt individuals and/or brain regions as well as the matching peripheral blood mononuclear cells (PBMC) samples. In red, histograms of all Pgbd5-/- mouse brain regions.

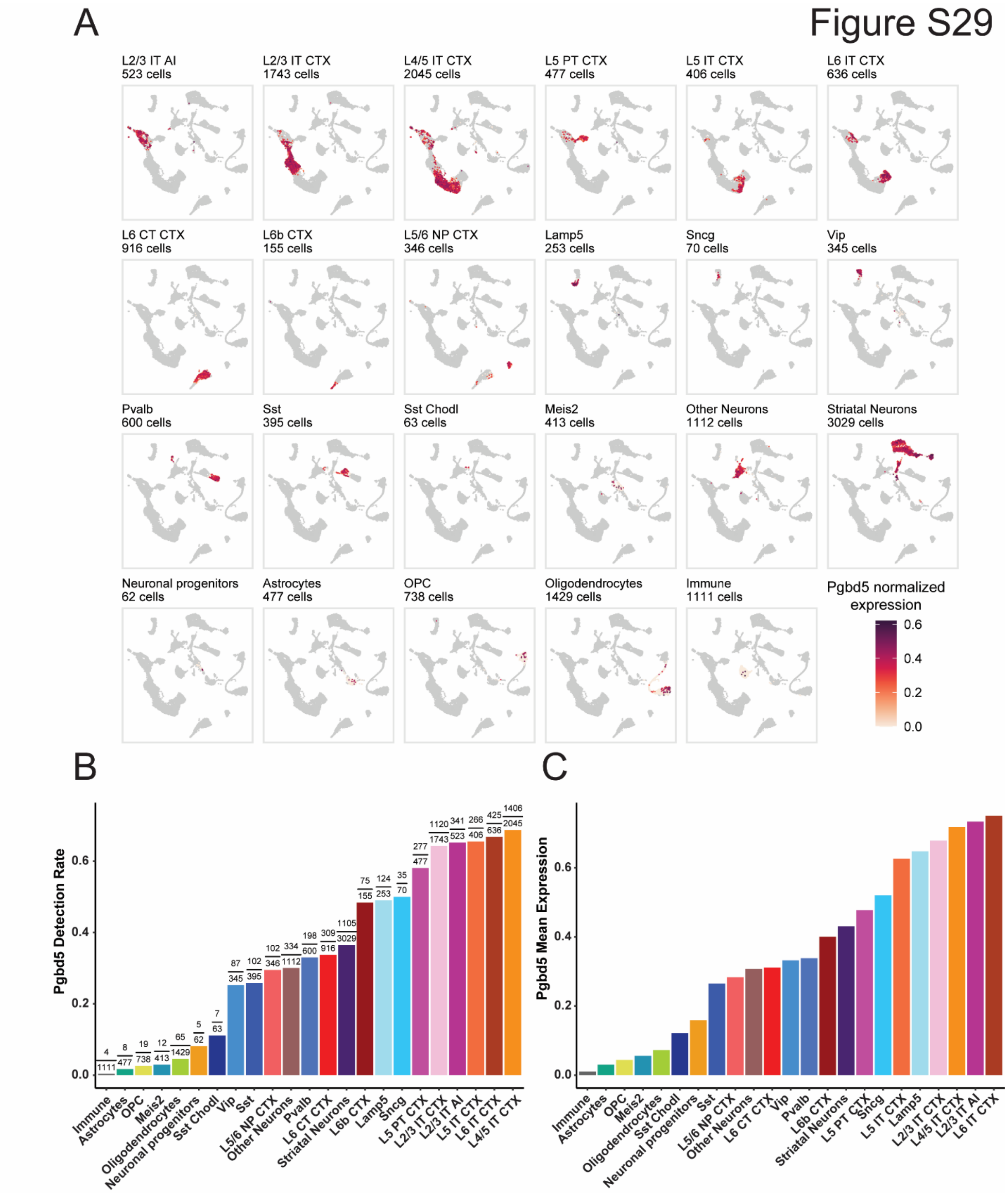

A
Figure S29
L2/3 IT AI
523 cells
L2/3 IT CTX
1743 cells
L4/5 IT CTX
2045 cells
L5 PT CTX
477 cells
L5 IT CTX
406 cells
L6 IT CTX
636 cells
L6 CT CTX
916 cells
L6b CTX
155 cells
L5/6 NP CTX
346 cells
Lamp5
253 cells
Sncg
70 cells
Vip
345 cells
Pvalb
600 cells
Sst
395 cells
Sst Chodl
63 cells
Meis2
413 cells
Other Neurons
1112 cells
Striatal Neurons
3029 cells
Neuronal progenitors
62 cells
Astrocytes
477 cells
OPC
738 cells
Oligodendrocytes
1429 cells
Immune
1111 cells
Pgbd5 normalized expression
0.6
0.4
0.2
0.0
B
Pgbd5 Detection Rate
0.0
0.2
0.4
0.6
Immune
Astrocytes
OPC
Meis2
Oligodendrocytes
Neuronal progenitors
Sst Chodl
Vip
Sst
L5/6 NP CTX
Other Neurons
Pvalb
L6 CT CTX
Striatal Neurons
L6b CTX
Lamp5
Sncg
L5 PT CTX
L2/3 IT CTX
L2/3 IT AI
L5 IT CTX
L6 IT CTX
L4/5 IT CTX
C
Pgbd5 Mean Expression
0.0
0.2
0.4
0.6
Immune
Astrocytes
OPC
Meis2
Oligodendrocytes
Sst Chodl
Neuronal progenitors
Sst
L5/6 NP CTX
Other Neurons
L6 CT CTX
Vip
Pvalb
L6b CTX
Striatal Neurons
L5 PT CTX
Sncg
L5 IT CTX
Lamp5
L2/3 IT CTX
L4/5 IT CTX
L2/3 IT AI
L6 IT CTX

**Fig. S29. Pgbd5 expression does not affect neuronal population proportions. A,** Uniform Manifold approximation and projection (UMAP) plot of single nuclei RNA seq from motor cortices of *Pgbd5*$^{wt/wt}$ littermates, samples (N = 18,107 WT) highlighting *Pgbd5* normalized expression in cells from each cluster containing >200 cells. Each cluster and their corresponding Pgbd5 expression in each neuronal identity are colored in a gradient. **B-C,** Box plots of *Pgbd5* detection rate and normalized mean expression in all the neuronal identity clusters. In (**B**) Pgbd5 detection rate is shown as a fraction of expressing cells over the total. In (**C**) Pgbd5 mean expression is colored by neuronal identity.

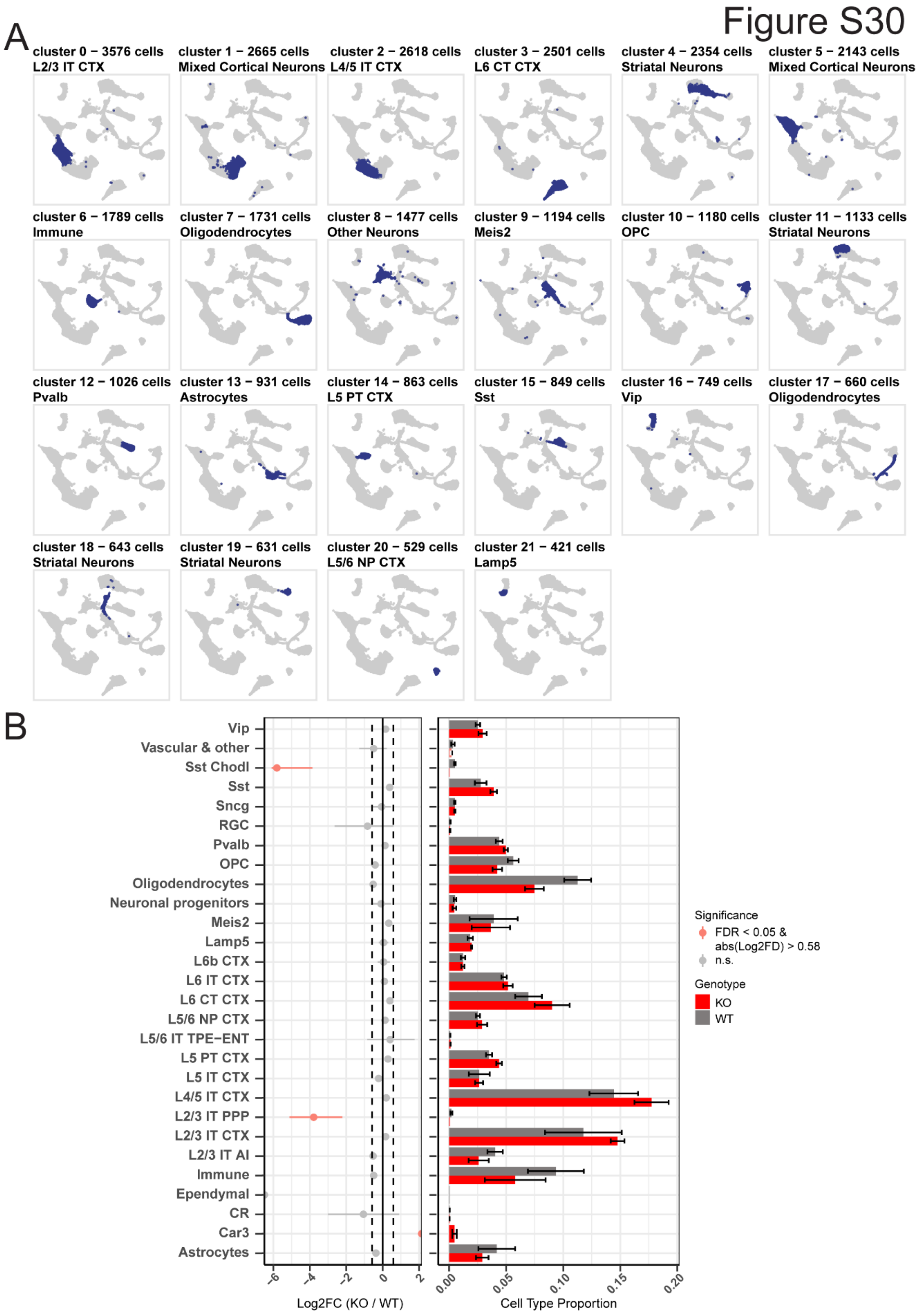
Figure S30
A
cluster 0 − 3576 cells
L2/3 IT CTX
cluster 1 − 2665 cells
Mixed Cortical Neurons
cluster 2 − 2618 cells
L4/5 IT CTX
cluster 3 − 2501 cells
L6 CT CTX
cluster 4 − 2354 cells
Striatal Neurons
cluster 5 − 2143 cells
Mixed Cortical Neurons
cluster 6 − 1789 cells
Immune
cluster 7 − 1731 cells
Oligodendrocytes
cluster 8 − 1477 cells
Other Neurons
cluster 9 − 1194 cells
Meis2
cluster 10 − 1180 cells
OPC
cluster 11 − 1133 cells
Striatal Neurons
cluster 12 − 1026 cells
Pvalb
cluster 13 − 931 cells
Astrocytes
cluster 14 − 863 cells
L5 PT CTX
cluster 15 − 849 cells
Sst
cluster 16 − 749 cells
Vip
cluster 17 − 660 cells
Oligodendrocytes
cluster 18 − 643 cells
Striatal Neurons
cluster 19 − 631 cells
Striatal Neurons
cluster 20 − 529 cells
L5/6 NP CTX
cluster 21 − 421 cells
Lamp5
B
Vip
Vascular & other
Sst Chodl
Sst
Sncg
RGC
Pvalb
OPC
Oligodendrocytes
Neuronal progenitors
Meis2
Lamp5
L6b CTX
L6 IT CTX
L6 CT CTX
L5/6 NP CTX
L5/6 IT TPE−ENT
L5 PT CTX
L5 IT CTX
L4/5 IT CTX
L2/3 IT PPP
L2/3 IT CTX
L2/3 IT AI
Immune
Ependymal
CR
Car3
Astrocytes
-6
-4
-2
0
2
Log2FC (KO / WT)
0.00
0.05
0.10
0.15
0.20
Cell Type Proportion
Significance
FDR < 0.05 &
abs(Log2FD) > 0.58
n.s.
Genotype
KO
WT

**Fig. S30. Pgbd5 expression does not affect neuronal population proportions.** **A**, Uniform Manifold approximation and projection (UMAP) plot of single nuclei RNA seq from motor cortices of *Pgbd5*$^{wt/wt}$ and *Pgbd5*$^{-/-}$ littermates, samples (N = 18,107 WT, N = 14,359 KO) highlighting cells from each cluster containing >200 cells. Each cluster and their correspondent neuronal identity are colored blue. **B**, Neuronal proportions in *Pgbd5*$^{wt/wt}$ and *Pgbd5*$^{-/-}$ littermates (excluding "Striatal Neurons", "Other Neurons", and "No Consensus" cells). On the left: for each cell population the dot corresponds to the observed difference between proportions in the different genotypes, marked with confidence intervals based on the permutation test (n = 10,000). Significant comparisons (FDR < 0.05 & mean |$Log_2$ fold difference| > 0.58) were colored red showing that only Somatostatin Chodl (SSt Chodl) and L2/3 intratelencefalic para/post/presubiculum (L2/3 IT PPP) neuronal populations have a significant difference in proportion between wildtype and knock-out mice. On the right: Mean proportions for each cell type across replicates within each genotype (*Pgbd5*$^{wt/wt}$ (red) and *Pgbd5*$^{-/-}$(grey)). Error bars correspond to the standard error mean.

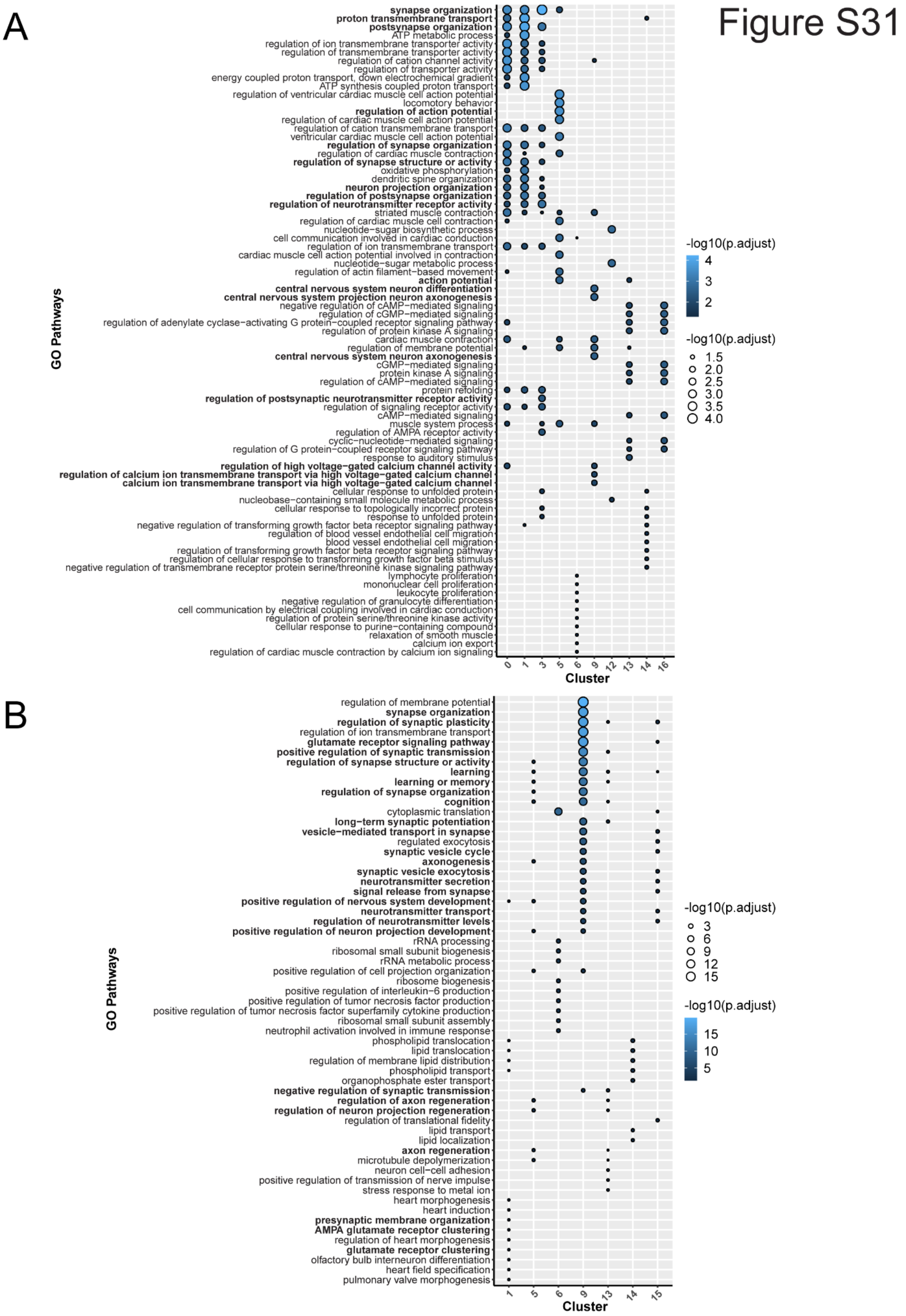
Figure S31
A
synapse organization
proton transmembrane transport
postsynapse organization
ATP metabolic process
regulation of ion transmembrane transporter activity
regulation of transmembrane transporter activity
regulation of cation channel activity
regulation of transporter activity
energy coupled proton transport, down electrochemical gradient
ATP synthesis coupled proton transport
regulation of ventricular cardiac muscle cell action potential
locomotory behavior
regulation of action potential
regulation of cardiac muscle cell action potential
regulation of cation transmembrane transport
ventricular cardiac muscle cell action potential
regulation of synapse organization
regulation of cardiac muscle contraction
regulation of synapse structure or activity
oxidative phosphorylation
dendritic spine organization
neuron projection organization
regulation of postsynapse organization
regulation of neurotransmitter receptor activity
striated muscle contraction
regulation of cardiac muscle cell contraction
nucleotide−sugar biosynthetic process
cell communication involved in cardiac conduction
regulation of ion transmembrane transport
cardiac muscle cell action potential involved in contraction
nucleotide−sugar metabolic process
regulation of actin filament−based movement
action potential
central nervous system neuron differentiation
central nervous system projection neuron axonogenesis
negative regulation of cAMP−mediated signaling
regulation of cGMP−mediated signaling
regulation of adenylate cyclase−activating G protein−coupled receptor signaling pathway
regulation of protein kinase A signaling
cardiac muscle contraction
regulation of membrane potential
central nervous system neuron axonogenesis
cGMP−mediated signaling
protein kinase A signaling
regulation of cAMP−mediated signaling
protein refolding
regulation of postsynaptic neurotransmitter receptor activity
regulation of signaling receptor activity
cAMP−mediated signaling
muscle system process
regulation of AMPA receptor activity
cyclic−nucleotide−mediated signaling
regulation of G protein−coupled receptor signaling pathway
response to auditory stimulus
regulation of high voltage−gated calcium channel activity
regulation of calcium ion transmembrane transport via high voltage−gated calcium channel
calcium ion transmembrane transport via high voltage−gated calcium channel
cellular response to unfolded protein
nucleobase−containing small molecule metabolic process
cellular response to topologically incorrect protein
response to unfolded protein
negative regulation of transforming growth factor beta receptor signaling pathway
regulation of blood vessel endothelial cell migration
blood vessel endothelial cell migration
regulation of transforming growth factor beta receptor signaling pathway
regulation of cellular response to transforming growth factor beta stimulus
negative regulation of transmembrane receptor protein serine/threonine kinase signaling pathway
lymphocyte proliferation
mononuclear cell proliferation
leukocyte proliferation
negative regulation of granulocyte differentiation
cell communication by electrical coupling involved in cardiac conduction
regulation of protein serine/threonine kinase activity
cellular response to purine−containing compound
relaxation of smooth muscle
calcium ion export
regulation of cardiac muscle contraction by calcium ion signaling
GO Pathways
0 1 3 5 6 9 12 13 14 16
Cluster
-log10(p.adjust)
4
3
2
-log10(p.adjust)
1.5
2.0
2.5
3.0
3.5
4.0
B
regulation of membrane potential
synapse organization
regulation of synaptic plasticity
regulation of ion transmembrane transport
glutamate receptor signaling pathway
positive regulation of synaptic transmission
regulation of synapse structure or activity
learning
learning or memory
regulation of synapse organization
cognition
cytoplasmic translation
long−term synaptic potentiation
vesicle−mediated transport in synapse
regulated exocytosis
synaptic vesicle cycle
axonogenesis
synaptic vesicle exocytosis
neurotransmitter secretion
signal release from synapse
positive regulation of nervous system development
neurotransmitter transport
regulation of neurotransmitter levels
positive regulation of neuron projection development
rRNA processing
ribosomal small subunit biogenesis
rRNA metabolic process
positive regulation of cell projection organization
ribosome biogenesis
positive regulation of interleukin−6 production
positive regulation of tumor necrosis factor production
positive regulation of tumor necrosis factor superfamily cytokine production
ribosomal small subunit assembly
neutrophil activation involved in immune response
phospholipid translocation
lipid translocation
regulation of membrane lipid distribution
phospholipid transport
organophosphate ester transport
negative regulation of synaptic transmission
regulation of axon regeneration
regulation of neuron projection regeneration
regulation of translational fidelity
lipid transport
lipid localization
axon regeneration
microtubule depolymerization
neuron cell−cell adhesion
positive regulation of transmission of nerve impulse
stress response to metal ion
heart morphogenesis
heart induction
presynaptic membrane organization
AMPA glutamate receptor clustering
regulation of heart morphogenesis
glutamate receptor clustering
olfactory bulb interneuron differentiation
heart field specification
pulmonary valve morphogenesis
GO Pathways
1 5 6 9 13 14 15
Cluster
-log10(p.adjust)
3
6
9
12
15
-log10(p.adjust)
15
10
5

**Fig. S31. Pgbd5 affects synapse formation as well as regulation of action potential in both interneuron and glutamatergic populations. A-B,** Top GO pathways corresponding to the DEGs identified in each cluster within the snRNA-seq joint sample (*Pgbd5*$^{wt/wt}$ and *Pgbd5*$^{-/-}$) object. Separate dot plots depict the GO pathways enriched in DEGs upregulated in *Pgbd5*$^{wt/wt}$ (**A**), and *Pgbd5*$^{-/-}$ (**B**) cells from each cluster. GO pathways appearing in the top 10 significant pathways for at least 1 cluster are plotted. Only clusters with greater than 200 cells, corresponding to cell populations of cortical origin (excluding clusters of 'Striatal Neurons' and 'Other Neurons'), and with at least one significant GO pathway (p.adj < 0.05) are shown in each plot. Both bubble size and color gradient correspond to log10 of p adjusted value.

Figure S32

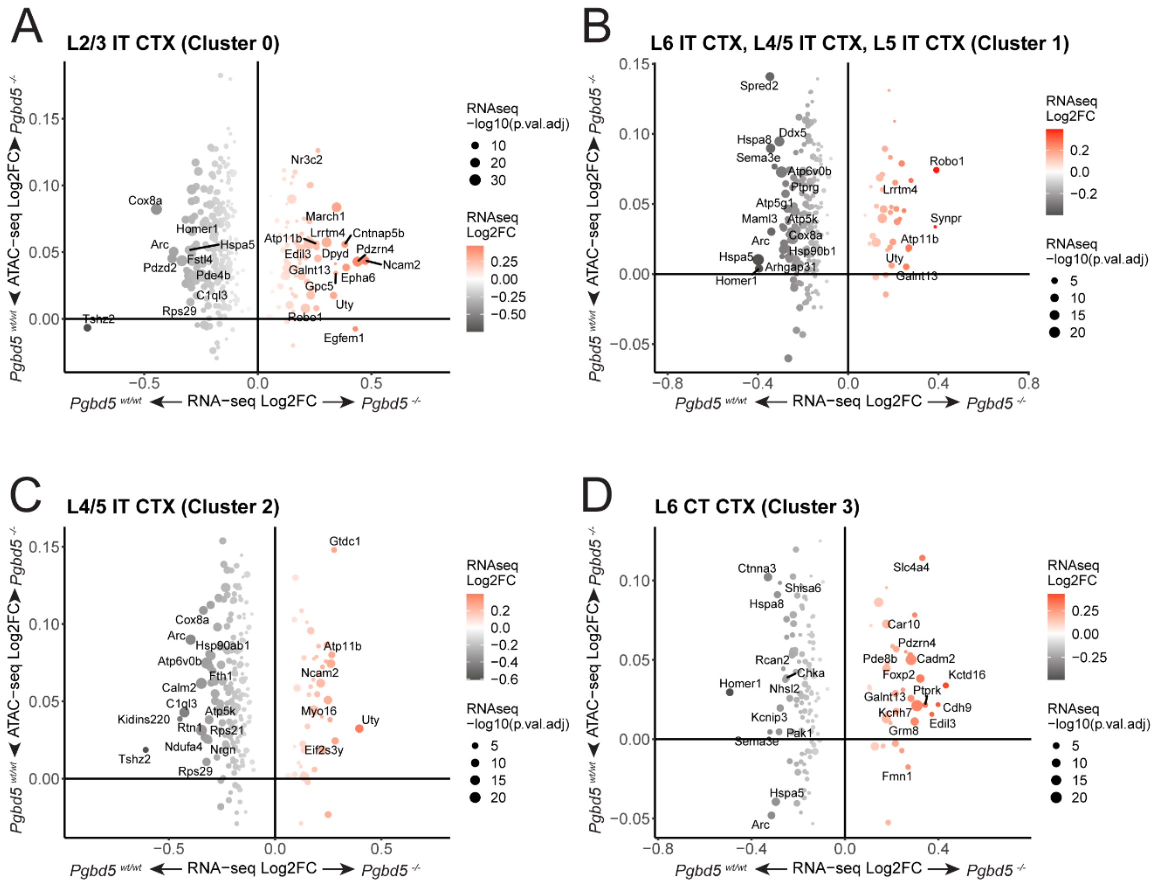


**Fig. S32. Some of the Intratelencephalic glutamatergic population significant transcriptional changes do not correlate with its accessibility profile. A-D,** 2D bubble plot depicting changes in gene expression (RNA-seq Log2FC) correlated with changes in chromatin accessibility (ATAC-seq Log2FC) at the corresponding gene promoter region (+/- 2.5kb from TSS) between *Pgbd5*$^{wt/wt}$ (in black) and *Pgbd5*$^{-/-}$ (red) littermates in Cluster 0 containing L2/3 IT CTX (**A**), cluster 1 containing L6, L4/5 and L5 IT CTX (**B**), cluster 2 containing L4/5 IT CTX (**C**) and cluster 3 containing L6 CT CTX (**D**). Only genes with significant changes in expression (P.Adj < 0.05) are plotted. In all these neuronal populations there are significant transcriptional changes that do not always correlate with the DNA accessibility measured by ATACseq.

**Supplemental Data Files**

**Data S1 (separate file)**

Clinical information for human patients.

**Data S2 (separate file)**

Pgbd5_KI_sequencing: contains a table listing the structural variants found in whole genome sequencing of *$Pgbd5^{ki}$* founder mice.

**Data S3 (separate file)**

Quantified_images: contain all the original files that were used to manually quantify the DNA damage of *$Pgbd5^{wt/wt}$*, *$Pgbd5^{-/-}$*, *$Pgbd5^{ki/ki}$*, *$Xrcc5^{-/-}$;$Pgbd5^{wt/wt}$* and *$Xrcc5^{-/-}$; $Pgbd5^{-/-}$* E14.5 embryos.

**Data S4 (separate file)**

WGS_data: contains files and information related to the whole genome sequencing of *$Pgbd5^{wt/wt}$* and *$Pgbd5^{-/-}$* mice, including metadata of SRA deposit of the whole genome sequencing data, the generated vcf files used in the analysis as well as a table listing the recurrent rearrangements found in *$Pgbd5^{wt/wt}$* .
