## Supplementary material for "A transposase-derived gene required for human brain development": Data_S1

**PGBD5 Supplemental clinical summaries**

**Family 1**

**Patient 1**: The proband was born to consanguineous (2nd cousin) Moroccan parents after an unremarkable term pregnancy. He presented with severe cognitive deficiency, no language, and has ASD. There was significant spastic tetraparesis. Tendon reflexes are increased and Babinski sign is present. Starting at 8 months and spanning the 3 first years of life, he presented multiple febrile and non febrile generalized tonic-clonic status epilepticus. He is seizure free on valproate.

Video analysis of movements (MCK) identified severe axial hypotonia and ataxic features of dyssynergic steps and dysmetria in fine motor tasks.

Height, weight, and orbital frontal circumference (OFC) were normal. Dysmorphic features present and reported in **Table S2**.

MRI was performed at 4 and 10 years of age. Normal signal, myelination, sulcation, and formation of structures. No lesions. Narrowing of nasopharyngeal airway. At 4 years, the anteroposterior diameter of the corpus callosum is 56 mm, within the 3rd percentile for age and gender. The measurements of each callosal segment between the 3rd percentile and median for age. A follow up MRI at 10 years revealed cerebellar atrophy (H-V 36.6 mm, APD-V 19.1 mm, APD-P 20.2 mm, APD-MP 10.5 mm), characterized by cerebellar/vermian folia widening, more evident in the upper-mid lobes. Cerebellar volume analysis (AP) confirmed significant reduction for age (-2.23 standard deviations (sd)).

**Patient 2:** The sister resembles the older brother with a slightly milder phenotype. She presented with febrile and non-febrile generalized tonic-clonic status epilepticus. She is currently seizure free on valproate. She also has a history of spasticity, increased tendon reflexes with Babinski sign, and language delay.

Height was -2.0 sd, weight was +2.0 sd, and OFC was normal. Dysmorphic features were present and are reported in **Table S2**.

Video analysis (MCK) revealed mild axial hypotonia, titubation while crawling, ataxic gait, and dysmetria in fine motor tasks.

MRI at 2 years demonstrated corpus callosum thinning with otherwise normal findings. 3D Cerebellar volume measurement (AP) were within normal parameters for age (-0.73 sd).

Whole exome sequencing revealed homozygous nonsense variant in PGBD5 (NM_001258311.2)c.49G>T p.(Glu17*), which was confirmed heterozygous in both parents and homozygous in the female sibling via Sanger sequencing.

**Family 2**

**Patient 1:** The index patient was the product of a consanguineous (2nd cousin) union, born to a 35 year-old mother at term without complications. He weighed 2350 grams at birth. He received phototherapy for indirect hyperbilirubinemia. At one month of age, he started to develop attacks of screaming followed by hyporesponsiveness diagnosed as epileptic seizures. Oxcarbazepine was initiated, followed by valproate and he became seizure-free by 10 months of age. Oxcarbazepine was later switched to clonazepam. He managed to hold his head at 5 months. He learned to sit independently after physiotherapy at age 3 years. Physical examination age at six years showed weight 19 kg (25%), height 118 cm (50-75%), head circumference 53 cm (50%). He had prominent ears and 4 cafe au lait spots (the largest measuring 1 x 1.5 cm). He had axial hypotonia with distal laxity and hyperextensibility of the fingers. He had spasticity of his lower extremities, increased deep tendon reflexes, positive Babinski reflex and clonus bilaterally. He had stereotypic movements of his upper extremities, did not make eye contact, and uttered no words.

Laboratory investigations, including complete blood count with differential, chemistry panel and blood gases were normal. Ammonia, amino acid profile and biotinidase activity were normal. Succinyl-5-aminoimidazole-4-carboxamide-1-ribose-5’-phosphate (SAICAR) was normal. MECP2 DNA sequence analysis and chromosomal microarray were unremarkable.

Urine organic acid analysis revealed no findings consistent with any known disorder. Lactate, pyruvate, guanidinoacetatem very long chain fatty acids and transferrin isoelectric focusing was all unremarkable. An electroencephalogram (EEG) performed at age 6 demonstrated epileptiform activity independently arising from both frontal lobes with spread to the right temporal lobe, activated during sleep. Flash visual evoked potentials revealed bilateral prolonged P1 latencies while flash electroretinogram exhibited prolonged latency on the right.

Brain stem auditory evoked potentials (75 dB) was normal on the right, but with no response on the left.

MRI at 14 years and 11 months revealed marked vermian-cerebellar atrophy with H-V 36.6 mm and APD-V 19.5 mm, both below the 3^rd^ percentile for age. The corpus callosum was also notably thin, with the genu below the 3^rd^ percentile (7.1mm), and the remaining measurements between the 3^rd^ percentile and median for age (body 4.1 mm, isthmus 3.1 mm, and splenium 8.8 mm).

**Patient 2**: The second affected boy was born at term via caesarian section, weighing 3350 grams. Despite some initial cyanosis, he did not require any special care in the nursery. He began to have generalized tonic-clonic seizures at the age of 6 months but achieved seizure-freedom with carbamazepine. He never learned to walk but could sit independently on his knees. He achieved only a few expressive words and experienced dysphagia. On examination at age 14, weight was 43 kg (below 3^rd^ percentile). He had intellectual disability, strabismus, ophthalmoplegia, and generalized dystonia prominent in both upper extremities and his cervical region. There was also evidence of spasticity and contractures, particularly in the lower extremities, with exaggerated deep tendon reflexes. He was treated with biperidene hydrochloride with limited benefit.

Laboratory investigations including complete blood count, chemistry panel, ammonia, amino acid profile and biotinidase activity were normal. SAICAR was negative. Lactic acid was unremarkable. Creatinine kinase, lysosomal enzyme analysis, urine organic acid analysis, ceruloplasmin, very long chain fatty acid analysis, MECP2 gene sequencing and chromosomal microarray were all unrevealing.

Whole exome sequencing revealed a single bp deletion causing a frameshift and early stop in PGBD5 (NM_001258311.2) c. 509del, p.(Phe170Serfs*5) which was confirmed heterozygous in both parents and an unaffected sister and homozygous in both affected male siblings via Sanger sequencing.

**Family 3**

**Patient 1:** She was born in a consanguineous union after an uneventful pregnancy and delivery with normal early life. At 3 months, she developed fits in form of up rolling of eyes and limb jerks. Seizures remained for few seconds and resolved spontaneously. At age 6 months, there was worsening of duration and frequency of seizures which became tonic clonic. Her EEG was suggestive of diffuse epileptic foci and she was started on Carbamazepine along with valproic acid. Seizures partially controlled initially and became settled with maximum doses of both antiepileptic medications.

She has cognitive, motor, and speech delays. She can sit, stand with support, but not walk, and has issues with fine motor control and coordination. She can only speak 2-3 words with no sentences. Neurological exam found signs of upper motor neurons lesion with hypertonia, hyperreflexia, and intermittent dystonia. Video review (MCK) identified titubation while crawling and confirmed difficulties with fine motor control.

 	Weight was 18 kg, height was 105 cm at 7 years. Plasma and urine analysis was inconclusive. Dysmorphic features were noted (**Table S2**).

**Patient 2**: She was born in a consanguineous union with one older affected sibling and one normal sibling after an uneventful pregnancy and delivery with normal early life. At 5 months she developed tonic clonic seizures, which were generalized, lasted 2- 3 minutes, and were associated with post ictal vomiting and drowsiness. EEG found diffuse abnormal discharges consistent with epilepsy. She was started on anti-epileptics Valproic acid and carbamezipine plus fat-soluble vitamins which has controlled the seizures.

She has progressive regression of developmental milestones along with failure to achieve age-appropriate milestones for motor, cognitive, and speech. Speech is limited to 2-3 words with no sentences or meaningful speech. She can sit and crawl but is unable to stand or walk. She has issues with fine motor control and poor coordination. Neurological exam revealed signs of upper motor neurons lesion with hypertonia and brisk reflexes.

At 4 years, weight was 15 kg and height was 99 cm. Blood plasma and urine analysis was inconclusive. Dysmorphic features were noted (**Table S2**).

Whole exome sequencing with Centogene revealed a nonsense variant in PGBD5 c.138C>A (p.Tyr46*).

**Family 4**

**Patient 1**: He was the first child of healthy consanguineous parents of Tunisian descent. He was born at term following an unremarkable pregnancy via cesarean delivery. No perinatal complications were reported and birth measurements were within the normal range. He was referred to the department of pediatric neurology at 6 years because of developmental delay, epilepsy and ataxia. He had positive family history of developmental delay and epilepsy in three cousins. Global developmental delay was noted since the first months of life. He had head control at 9 months of age, sat independently at 15 months, started babbling at 2 years, stand with support at 3 and walked aided at 4. By the age of 9 months, the child experienced his first seizure as febrile generalized tonic episodes. Valproic acid was started with satisfactory response, having had only one further seizure reported so far.

Neurological evaluation revealed spastic ataxic gait, truncal hypotonia, intention tremor dysmetria, brisk reflexes, bilateral Babinski sign. He could speak with 2-worded sentences and had severe intellectual disability. At repeat neurological examination, he developed contracture in lower limbs and lost the ability to walk by the time he reached 11 years.

On physical examination at 8 years, his weight, height and occipito-frontal circumference (OFC) were within normal limits without any striking dysmorphic features. Electromyography and alpha-fetoprotein were normal. Brain magnetic resonance imaging (MRI) was normal at the first year of life but demonstrated cerebellar atrophy at the age of 8.

**Patient 2**: The younger sister of was referred to the same department at 3 years because of similar complaints. She was born at term following an unremarkable pregnancy. No perinatal complications were reported and birth measurements were within the normal range. By the age of 6 months, she experienced afebrile generalized tonic-clonic seizures that were treated with valproic acid. After this event, she remained seizure free. The developmental milestones were remarkably delayed. She had head control and sat independently at one year of age and said her first words at 3. Her mother observed sialorrhea at 4 years.

Physical examination at 4 years and half revealed microcephaly with an OFC of 47 cm (-2.3 SD) with no dysmorphic facial features, thoracic hyperkyphosis and bilateral pes valgus. On neurological examination, she had good social contact, could walk supported with a spastic ataxic gait, she had pyramidal tract signs: generalized spasticity, brisk reflexes, bilateral Babinski sign and cerebellar syndrome: truncal hypotonia, dysmetria and horizontal nystagmus.

At her last follow-up at the age of 8, she could speak individual words, developed contracture in lower limbs and lost the ability to walk.

Complete blood count, serum biochemistry, complete metabolic panel (lactate and pyruvate in blood and in cerebrospinal fluid, ammonemia, plasma amino acids, urine organic acids, serum creatine and uric acid), transferrin isoelectric focusing, electroencephalography, ophthalmologic examination and audiometric evaluation were all normal in both siblings.

Brain MRI at age 6 showed diffuse cortical and subcortical atrophy, thin corpus callosum and cerebellar atrophy (**Figure 1**).

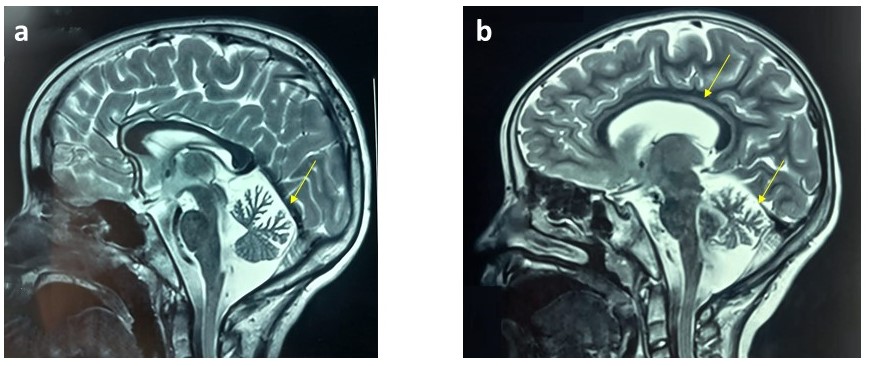

Figure 1 : Brain MRI findings:

a) T2 weighted image of patient 1 at the age of 8 years, demonstrates cerebellar atrophy

b) T2 weighted image of patient 2 at the age of 6 years shows diffuse cortical and subcortical atrophy, thin corpus callosum and cerebellar atrophy

Whole exome sequencing identified a deletion in PGBD5 resulting in a frameshift and early truncation (NM_001258311.2) c.1442_1490del, p.(Ile481ThrfsTer2). Sanger sequencing confirmed this variant was heterozygous in the unaffected mother and father and present in both affected siblings.

**Family 5**

**Patient 1:** He was born to parents of Syrian descent as a product of a consanguineous union. His birth was notable for fetal distress resulting in a caesarion delivery.

At 4 months, he presented with hypotonia, decreased reflexes, seizures, cognitive delay. He had several EEGs; at 4+5 months Sleepstage II; no epileptiform activity, by 6 months there were spikes left occipital and a small amount of spikes left and right parietal in a second test. 8 months there was wakefulness with no epileptic activity. He has been seizure free after treatment with Zonisamide and levetiracetam.

He currently has axial hypotonia, cognitive delay and no speech. He cannot stand or walk and exhibits mild titubation of his head.

Height is very short at 98cm (-5.41 sd) with normal weight for his height at 15kg (-0.20 sd weight for length) and OFC within normal parameters at 50cm (-1.26sd). Dysmorphic features are present (**Table S2**).

Brain MRI was conducted at 4 months and 3 years of age. Sizes of structures and myelination within normal parameters for age. No lesions. Symmetric widening of subarachnoid spaces and ventricles thought to be within physiologic parameters at 4 months, not apparent at 3 years. Cerebellar size appeared within normal parameters at both ages. Corpus callosum appeared normal at 4 months with thinning apparent at 3 years with the splenium below the 3^rd^ percentile for age (6.7 mm), with the other parameters between 3^rd^ percentile and median (genu 6.6mm, isthmus 4.0 mm, and splenium 3.7 mm). Cerebellar volume analysis (AP) indicates within normal range for age (-0.96 sd).

**Patient 2:** Her prenatal and delivery history was unremarkable. At 4 months, seizures were noted. Multiple EEG tests were conducted; at 4 months there was multifocal epileptic activity and no epileptic activity in a second test. At 5 months, there was a small amount of spikes left and right parietal. Despite ongoing treatment with Zonisamide, levetiracetam, and Carbamazepine, she has ongoing focal seizures.

She has axial hypotonia, cognitive delay and no speech. She cannot stand or walk. A VEP at 3 years found normal velocity and amplitude.

Brain MRI was conducted at 3 and 21 months. Sizes of structures and myelination within normal parameters for age. No lesions. Corpus callosum and cerebellar measurements were normal for that period. Cerebellar volume analysis (AP) within normal parameters for age (-1.45 sd). Dysmorphic features were present (**Table S2**).

Whole exome sequencing revealed a deletion variant in PGBD5 resulting in a frameshift and early truncation (NM_001258311.2) c.214del, p.Ala72Profs*91.
